# Genetic Manipulation of Mammalian Cells in Microphysiological Hydrogels

**DOI:** 10.1101/2025.03.21.644519

**Authors:** Anna C. Jäkel, Dong-Jiunn Jeffery Truong, Friedrich C. Simmel

## Abstract

Engineering functional 3D tissue constructs is essential for developing advanced organ-like systems, with applications ranging from fundamental biological research to drug testing. The generation of complex multicellular structures requires the integration of external geometric and mechanical cues with the ability to activate genetic programs that regulate and stimulate cellular self-organization. Here, we demonstrate that gelatin methacryloyl (GelMA) hydrogels serve as effective matrices for 3D cell culture, supporting both in situ genetic manipulation and cell growth. HEK293T cells embedded in GelMA remained viable and proliferated over 16 days, forming clusters within the matrix. We achieved efficient gene delivery in the 3D hydrogel environment using both plasmid DNA and mRNA as gene vectors. Furthermore, we applied in situ prime editing to induce permanent genetic modifications in embedded cells. To achieve spatially confined gene expression, we introduced gel-embedded channels that allowed localized stimulation via doxycycline perfusion through a Tet-On system. Our findings establish GelMA hydrogel matrices as a versatile platform for generating spheroidal cell cultures while enabling precise genetic control and spatially resolved cellular manipulation through diffusible cues.

## 1 Introduction

There are two general approaches to the realization of synthetic tissues and organ-like systems from living cells. One approach focuses on the generation of external scaffold structures via bio-fabrication techniques such as soft lithography and 3D bioprinting. The complementary approach relies on the natural self-organization properties of cells, where cells autonomously form intricate structures through processes such as differentiation, growth, and intercellular communication (Figure 1a). Ideally, both approaches would be combined - using geometric and mechanical boundary conditions provided by the scaffold, while simultaneously allowing cellular self-organization processes to unfold. These self-organization processes are often governed by pattern-forming genetic programs, which need to be appropriately regulated to guide tissue formation.

**Figure 1:**
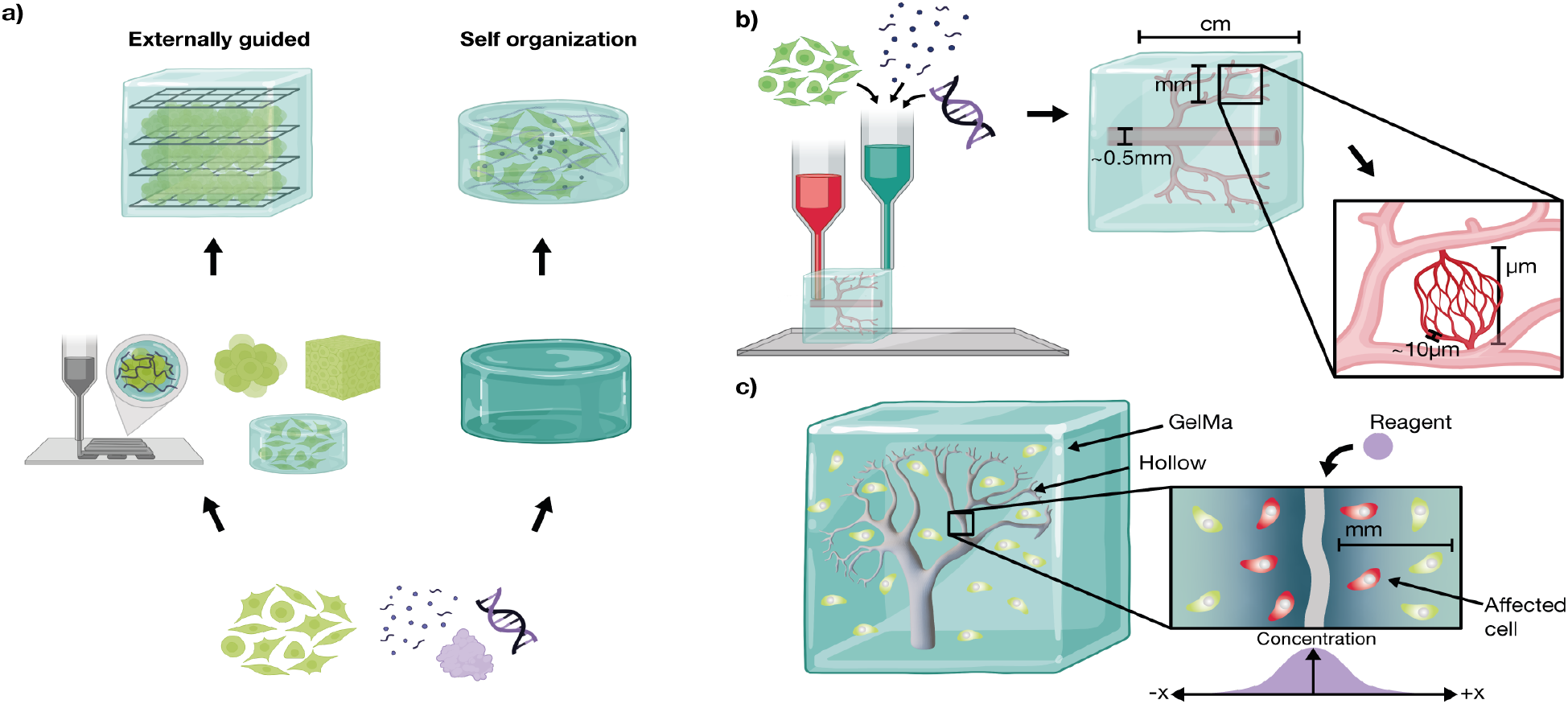
Approaches and Challenges in Tissue Fabrication. a) Tissue engineering can follow two primary strategies: externally guided construction and self-organization. The externally guided approach employs techniques such as 3D bioprinting to prestructure tissues using cell sheets, organoids, cell aggregates, or hydrogels as scaffolds. In contrast, self-organization relies on the intrinsic ability of cells to grow and establish structures within a hydrogel matrix, driven by cellular interactions and biochemical cues. b) To integrate both approaches, 3D bioprinting can provide a macroscopic framework at the cm to mm scale by embedding cells, DNA, proteins, and vascular-like structures within a matrix. At a finer resolution, self-organization mechanisms guide microscale tissue formation, leveraging the natural capacity of cells to refine structural details. This combination enables precise control over tissue architecture while preserving the ability of cells to develop functional microenvironments. c) Vascular-like structures (here: kidney vasculature from (*11*)) are essential for overcoming diffusion limitations in engineered tissues. Molecular transport in hydrogels is constrained by slow diffusion, which affects nutrient and signal delivery in mm to cm-sized constructs. To study localized stimulation in such environments, we employ a hydrogel block with embedded channels. Reagents perfused through these channels selectively affect cells in proximity, enabling their supply with nutrients and biochemical stimulation.

A major challenge in the creation of thick, tissue-like constructs is the limited effectiveness of passive diffusion over larger length scales. This limitation impairs nutrient supply, slows the removal of waste products, and hampers genetic control over cellular self-organization, including the exchange of intercellular signaling molecules. In natural tissues, this challenge is overcome by the integration of vascular systems, which span from microvasculature (with channel diameters ranging from approximately 5 to 100 *µ*m) to macrovasculature (with diameters up to several cm) (*1*). Researchers have already explored various strategies to generate synthetic vasculature for tissue engineering. On the one hand, cells themselves can be used to autonomously form vascular structures within engineered tissues (Figure 1b).

For instance, Griffith et al. demonstrated that human umbilical vein endothelial cells (HUVECs) can form vascular networks by self-organization in hydrogel matrices (*2*). In contrast, Paulsen et al. proposed 3D printing of vascular networks to guide structured tissue formation (*3*). This concept was subsequently implemented by multiple groups, including Kolesky et al., who used Pluronic as a fugitive ink to create perfusable channels within hydrogels (*4*), and Pimentel et al., who utilized polyvinyl alcohol (PVA) as a sacrificial material (*5*).

In the present work, we demonstrate that cells can be cultured within Gelatin methacryloyl (GelMA) hydrogels for extended periods, while enabling in situ genetic manipulation. Both nutrient supply and the delivery of genetic constructs can be facilitated via synthetic channel structures, which ultimately could allow to combine externally guided tissue formation with genetically programmed self-organization, enabled by synthetic biology tools (Figure 1c).

GelMA is a versatile hydrogel scaffold widely used in tissue engineering due to its biocompatibility, bioactivity, and tunable mechanical properties. Its arginine-glycine-aspartate (RGD) motifs promote cell adhesion and spreading, while its rapid photo-crosslinking allows precise structural control at physiological temperatures (*6*). By adjusting concentration and crosslinking density, GelMA can be tailored for applications ranging from 3D tissue constructs to bioreactor systems. Concentrations between 10–15% have been shown to balance structural integrity with cellular ingrowth and viability (*7*, *8*). GelMa has a pore size of order ≈100 µm (*9*). The diffusion coefficient *D* for small molecules with typical sizes of 1 nm in 10% GelMa has been found to be on the order of 10^*−*6^ cm^2^ s^*−*1^ (*10*), which is reduced roughly tenfold with respect to free diffusion. This results in diffusion times *t ∼ L*^2^*D*^*−*1^ exceeding *t* = 100 h over distances of *L* = 1 cm, rendering passive diffusion impractical for nutrient delivery in thick constructs.

Rather than using typical tissue engineering cell lines, we focused mainly on HEK293T cells due to their well-established genetic manipulation methods. HEK293T cells are widely used for synthetic gene circuit testing, protein expression, and genome editing (*12*, *13*), making them an ideal starting point before transitioning to primary or stem cell-based systems. Moreover, they can be co-cultured with fibroblasts, endothelial cells, or MSCs to study heterotypic cell-cell interactions, and they function as bioactive factories, secreting growth factors, cytokines, or signaling molecules that influence neighboring cells.

We use our model system to demonstrate in situ gene induction, gene delivery as well as genome editing within 3D hydrogel matrices. Gene delivery within 3D cell cultures is an emerging tool in tissue engineering and gene therapy (*14*). While transfection in 2D is well established, adapting these strategies to 3D hydrogels poses additional challenges, including reagent penetration, diffusion kinetics, and vector stability. Previous work has tested various transfection reagents for hydrogel cultures (*15*, *16*), showing that mRNA transfection can be more efficient than plasmid DNA delivery in certain 3D models (*17*). Beyond transient gene expression, permanent genomic modifications allow long-term functional control of engineered tissues. As a step towards this goal, we demonstrate CRISPR/Cas-based prime editing (*18*) within GelMA hydrogels, with its necessary components delivered either as plasmid DNA or, alternatively, RNA.

## 2 Results

### Growth Dynamics of Cells Seeded in Hydrogel

Cell growth and viability within 3D gel matrices are strongly influenced by structural properties of the matrix such as pore size, porosity, and interconnectivity, which impact nutrient transport, waste removal, and overall scaffold stability. Smaller pores enhance cell attachment and intra-cellular signaling, while larger pores facilitate gas diffusion and vascularization, necessitating an optimal balance between porosity and mechanical integrity (*19*). As we envisioned the inclusion of vascular structures in our hydrogel constructs we chose a GelMa concentration of 15%, as this has been proven to show good shape fidelity (*20*).

To evaluate cellular behavior over an extended period of time, HEK293T cells were embedded in 15% GelMa containing 0.25% Lithium-Phenyl-2,4,6-Trimethylbenzoylphosphinate (LAP) for crosslinking and cultured for 16 days, with medium exchange every 3–4 days. The rapid crosslinking of GelMa provided a stable 3D structure, enabling cell proliferation in a physiologically relevant environment. One day after seeding, cells were observed as singlets, doublets, and triplets, with cluster sizes reaching up to 44 µm in diameter (Figure 2a). Over 18 days, cells proliferated into spherical clusters, reaching diameters of approximately 500 µm.

**Figure 2:**
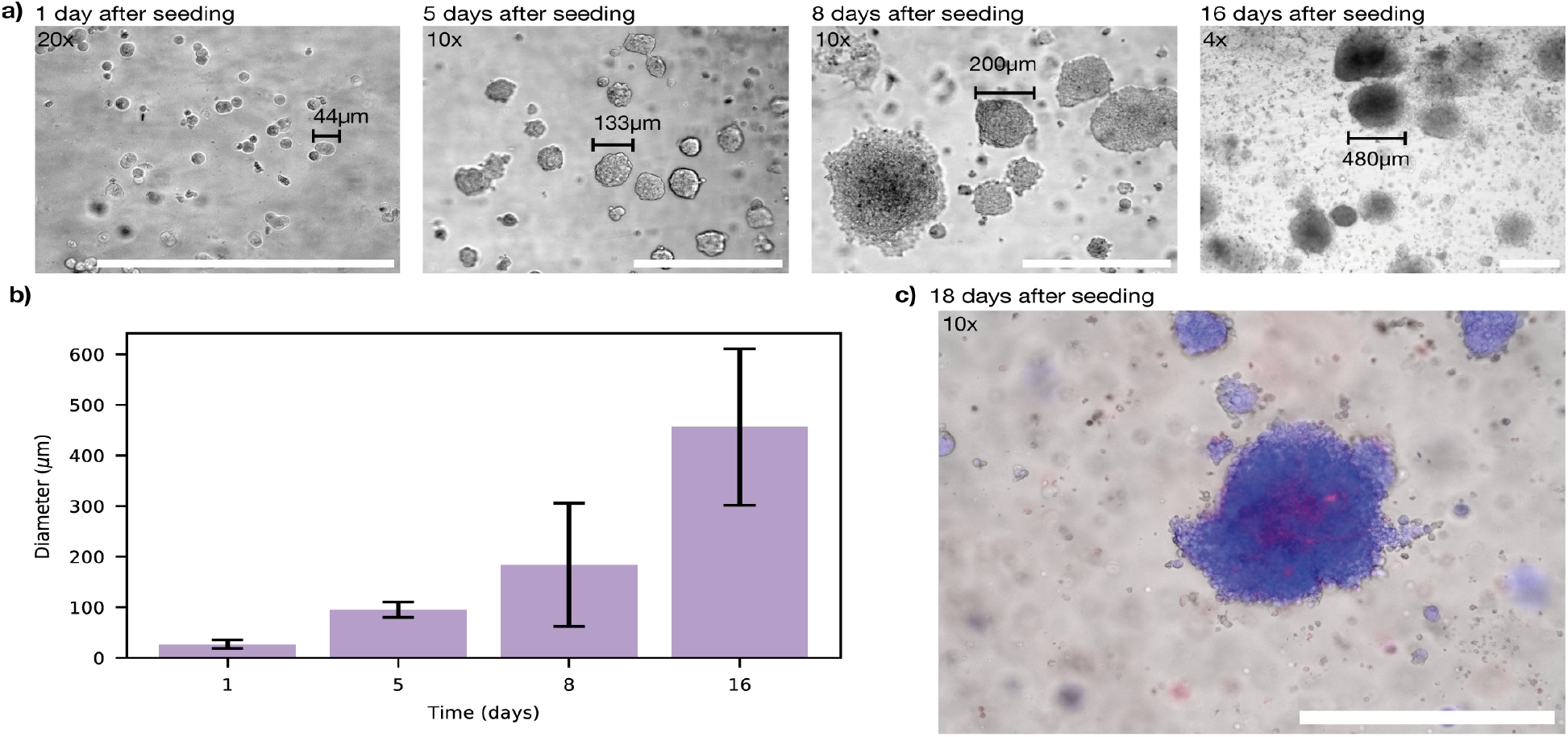
Cell growth in 15% homemade GelMa crosslinked with 0.25% LAP using 405 nm UV for 30 s. a) Proliferation of HEK293T cells embedded in GelMa over a 16-day culture period. b) Formation of multicellular clusters within the hydrogel, with an estimated doubling time of approximately two days. Error bars represent standard deviation, for each timepoint 10 clusters were measured. c) Live/dead staining using Hoechst (blue, live) and Propidium Iodide (red, dead) to assess cell viability. The majority of cells remain viable, with minimal cell death observed, primarily localized at the center of cell clusters. Scale bar: 500 µm for all images.

A quantitative analysis of the microscope images (Figure 2b) confirms consistent cell growth within GelMa indicating that the matrix provided adequate structural and biochemical support for proliferation. Assuming roughly spherical clusters, we can calculate the cluster volume 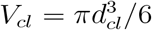 from the measured diameter *d*_*cl*_. *V*_*cl*_ is proportional to the number of cells in the cluster, i.e., *V*_*cl*_ = *N*_*c*_ *· V*_*c*_, where *V*_*c*_ is the cell volume. For constant growth the number of cells increases over time as 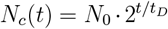. We can thus extract an estimate for the cell doubling time in the gel from our data, which results in *t*_*D*_ ≈ 2 days (Supplementary Figure S3). The final clusters contain *N*_*c*_ ≈ 5000 cells.

Live/dead staining performed on day 18 (Figure 2c) confirmed high cell viability throughout the culture period. Minimal cell death was observed, primarily localized at the cluster centers. This observation is consistent with previous studies that have reported a maximum cell cluster size of 500 µm before central necrosis occurred (*21*), likely due to diffusion limitation of nutrients.

To compare if the proliferation behavior into clusters in our homemade GelMa is similar for different cell types, we assessed the compatibility of our hydrogel with two other cell lines: NIH cells and hMSCs. The results, presented in the SI (Figure S9), demonstrate that both HEK293T and NIH cells proliferated within the hydrogel and exhibited cluster formation, whereas hMSCs distributed homogeneously throughout the matrix without forming clusters. For subsequent studies, HEK293T cells were selected as the primary model due to their well-established suitability for genetic modification.

### Induction of Gene Expression in Matrix-Embedded Cells

Inducible gene expression in cells embedded in hydrogel was assessed using the Tet-On 3G system (*22*, *23*). HEK293T cells were genetically modified to stably express the fluorescent protein mGreenLantern as a baseline reporter, while mScarlet-I expression could be induced with doxycycline via the Tet-On 3G system (Experimental Section). In these cells, both components of the Tet-On 3G system are integrated into a single vector (Figure 3a). mGreenLantern is expressed under a CAG promoter in the forward orientation to enhance contrast for improved visualization of the cells in hydrogel. The Tet-On 3G transactivator is driven by the constitutive human phosphoglycerate kinase 1 promoter (hsPGK1) in reverse orientation, alongside the gene of interest – mScarlet-I –, which is put under the control of the inducible TRE3GS (TRE 3G) promoter. Cells in adherent culture displayed homogeneous constitutive expression of mGreenLantern as well as induced expression of mScarlet-I 24 h after induction with 500ng/mL doxycycline, with fluorescence levels being roughly similar across the cell population (Figure 3a).

**Figure 3:**
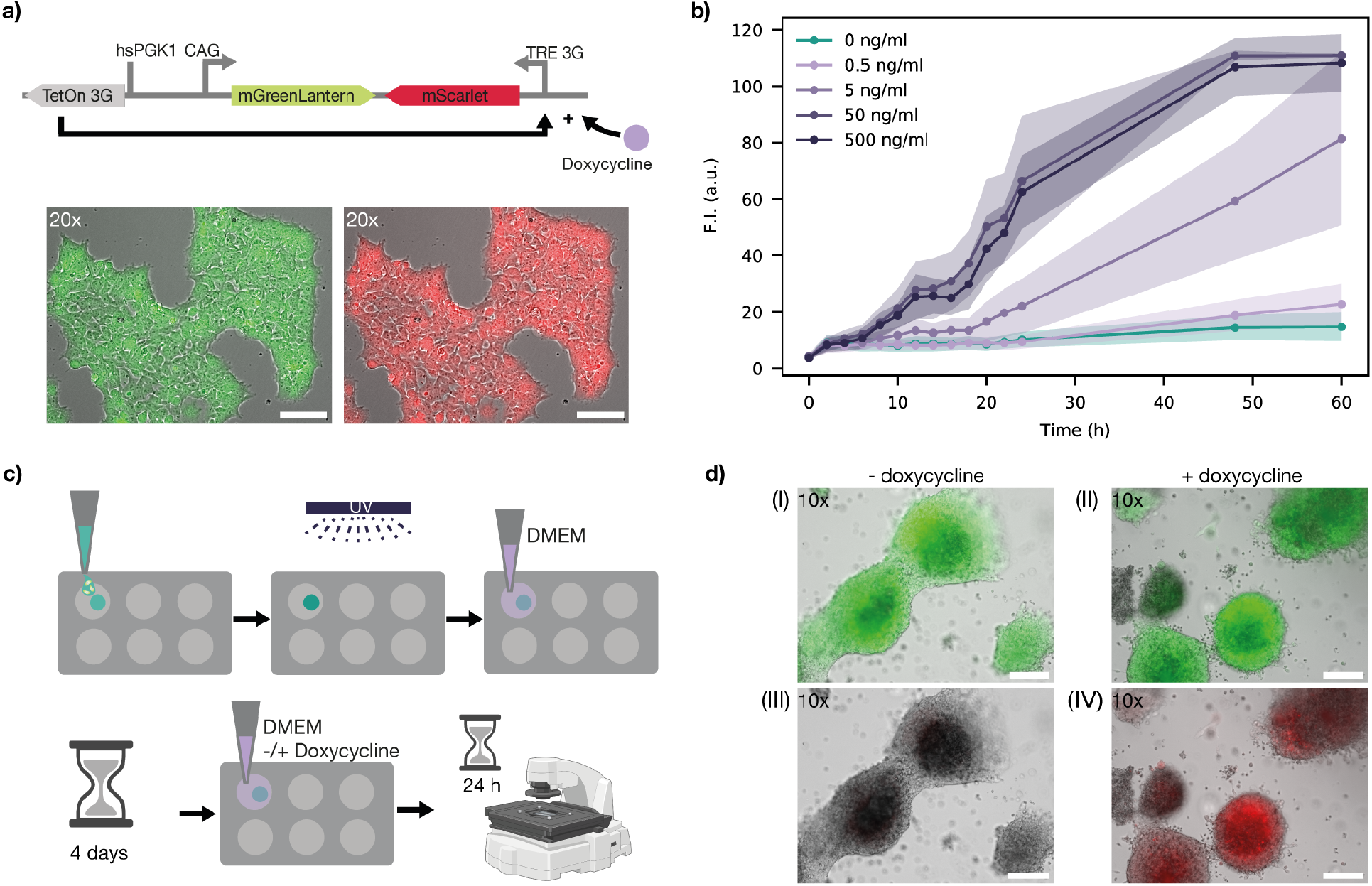
In situ gene induction. a) For gene induction experiments a genetically modified HEK293T cell line was used that constitutively expressed mGreenLantern from a CAG promoter, while mScarlet-I expression could be induced by the addition of doxycycline via the Tet-On system. The microscope panels show composites of brightfield and fluorescence channels for mGreen-Lantern expression (left) and mScarlet-I expression (right), 24 h after treatment with 500 ng/mL doxycycline. b) Dose-dependent induction of mScarlet-I in adherent HEK293T cells. A doxycycline concentration of 50 ng/mL is sufficient for full activation. c) HEK293T cells embedded in 15% GelMa at a density of 1 million cells per ml were cultured in well plates containning DMEM and induced with 500 ng/mL doxycycline after four days of incubation. Fluorescence intensities were recorded 24 h post-induction. d) Microscopy images of Gel-embedded cell clusters (overlays of brightfield and fluorescence channels). (I–II) Control and induced samples show mGreenLantern fluorescence. (III–IV) mScarlet-I fluorescence is only observed in the doxycycline induced sample. Scalebars: 200 *µ*m.

Although already 50ng/mL doxycycline were sufficient for full induction of adherent cells (Figure 3b), we chose 500ng/mL for all experiments in hydrogel-embeddded cells to ensure sufficient doxycycline penetration through the gel matrix for full induction. Cells were mixed into 15% GelMa at 10^6^ cells/mL, seeded into a 48-well plate, and crosslinked using UV light, as illustrated in Figure 3c. 300 µl of the cell gel mixture was applied to the well. As this has a surface of 1.1 cm^2^ the height of the hydrogel was ≈270 µm. UV exposure was limited to only a few seconds to keep its effect on the cells as low as possible. Culture medium was applied on top, and cells were grown until clusters reached a size of approximately 100 µm, typically after 4 days. Doxycycline was then added, and fluorescence was measured 24 h later (Figure 3d). Only doxycycline-exposed cells exhibited robust mScarlet-I fluorescence, demonstrating that in situ gene induction proceeds effectively within the gel. Induced cell clusters could be observed in the upper part (roughly 200 µm) of the gel which was exposed to 500 µl of DMEM.

### In situ Plasmid and mRNA Delivery

We next evaluated the influence of the GelMA matrix on transfection efficiency. While transfection protocols are well established in 2D cultures, adapting these strategies to 3D hydrogels presents additional challenges, including restricted diffusion and reagent penetration, reduced vector stability, and altered cell-matrix interactions that can affect vector uptake. Previous studies have explored transfection methods in 3D environments (*15*) and tested various reagents for hydrogel-based cultures (*16*). Notably, some studies have reported higher mRNA transfection efficiencies in specific 3D models (*17*).

To evaluate transfection performance in our system, we tested both pDNA and mRNA vectors, using jetOPTIMUS for plasmid DNA transfection and Lipofectamine 2000 for mRNA delivery. Both vectors must undergo several steps before generating a measurable expression signal (Figure 4a). First, the vectors must diffuse through the hydrogel matrix to reach the cells. The transfection complexes are expected to exhibit sizes in the range of 100 to 200 nm, with pDNA complexes likely being slightly larger than mRNA lipoplexes. As a result, minor differences in diffusivity can be expected.

**Figure 4:**
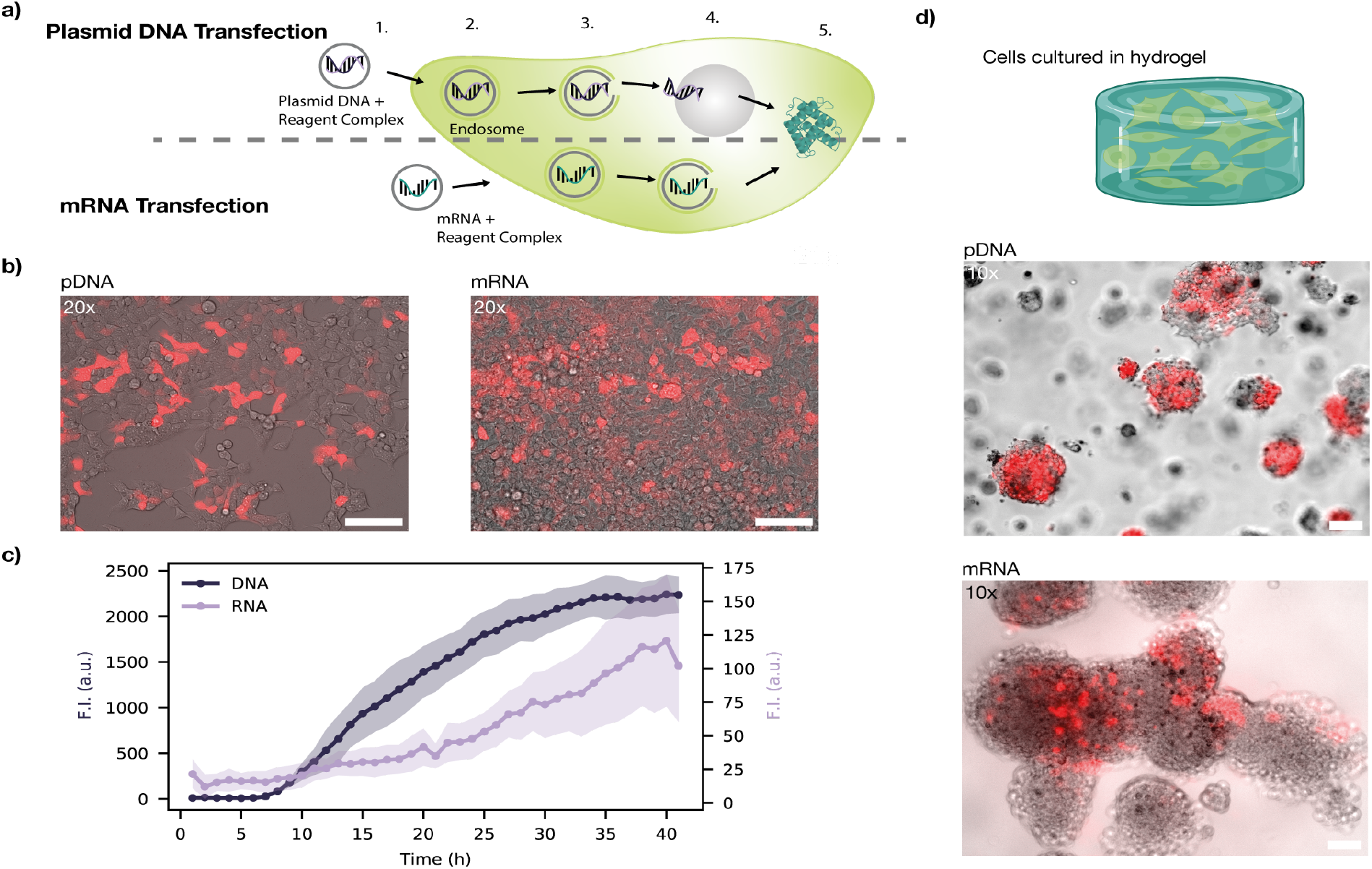
Transfection of HEK293T cells cultured in hydrogel with plasmid DNA (pDNA) versus mRNA. a) Schematic representation of the steps involved in pDNA vs. mRNA transfection: (1) Complexation of pDNA or mRNA with transfection agent, (2) cellular uptake via endocytosis, (3) endosomal escape, (4) for pDNA only: nuclear entry, transcription, and mRNA export (5) translation of mRNA into protein in the cytoplasm. b) Composite images of brightfield and mScarlet-I fluorescence in cells transfected with pDNA (left) and mRNA (right). Scale bar: 100 µm. c) Fluorescence intensity of mCherry expression following transfection with pDNA versus mRNA. For each curve fluorescence was determined in an ROI measured containing several cells (see SI Figure S1 for details.) The shaded area indicates the standard deviation, for each timepoint 3 ROIs were measured. d) Cells were mixed with 15% GelMa, pipetted into a well, and GelMa containing 0.25% LAP was crosslinked using 405 nm UV light for 30 s before the medium was applied on top. Composite images of brightfield and mScarlet-I fluorescence show HEK293T cells cultivated in GelMa and transfected with either pDNA or mRNA. Scale bar: 100 µm.

Further, plasmid DNA is more stable compared to mRNA, but has to be processed by the cell before the gene of interest is expressed. After cellular uptake, pDNA must enter the cell’s nucleus, where it undergoes transcription, followed by mRNA nuclear export before the protein can be translated. In contrast, mRNA transfection bypasses these steps and can be directly translated after entering the cytoplasm.

Using mCherry as a fluorescent reporter, we first tested transfection in adherent HEK293T cells (Figure 4b). Transfection with pDNA resulted in a signal increase after 6 h and levelled off after ≈ 30 h, resulting in an overall sigmoidal time-course of the fluorescence intensity. In contrast, mRNA transfection produced a weak signal with no apparent lag time and the fluorescence intensity followed a linear trend. mRNA transfection resulted in overall lower tenfold fluorescence intensities than when using pDNA as a vector (Figure 4c).

Next, we investigated the feasibility of in situ transfection of hydrogel-embedded cells. Transfection efficiency, defined as the percentage of cells that successfully take up and express the introduced genetic material, was used to evaluate the success of gene delivery. We found that pDNA vectors led to significantly higher transfection efficiencies compared to mRNA vectors (Figure 4d). In pDNA transfection, we consistently observed uniform expression within a cell cluster. In contrast, mRNA transfection primarily led to fluorescent protein expression in individual cells, but not in whole clusters.

### Precision Genome Editing in Hydrogel-Embedded Cells

To further advance the genetic manipulation of gel-embedded cells, we explored the possibility of in situ genome engineering via CRISPR-based prime editing (Figure 5a) (*18*). In prime-editing, a prime-editing guide RNA (pegRNA) containing a primer binding site (PBS) and a reverse transcriptase template binds with a Cas9-derived nickase (SpCas9n from Streptococcus pyogenes) fused to a reverse transcriptase (RT) domain. This complex binds to the target DNA at a sequence complementary to the pegRNA spacer, where SpCas9n cuts the unbound target strand. The PBS of the pegRNA binds the nicked strand and the sequence of the RT template is reverse transcribed onto the nicked DNA strand. A DNA repair mechanism then leads to the integration of the newly synthesized DNA at the target site.

**Figure 5:**
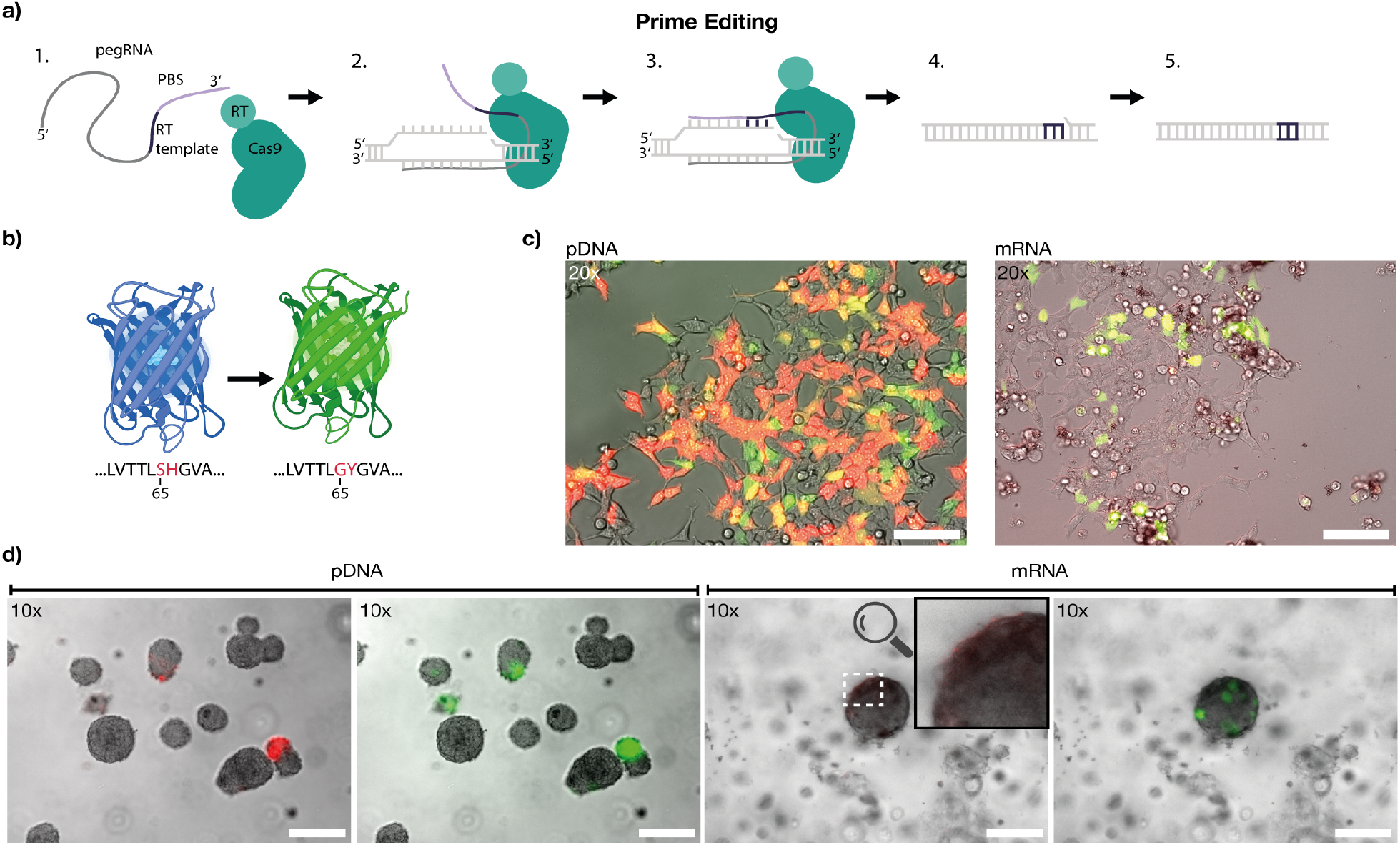
In situ prime editing. a) Prime editing mechanism: a prime editing guide RNA (pegRNA) contains the sequence for the desired edit along with a primer binding site (PBS), while the Cas9 protein is fused to a reverse transcriptase. Upon binding to the target DNA, one of the target DNA strands is nicked, the RT template is reverse-transcribed and the new DNA sequence is inserted at the edit position utilizing endogenous host DNA repair machinery. b) Two mutations in the mGreenLantern amino acid sequence (G65S and Y66H) shift its fluorescence from green to blue. Prime editing restores the original sequence, reverting the fluorescence to green. c) Left: HEK293T cells transfected with plasmid DNA encoding pegRNA and prime editor on two separate plasmids. mScarlet-I is co-expressed from the same plasmid as the prime editor as a control. Scale bar: 100 µm. Right: HEK293T cells transfected with pegRNA and prime editor mRNA (capped, polyadenylated, and transcribed with pseudouridine), co-expressing mScarlet-I. Scale bar: 100 µm. d) HEK293T cells seeded in 15% GelMa were transfected three days post-incubation, and images were acquired 24 h later to assess fluorescent protein expression. Left (pDNA): Composite images of brightfield and RFP channel, and composite image of brightfield and GFP channel of cell clusters in GelMa, for which cells were transfected with plasmid DNA, showing expression of mScarlet-I and mGreenLantern in different clusters. Right (mRNA): Corresponding composite images of cell cluster after transfection with mRNA. Expression of mScarlet-I is barely detectable (see faint mScarlet-I signal highlighted at the upper left side of the cell cluster), whereas mGreenLantern expression is clearly visible. Scale bar: 100 µm for all images.

For readout, we used a genetically modified cell line that constitutively expresses a mutant mGreenLantern, in which two amino acids were altered at positions 65 and 66 (G65S and Y66H), resulting in a weakly blue fluorescing mutant, similar as was shown for GFP by Heim et al. (*24*). Prime-editing can be used to revert this mutation and thus recover fluorescence emission in the green (Figure 5b).

To enable direct monitoring of its expression, mScarlet-I was encoded on the same transcript as the prime editor. As in our transfection experiments, we investigated the delivery of the prime editor using both plasmid DNA and RNA vectors. For pDNA delivery, we transfected cells with one plasmid encoding the prime editor and mScarlet-I, and a second plasmid encoding the pegRNA. For RNA lipoplexes, we transfected cells with mRNA encoding the prime editor and mScarlet-I, mixed with pegRNA in a ninefold excess and packaged using Lipofectamine 2000.

The presence of green fluorescent cells confirmed successful prime editing with both pDNA and mRNA (Figure 5c). However, pDNA transfection appeared to yield a higher proportion of successful prime-editing events. Notably, in pDNA-transfected cells most cells expressed mScarlet-I ≈ 6h after transfection, but initially lacked a green mGreenLantern signal, which appeared with a time delay of ≈ 4h. This can be attributed to the sequential nature of protein expression: the prime editor must first be transcribed, simultaneously with mScarlet-I, and translated before the actual genomic editing occurs, leading to a delayed mGreenLantern signal. In contrast, mRNA-transfected cells exhibited mScarlet-I expression almost immediately, while an mGreenLantern signal became detectable after ≈ 6h. In this case, the prime editor must first be translated from the mRNA before it can bind to the co-transfected pegRNA. As a result, prime editing efficiency is influenced by pegRNA degradation within the cell, contributing to the overall lower prime editing efficiency observed.

As shown in Figure 5d, we observed successful editing events also in gel-embedded cell clusters. However, the fraction of cells expressing a fluorescent protein after prime editing appeared to be lower than after simple transfection with a mCherry encoding plasmid (cf. Figure 4). Crucially, prime editing leads to a permanent change in the genome of the cells and thus has a permanent effect on gene expression than the transient transfection with pDNA or mRNA. This is particularly evident in the lower mScarlet-I signal following mRNA delivery, whereas the recovered mGreenLantern signal in successfully edited cells remains comparable to the intensity level of pDNA-modified cells. This stands in stark contrast to the vastly different fluorescence levels of both reporters observed in the transfection experiments.

### Stimulation of Hydrogel-Embedded Cells via Channel Perfusion

Diffusion limitations prevent tissues above a certain size from being functional without vascularization. Distances larger than a few hundred micrometers cannot be sufficiently supplied with nutrients, signaling molecules, and other reagents by passive diffusion. To investigate how cells would grow in a hydrogel that was vascularized with a channel structure, we seeded cells in hydrogel around a channel with a diameter of 600 µm and flushed it with culture medium at a flow rate of 200 µL/h (Figure 6a). Even though the cells were initially homogeneously distributed throughout the gel matrix, the cells only formed clusters in proximity to the channel. A region of ≈ 500 µm around the channel was densily populated by cell clusters. This shows that the supply with nutrients over channel structures increases the area in which cell proliferation can be observed. The region increases from 200 µm in well plate experiments to a final tissue thickness of 1600 µm including the channel with diameter of 600 µm. When substracting the channel the tissue height increased still by 5 fold. This can be attributed to the constant supply with fresh medium as well as the removal of waste products.

**Figure 6:**
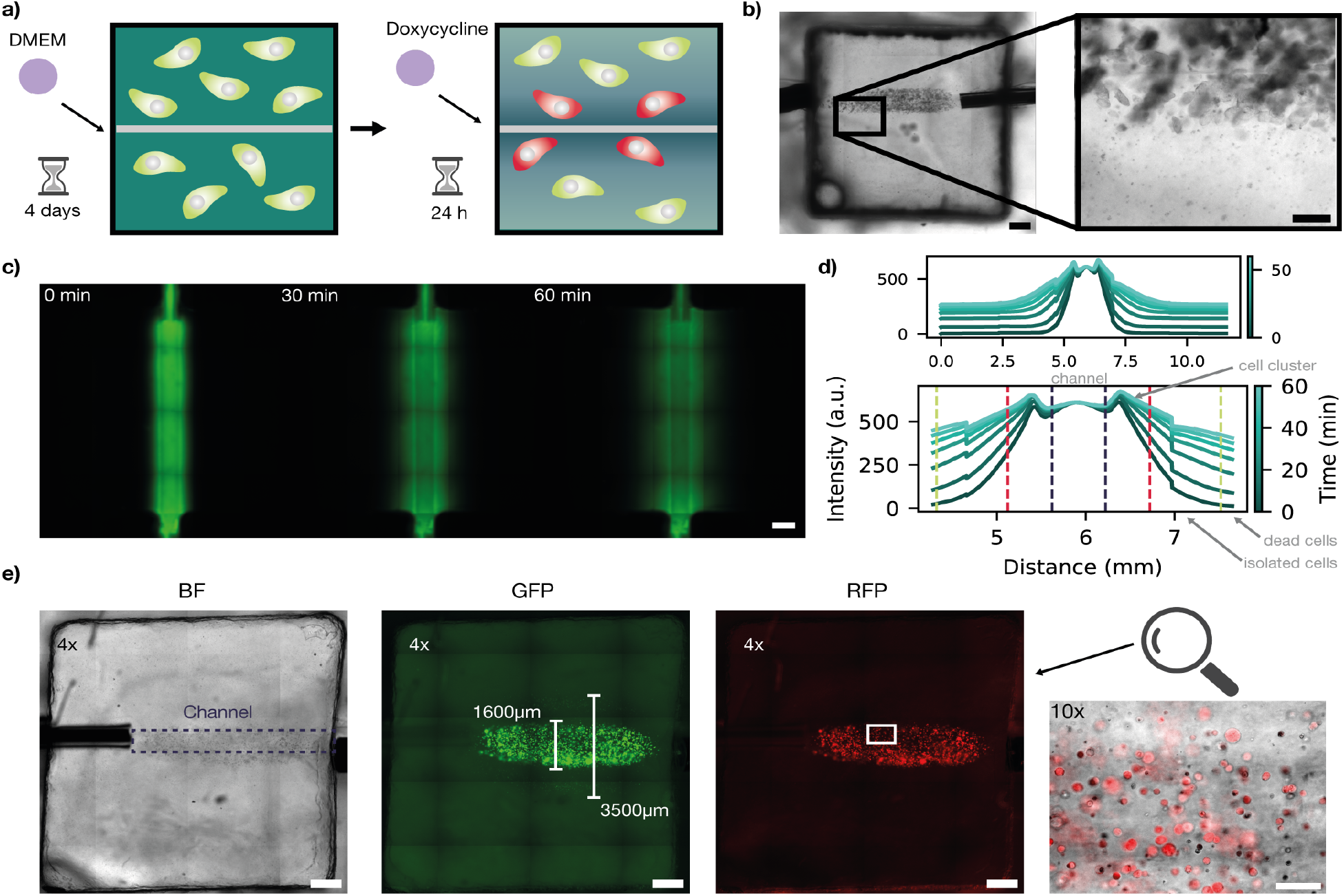
Construction of vascular-like structures. a) Schematic representation of the experiment. Cells were mixed with GelMa (Cellink) containing 0.25% LAP and seeded in a homebuilt bioreactor. DMEM was perfused through the channel at a flow rate of 200 µL/h for four days. Doxycycline at a concentration of 500 ng/ml was introduced into the medium and perfused for 24 h. b) Brightfield image of the printed channel. Cells proliferate around the channel. Scale bar: 1 mm. c) To investigate the diffusion of small molecules in the hydrogel matrix, fluorescein was perfused through the channel. Representative images at 0 min, 30 min, and 60 min are shown. Scale bar: 1 mm. d) Time evolution of the fluorescein diffusion profile around the channel within 1 h. Images were acquired every 5 min. The dashed lines indicate regions with different growth conditions - purple: central channel, red: region with high concentration of nutrients supporting cell growth, green: nutrients support growth of isolated cells. Outside of the region marked with green dashed lines, cells are not viable. e) From left to right: Brightfield, green and red fluorescence images of the bioreactor. The central channel is barely visible, and therefore indicated with a dashed line in the brightfield image. Cells extend up to a distance of 1750 µm from the channel center, regions with high cell density and larger cell clusters are confined within 800 µm from the channel. The rightmost image provides a magnified view of this region in the RFP channel, showing that cells within its proximity express mScarlet-I upon addition of doxycycline. Scale bar: 1 mm.

To assess the diffusion profile in the gel we introduced fluorescein by flushing the central supply channel with the dye at a constant flow rate of 200 µL/h. Fluorescein has a size comparable to that of small metabolites in the culture medium. Figure 6c shows fluorescence images of the vascularized gel taken at different time points, the corresponding mean fluorescence values in the direction perpendicular to the channel are plotted in Figure 6d.

We used the channel structure to regulate gene expression in gel-embedded HEK293T cells via a diffusible inducer. Cells were seeded in the bioreactor, cultured under continuous DMEM flow for four days, and then exposed to 500 ng/mL doxycycline to induce gene expression.

The diffusion zone (red-dashed lines in Fig. 6d) closely matched the region where cells proliferated and formed clusters, extending up to approximately 800 µm from the channel center. Beyond this, isolated live cells were detected up to 1750 µm (green-dashed line, Fig. 6d), expressing fluorescent proteins but not forming clusters. Further from the channel, gene expression and proliferation ceased.

For future applications, more advanced channel structures will be needed to optimize nutrient supply, waste removal, and spatially controlled gene expression via diffusible inducers. As a proof of concept, we implemented a 3D bioprinting strategy using a sacrificial ink approach (*25*) to fabricate vascular-like channels. This allowed us to investigate complex vascular architectures and assess their influence on small molecule diffusion within the gel matrix (cf. Supplementary Information S11–S13).

## 3 Conclusion

Our study demonstrates that GelMA hydrogel scaffolds support cell proliferation, localized stimulation, and genetic manipulation in a 3D environment. HEK293T cells embedded in GelMA remained viable and formed dense clusters over 18 days with minimal cell death. Additionally, vascular-like supply channels facilitated controlled molecular delivery and enabled in situ genetic manipulation of matrix-embedded cells. We systematically progressed from chemical gene induction to more advanced genetic modifications, evaluating how embedded cells respond to external stimuli. By delivering genetic constructs via both plasmid DNA and mRNA, we demonstrated conventional fluorescent protein expression as well as in situ prime editing, leading to permanent genetic modifications. The emergence of the final gene product follows distinct kinetic processes: mRNA enables rapid expression but has a short half-life, whereas plasmid DNA requires multiple processing steps, introducing a time delay. When combined with mRNA delivery, prime editing provides a transient initial stimulus while inducing a permanent genomic modification in targeted cells. We also explored how embedded channels within GelMA hydrogels influence spatial control over gene expression and molecular transport. While vascular-like structures have the potential to improve nutrient delivery and gene induction, further optimization will be necessary to enhance their physiological relevance. Future research has to focus on refining hydrogel properties, optimizing gene delivery strategies, and improving molecular transport dynamics through synthetic vasculature to better mimic native tissue environments. Advancing these aspects could further drive applications in tissue engineering, regenerative medicine, and drug testing.

## 4 Experimental Section

### Cell lines and Plasmids

Three HEK293T cell lines were used for growth, transfection, doxycycline induction, and prime editing experiments:

i. A standard HEK293T cell line (ATCC, CRL-3216−) was employed for growth studies and general transfection experiments due to its high transfection efficiency and suitability for protein expression analyses. A plasmid encoding mCherry under the J23119 promoter was used (*26*) for transfection studies. The plasmid map is available in the SI (Figure S15). Two additional genetically modified HEK293T cell lines were engineered using the piggyBac transposon system, which enables stable genomic integration via two plasmids: one encoding the piggyBac transposase and another carrying the gene of interest.
ii. Doxycycline-Inducible Cell Line: This cell line constitutively expresses mGreenLantern and switches to mScarlet-I upon doxycycline induction via the Tet-On 3G system (*22*, *27*, *28*). This system allows for precise control and visualization of inducible gene expression. The plasmids used for stable transfection are provided in the SI (Figure S16).
iii. Prime Editing Cell Line: This cell line was designed for prime editing studies by modifying the green fluorescent protein (GFP) in G65S and Y66H, resulting in a hypsochromic spectral shift from green to blue fluorescence. Successful prime editing restores the original SH sequence, reverting fluorescence back to green. This fluorescence switch provides a visual and quantitative readout of editing (*29*). Plasmid maps used for prime editing of the prime editor and pegRNA can be found in the SI (Figures S17-S23).

### Plasmid Preparation and RNA Synthesis

Plasmids were prepared using the ZymoPURE Plasmid Miniprep Kit (Zymo Research) following the manufacturer’s protocol without modifications. The purified plasmid DNA was used directly for experiments or as a template for in vitro transcription (IVT) to generate mRNA.

For mRNA synthesis, the RiboMAX− Large Scale RNA Production System (Promega) was used. Plasmid DNA templates were linearized with AsiSI and BasI-HF-v2 (NEB) to ensure proper transcription termination. Following digestion, DNA was purified using the DNA Clean & Concentrator kit (Zymo Research) and used as transcription template.

IVT was performed according to the manufacturer’s instructions, incorporating pseudouridine (Φ) (Jena Bioscience) for enhanced RNA stability for the prime editor only. Afterwards, plasmid DNA was removed through DpnI digestion and RNA was isolated using the RNA Clean & Concentrator kit (Zymo Research). Transcribed pegRNA was stored directly at −80 °C and mRNA for the prime editor was capped with the FCE Capping system (NEB) and polyadenylated using Poly A Polymerase (NEB) to improve translational efficiency. The processed RNA was purified using the RNA Clean & Concentrator kit (Zymo Research), quantified with a Nanophotometer (Implen), and stored at −80 °C until use.

### Cell Culture Conditions

Cells were initially seeded in 75 cm^2^ cell culture flasks in Dulbecco’s Modified Eagle Medium (DMEM, high glucose, Gibco™) supplemented with 10% v/v Fetal Bovine Serum (FBS, Sigma-Aldrich) and 1% Penicillin-Streptomycin (P/S, Gibco™) to support growth and prevent bacterial contamination. Cultures were maintained at 37 °C with 80% humidity and a 5% CO_2_ atmosphere. The medium was fully exchanged every 3 to 4 days to replenish nutrients and remove waste products. Upon reaching 80% confluence, cells were passaged using 0.25% Trypsin-EDTA (Gibco™) and maintained up to passage 20.

For three-dimensional culture, cells were encapsulated in GelMa (gelatin methacrylate) prewarmed to 37 °C to facilitate mixing. To achieve a final concentration of 1 *×* 10^6^ cells/mL, 1 mL of GelMa was combined with 100 µL of a cell suspension containing approximately 1.1 *×* 10^6^ cells. The mixture was homogenized by gentle pipetting, then transferred to a well plate or bioreactor depending on the experiment. Constructs of varying thickness (1 mm to 3 mm) were crosslinked under 405 nm UV light for 10 s to 30 s, depending on the thickness.

Two types of GelMa were used: a homemade formulation adjusted to 15% with 0.25% Lithium phenyl-2,4,6-trimethylbenzoylphosphinate (LAP, Sigma-Aldrich) for well plate experiments, and commercially sourced GelMa (GelMa A, Cellink) containing 0.25% LAP for channel experiments. Both formulations exhibited similar behavior above 30 °C, as confirmed by rheometer analysis (see SI).

### Transfection of HEK293T Cells

Transfection of HEK293T cells was performed to introduce genetic material and assess the efficiency of different transfection reagents in 2D and 3D environments. In 2D cultures, DNA transfection was carried out using jetOPTIMUS (Polyplus), while RNA transfection was performed with Lipofectamine 2000 (Invitrogen−). In 3D GelMa hydrogels, DNA and RNA transfection efficiencies were directly compared using Lipofectamine 2000 (Invitrogen−). The amounts for buffer, DNA, and the transfection reagent were used according to the manufacturers protocol for DNA transfection. A 24-well plate was used and 0.7 µL of reagent were used. For RNA transfection with Lopfectamin amounts for transfection reagent and Optimem were also used according to the manufacturers protocol for a 24-well plate. Slightly higher amounts of RNA were used compared to the manufacturers protocoll: 800 ng instead of 500 ng. When transfecting RNA for prime editing 720 ng of the RNA for the prime editor and 80 ng of pegRNA were used. Transfection efficiency was determined with a flow cytometer (Attune NxT, Invitrogen) to yield ≈ 63%. The data for transfection with the plasmid encoding mCherry can be found in the SI (Figure S4 & S5).

### Homemade GelMa

Homemade GelMa foam was synthesized using 10 g of Type A gelatin (300 bloom, Sigma-Aldrich) which was dissolved in 100 ml of 1x phosphate-buffered saline (PBS tablets, Sigma-Aldrich) at room temperature with moderate stirring, then heated to 50 °C in a water bath until fully dissolved. While stirring continuously, 0.6 g of methacrylic anhydride (MAA, Sigma-Aldrich) per gram of gelatin was gradually added using a glass pipette to prevent plastic interaction. The reaction proceeded for 3 h under continuous stirring to achieve a high degree of methacryloyl functionalization (75(9)%) according to the protocol of Loessner et al. (*30*).

Following the reaction, the solution was centrifuged at 3500 g for 3 min to remove unreacted MAA. The supernatant was diluted with two volumes of preheated (40 °C) 1x PBS and transferred into a 12 kDa molecular weight cut-off dialysis membrane. Dialysis was performed against 5 L of 1x PBS at 40 °C for 5 to 7 days, with daily PBS changes.

After dialysis, the pH was adjusted to 7.4 using 1 M NaHCO_3_. The solution was filter-sterilized using 0.2 µm syringe filters, aliquoted into 50 ml tubes, and snap-frozen in liquid nitrogen. Without thawing, aliquots were lyophilized for 4 to 7 days until fully dehydrated. The lyophilized GelMa foam was stored at −20 °C in sealed screw-top tubes, protected from light and moisture.

### Peparation of Bioinks

For preparing the homemade GelMa solution, lyophilized GelMa foam was sterilized with UV light for 30 minutes before use. A 1.25% LAP stock solution was prepared by dissolving 50 mg of LAP in 4 mL of DPBS (Gibco™) under constant stirring at 70 °C for 30 minutes, with the solution protected from light using aluminum foil. The appropriate volume of LAP stock solution was then added to achieve a final concentration of 0.25% (e.g., 2 mL LAP stock for 10 mL total GelMa solution). Separately, GelMa precursor solution was prepared by dissolving the desired amount of lyophilized GelMa foam in PBS at 70 °C for 30 minutes under magnetic stirring (e.g., 1.5 g GelMa in 8 mL PBS for a 15% GelMa solution). Finally, the LAP and GelMa solutions were combined and stirred at 60 °C for 10 minutes to produce the final hydrogel solution.

### Bioreactor setup

The bioreactor system was constructed using custom chambers made of polydimethylsiloxane (PDMS, SYLGARD 184, Biesterfeld) to support printed GelMa structures. Each PDMS chamber was bonded to a glass microscope slide (VWR) via oxygen plasma treatment, providing a stable and sealed base. A needle tip was punched through the bioreactor chamber, and GelMa mixed with cells was cast within the PDMS chamber. A second glass slide (coverslip, 18×18 mm, DWK Life Science) was then secured on top using silicone glue (SI 595, Loctite, Henkel), creating a fully enclosed environment for cell culture. The glue was selected for its ability to bond glass and PDMS while maintaining biocompatibility. Biocompatibility was confirmed experimentally (see SI Figure S10).

The bioreactor was cross-linked under 405 nm UV light for 30 s and incubated for 5 min to allow glue solidification. The needle was then carefully removed, and catheters (indwelling venous cannula) were inserted through the PDMS to create inlets and outlets for media flow. A 20 mL syringe (Braun) filled with DMEM (or Fluorescein for the diffusion experiments in Figure1) was connected via tubing (extension line, Braun) to the catheters, and channels were flushed with DMEM. Media flow was maintained using a programmable multichannel syringe pump (Darwin) at a flow rate of 200 µL/h. The direct catheter-to-PDMS connection established a continuous, airtight flow system, facilitating nutrient exchange and supporting cellular growth within the GelMa matrix. Images of the setup are provided in the SI (Figure S14).

### Microscopy

Fluorescence imaging was performed using an M7000 microscope (Evos) equipped for live-cell imaging. To maintain optimal cell viability during imaging, samples were kept at 37 °C with 80% humidity and 5% CO_2_ using a Stage Top Incubation System, simulating physiological conditions. The microscope was equipped with GFP, RFP, and DAPI filter sets.

For live/dead analysis, cells were stained with Hoechst 34580 (Invitrogen) to label nuclei in live cells and Propidium Iodide (Life Sciences) to identify dead cells with compromised membrane integrity. This dual-staining approach enabled the assessment of cell viability within 3D GelMa constructs under real-time conditions.

### Data Analysis

Data analysis was performed using Fiji (ImageJ, version 1.54f) to process and quantify fluorescence images. Cell viability, spatial distribution, and expression levels were assessed through fluorescence intensity measurements. For doxycycline titration experiments, regions of interest (ROIs) were selected to ensure 100% confluence over time. Cell clusters for growth analysis were manually selected, and their sizes were approximated by fitting ellipsoids. Ten clusters per sample were analyzed for statistical evaluation. Representative images illustrating ROI selection and cluster identification are provided in the SI S1 & S2.

## Supporting Information

Supporting Information is available from the Wiley Online Library or from the author.

## Acknowledgements

The authors thank Christoph Schmidt and Tatiana Kovalchuk for their help with hydrogel preparation, 3D printing and cell culture; Magdalena Bock for her help with the rheometer measurements; Samuel Beerkens for his help with flow cytometer measurements; Gil Westmeyer for providing the mammalian cells and plasmids, and Julian Geilenkeuser for his help in plasmids and cell line creation.

Figures were partly created in BioRender. Jäkel, A. (2025) https://BioRender.com/z13c262 & https://BioRender.com/c98m883.

## Data Availability Statement

Raw microscopy data is available upon request from the authors.

