## Supplementary Information for "Genetic Manipulation of Mammalian Cells in Microphysiological Hydrogels"

### Genetic Manipulation of Mammalian Cells in Microphysiological Hydrogels Supporting Information

#### Doxycycline Titration

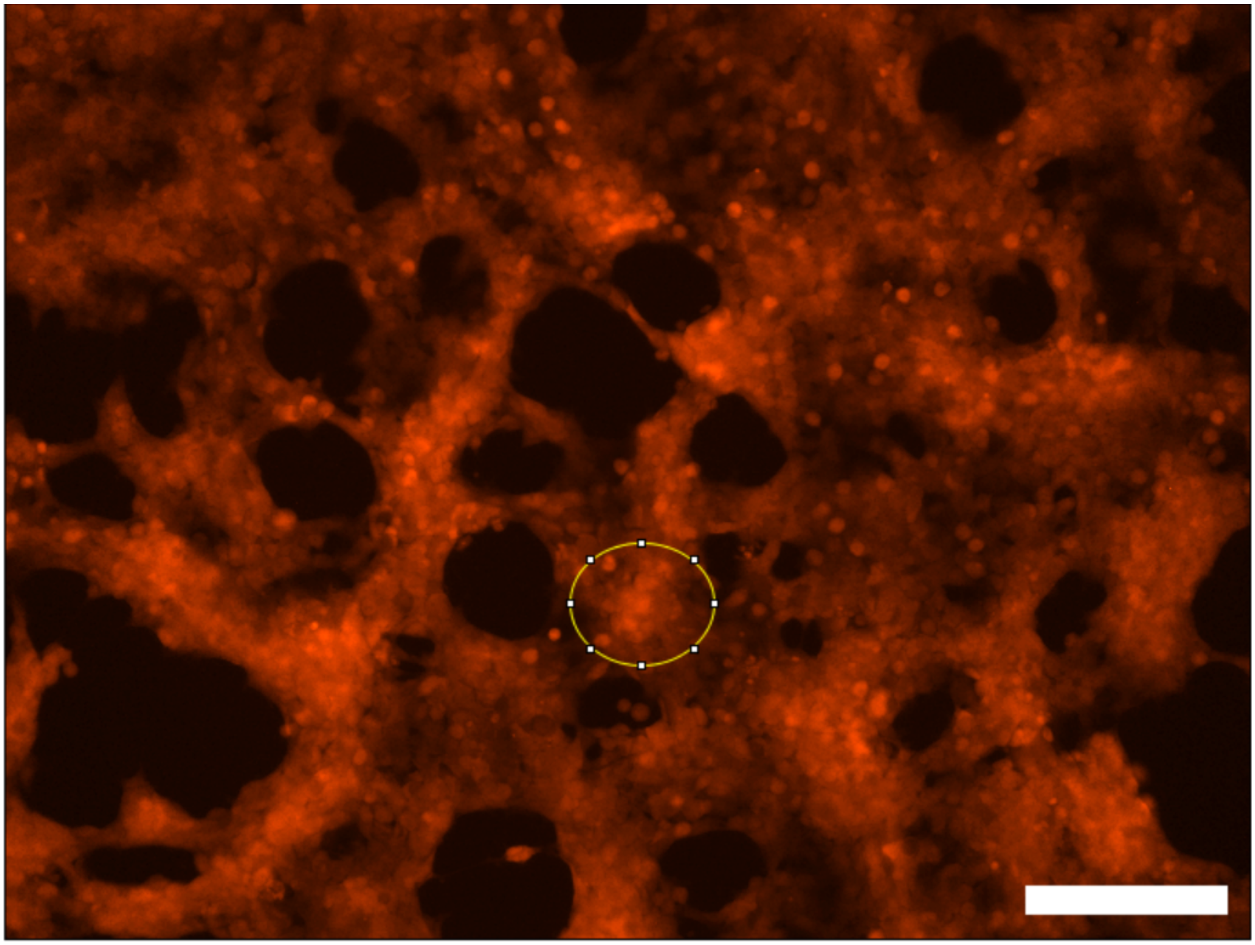

Figure S1: A region of interest (ROI) was selected for each sample that was $100\,\%$ confluent. The following parameters were measured for over a period of $60$ h: area, mean intensity, standard deviation, min and max intensity. Measurements were taken every $2$ h for the first $24$ h, then after $24$ h, and then after $12$ h. Scalebar: $100$ µm

#### Cell growth in GelMa

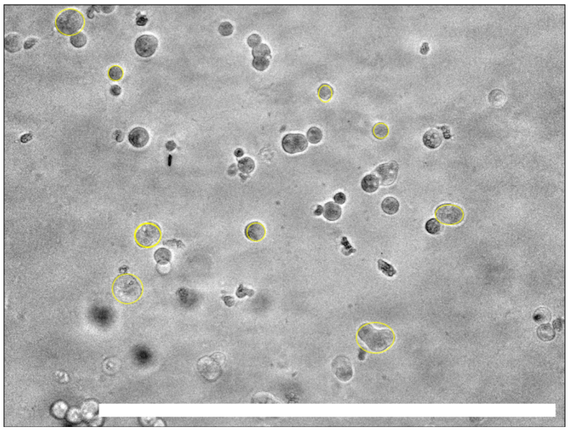

(a) Day 1 after seeding cells in GelMa.

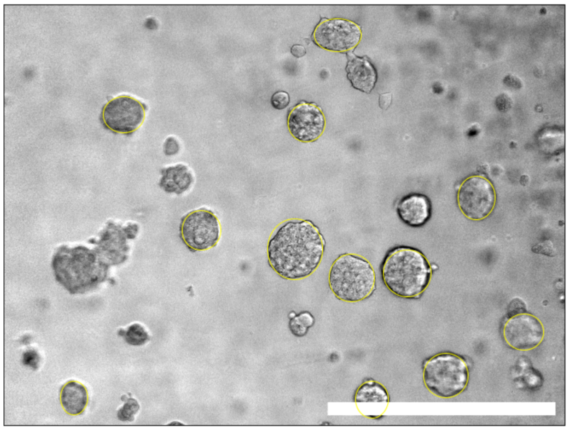

(b) Day 5 after seeding cells in GelMa.

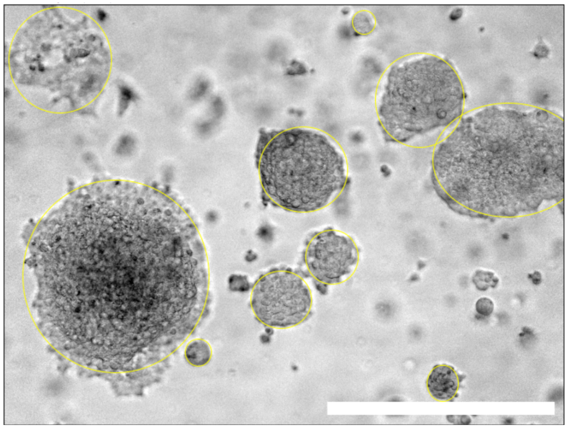

(c) Day 8 after seeding cells in GelMa.

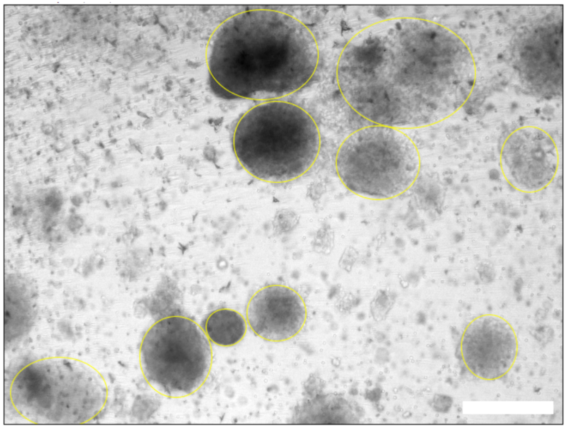

(d) Day 16 after seeding cells in GelMa.

Figure S2: For each sample 10 clusters were selected to measure cluster size. The feret radius was determined with Fiji and the mean was calculated for each sample. Scalebar: $500\,\mu m$

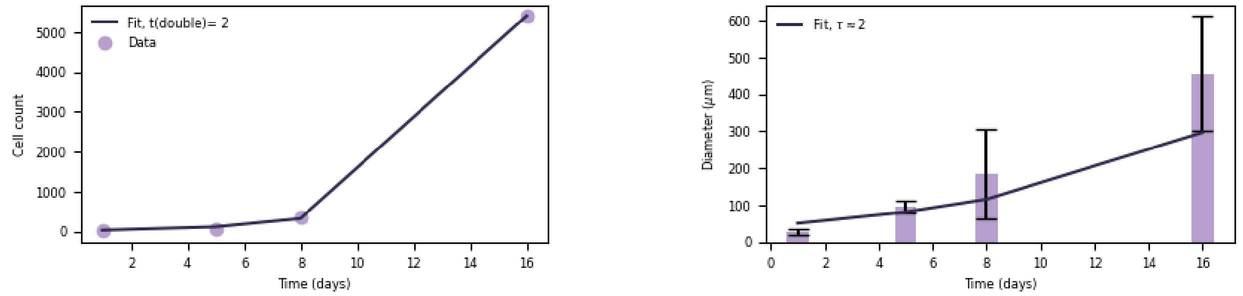


(a) Number of cells calculated in one cell cluster over time.

(b) Cell cluster diameter over time. Fitting the data results in a doubling time of ≈ 2 days.

Figure S3: The cell count $N_c$ was calculated assuming a spherical volume of each cluster. And the doubling time was calculated using the formula $N_c(t) = N_0 \times 2^{t/t_D}$.

### Transfection

#### Transfection of mCherry - FACS

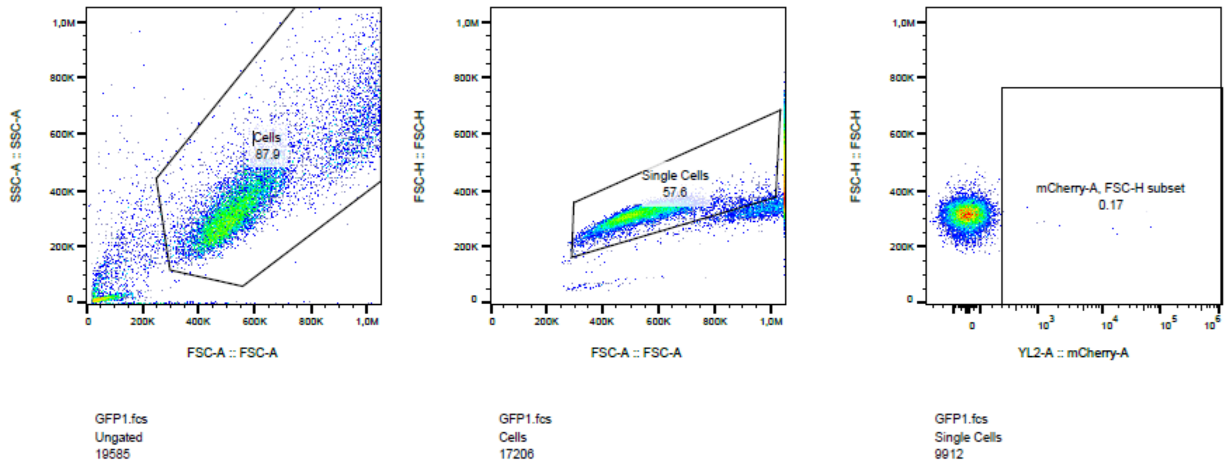


Figure S4: Control measurement of HEK293T cells.

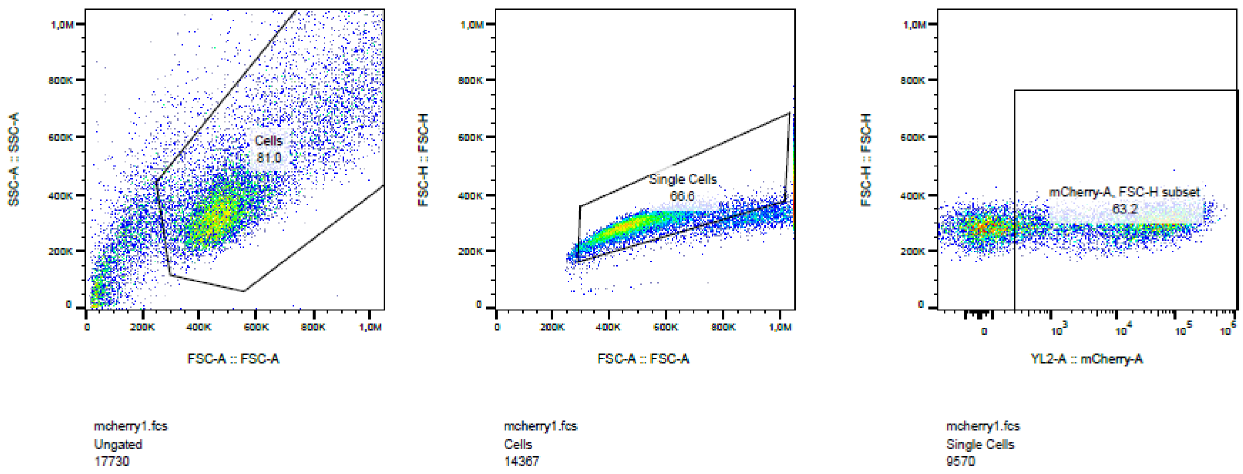


Figure S5: HEK293T were transfected using jetOptimus transfection reagent with plasmid DNA encoding for the fluorescent protein mCherry.

#### Rheometer Measurements

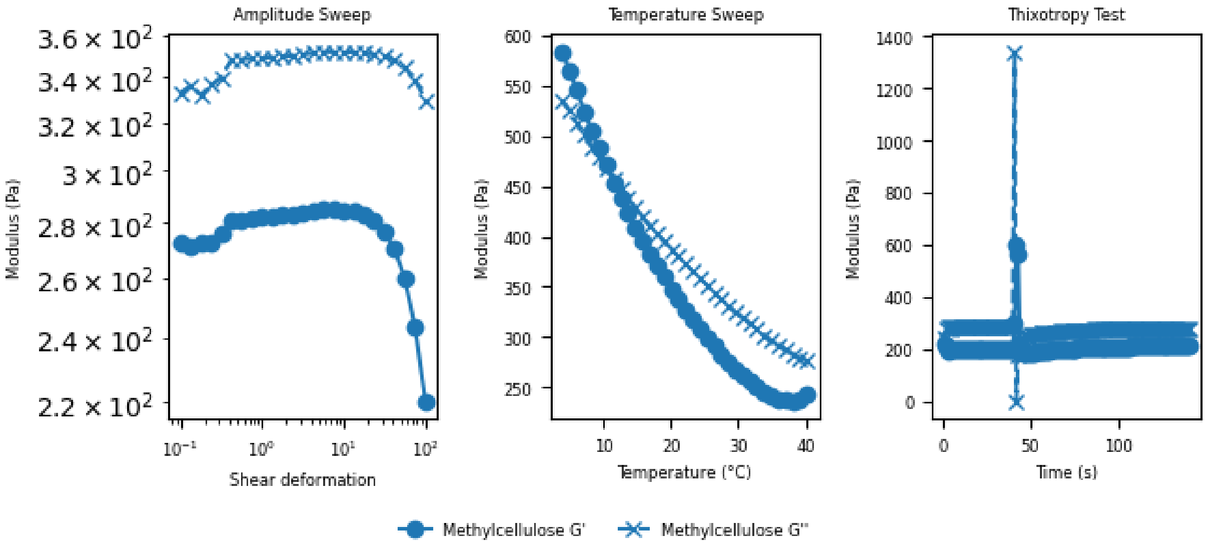


Figure S6: Methylcellulose ($5\,\%$ v/w ) was measured as benchmark. The amplitude sweep provides insight into the viscoelastic properties of methylcellulose by evaluating its storage modulus ($G'$) and loss modulus ($G''$) across increasing strain amplitudes. In the linear viscoelastic region (LVR), $G''$ remains higher than $G'$, indicating that the material can be described as a viscoelastic liquid. As strain increases beyond the critical strain, $G'$ decreases, and the limit of the LVE-region is reached. This suggests that methylcellulose maintains its gel-like properties under low deformation but undergoes yielding at higher strains. The temperature-dependent rheological behavior of methylcellulose is characterized by a decrease in $G'$ at higher temperatures, indicative of thermogelling properties. Initially, at lower temperatures, the material behaves more elastic with $G' > G''$. As temperature rises, the viscous proportion predominates, reflected in a crossover point where $G'$ surpasses $G''$. This transition temperature is a critical parameter for applications requiring thermal responsiveness. The thixotropy test assesses the recovery of methylcellulose after shear-induced breakdown. Upon applying high shear, both $G'$ and viscosity drop significantly, reflecting structural disruption. When shear is reduced, a gradual recovery of $G'$ is observed, indicating partial structural reformation. However, if full recovery is not achieved within the measured timeframe, this suggests that methylcellulose exhibits a degree of irreversible structural breakdown or slow rebuilding dynamics. This behavior is relevant for applications where shear-induced fluidization and recovery kinetics are important.

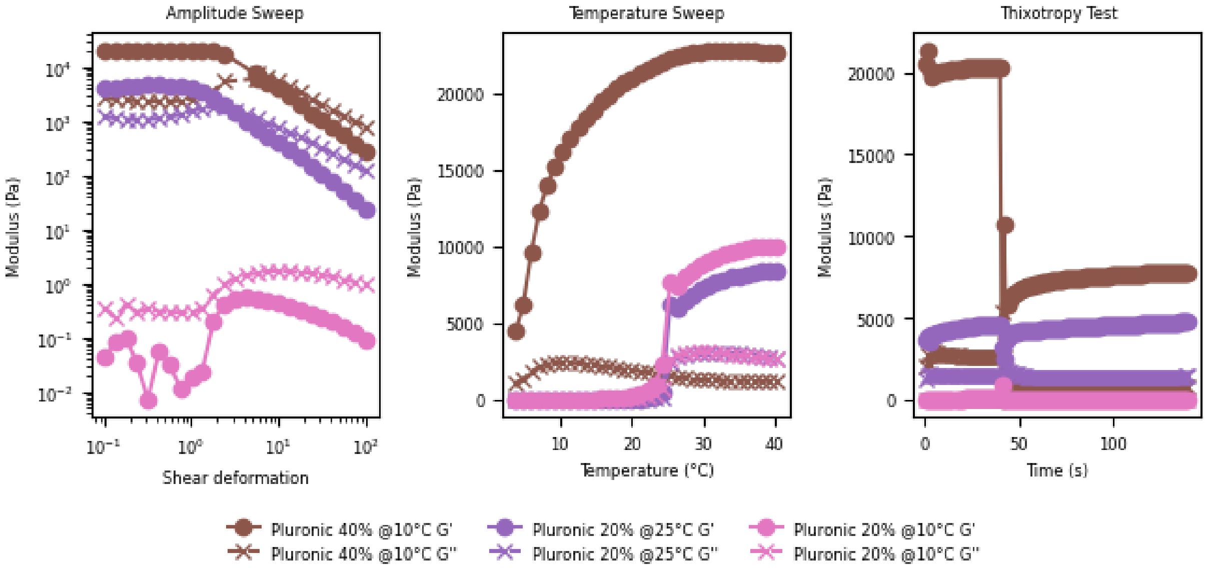


Figure S7: Pluronic is printable at $20\,\%$ and $40\,\%$ at room temperature for $\sim 15\,$min when stored before at $-4\,°$C. An amplitude sweep of pluronic hydrogels at different concentrations and temperatures are shown. Compared to methylcellulose, the pluronic curves exhibit a less steep decline in the storage modulus ($G'$) and loss modulus ($G''$) with increasing strain, indicating a more gradual structural breakdown. Additionally, pluronic $20\,\%$ at $10\,°$C shows significantly lower $G'$ values compared to pluronic $20\,\%$ at room temperature or pluronic $40\,\%$, suggesting a temperature-dependent shift in mechanical strength. This behavior reflects the thermoresponsive nature of pluronic, where gel stiffness varies with both concentration and temperature. Interestingly pluronic with a concentration of $40\,\%$ at $10\,°$C behaves similar as pluronic with a concentration of $20\,\%$ at $25\,°$C. The temperature sweep of pluronic hydrogels at different concentrations shows that in contrast to methylcellulose, which shows a more linear increase in modulus with temperature, pluronic exhibits a non-monotonic "hill-shaped" curve. The storage modulus ($G'$) initially increases with rising temperature, reaches a peak, and then declines at higher temperatures. This trend suggests a gelation process followed by structural weakening, likely due to micellar rearrangement or phase separation. Additionally, the difference between storage ($G'$) and loss modulus ($G''$) is more pronounced compared to methylcellulose, highlighting distinct viscoelastic behavior and phase transitions in pluronic hydrogels. The thixotropy test shows that plurnic at $40\,\%$ does not recover its structure and viscosity as the $20\,\%$ pluronic does.

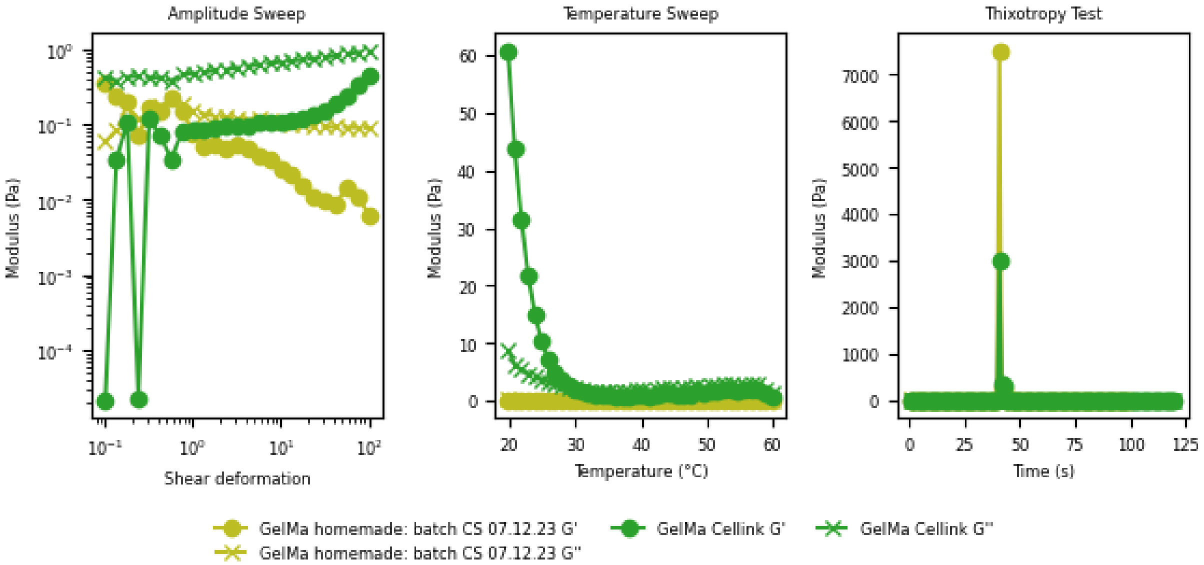


Figure S8: Rheological analysis of GelMa purchased from Cellink and our homemade GelMa. Amplitude-, temperature sweep, and thixotropy test. The temperature sweep shows similar behaviour of the hydrogels above $30\,^{\circ}$C. The thixotropy test shows that both hydrogels recover its structure and viscosity after deformation.

### Cell viability in different gels

#### Cell line test

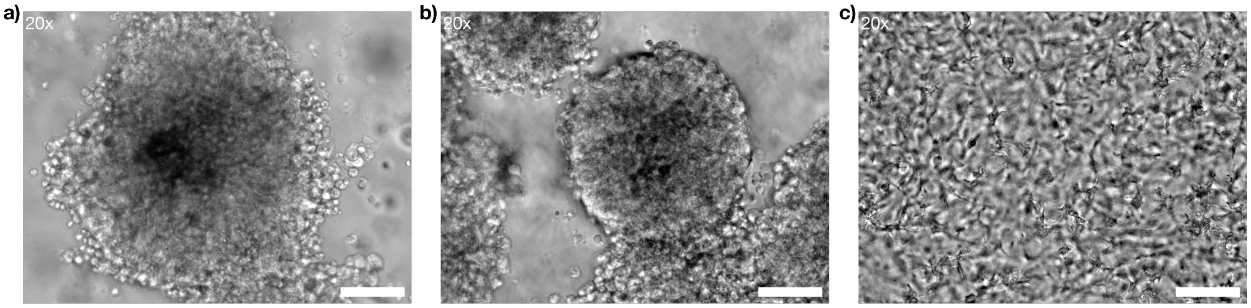


Figure S9: Different celltypes grown in GelMa for 16 days: a) HEK293T b) NIH c) hMSC. Scalebar: $100\,\mu m$.

#### Compatibility with glue

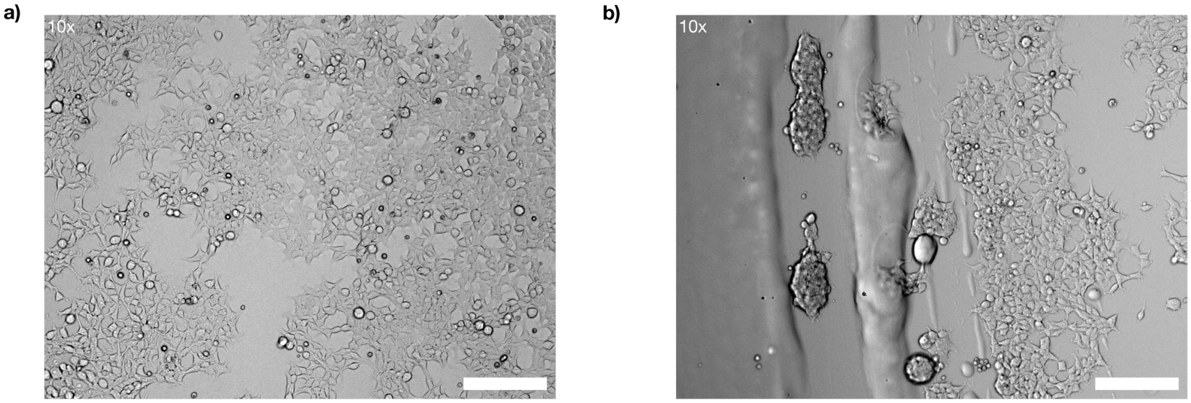


Figure S10: a) Cell growth in a well plate next to a glue drop. b) Cell growth inside the gel drop. Cells do grow in clusters similar to cells growing in GelMa. Scalebar: $200\,\mu m$.

### Test of nozzle precision

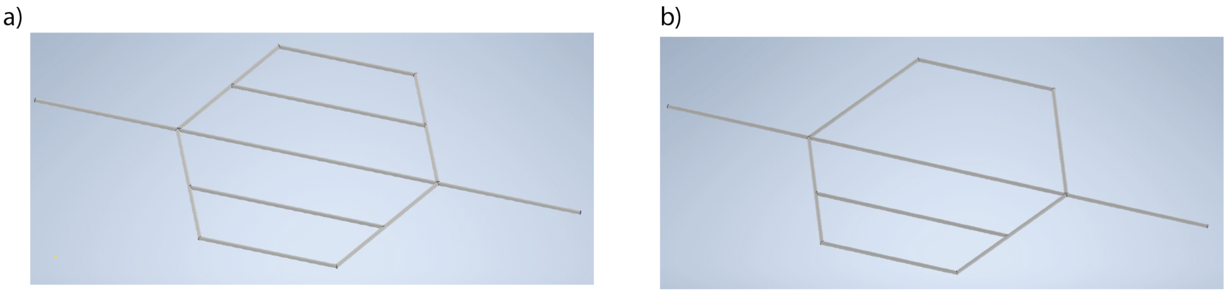


Figure S11: Printed vascular structures. a) Even distribution network. b) Uneven distribution network.

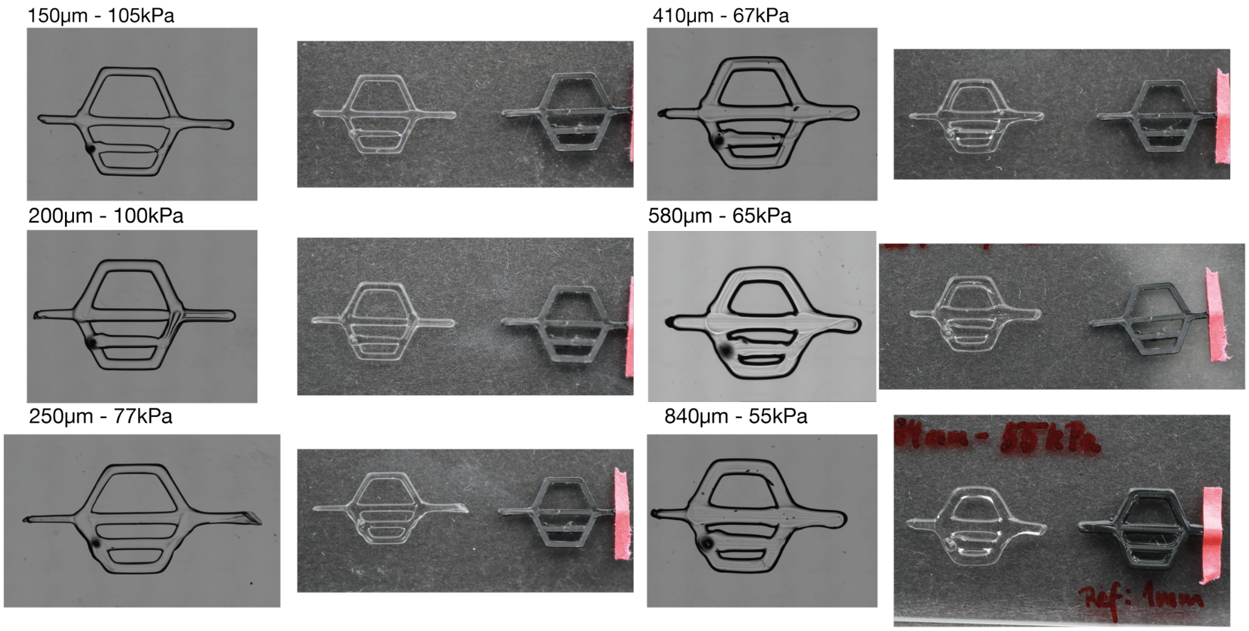


Figure S12: Channel structures printed with a BIOX2 printer (Cellink). Different needle sizes were tested and images are presented of the structure for each needle with the optimal pressure value. Right: microscope image, left: photograph of the structure next to a PLA printed construct which was printed with a Bambulab X1E printer. The channel thickness was 1 mm to compare resolution.

#### Fluorescein Channel Experiments

To fabricate vasculature-like structures, a $40\,\%$ (w/v) Pluronic F-127 solution was prepared by dissolving pluronic powder (Sigma-Aldrich) in double-distilled water (dd$H_2O$) at $4\,°$C overnight, with periodic vortexing to ensure complete dissolution. This temperature-responsive bioink served as a sacrificial support material, facilitating the creation of vascular channels within the GelMa matrix.

A BIOX 3D bioprinter (Cellink) equipped with a 200 μm nozzle was used to print the vascular structures. The pluronic solution, stored at $4\,°$C, was printed at room temperature, maintaining its optimal viscosity. To prevent premature gelation, printing was performed within 15 minutes of removing the solution from cold storage. If the solution warmed beyond a critical threshold, re-cooling was necessary before continuing the process. Printing parameters were optimized, including a pressure of $100\,$kPa, a printing speed of $5\,\mathrm{mm\,s^{-1}}$, and a preflow adjustment of $-50\,$ms to ensure controlled extrusion.

The bioprinting workflow for generating vascular-like channels is illustrated in Figure S13. First, Pluronic F-127 ($40\,\%$ w/v) was printed as a sacrificial ink to define the channel structure within GelMa. Figure S13a presents a photograph of a pluronic-printed channel stained in red, to highlight the channel structures within the hydrogel. The complete bioprinting process (Figure S13b) involved printing pluronic structures, overlaying the construct with GelMa, and crosslinking the hydrogel using $405\,$nm UV light for $30\,$seconds. Following crosslinking, the constructs were cooled to $4\,°$C for 5 minutes, liquefying the pluronic and enabling its removal, thereby creating perfusable channels.

To validate the functionality of these channels, Figure S13c shows a brightfield image of a pluronic structure before GelMa casting, demonstrating the precision and integrity of the printed channels. Additionally, Figure S13d presents fluorescence imaging of fluorescein diffusion from the channels into the hydrogel, confirming molecular transport into the surrounding matrix. This approach establishes a controlled environment for studying molecular diffusion and cellular responses within engineered hydrogel constructs.

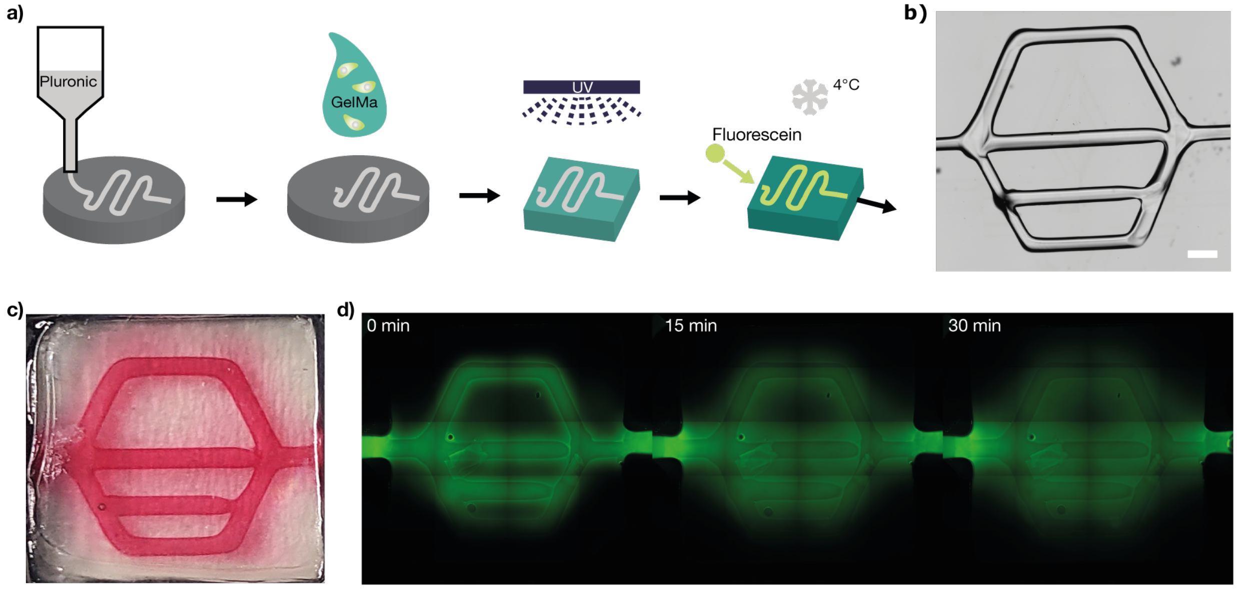


Figure S13: Construction of vascular-like structures. a) Schematic representation of the method for preparing vascular structures using a commercial 3D bioprinter. Pluronic F-127 is printed, followed by casting GelMa with or without cells on top. The GelMa is crosslinked using $405$ nm UV light for $30$ seconds. Subsequently, the construct is cooled to $4\,°$C for 5 minutes, liquefying the pluronic to enable channel perfusion with a material of interest. b) Brightfield image of the printed channel before hydrogel application. Scale bar: $1000$ m. c) Photograph of a constructed channel filled with red food dye to visualize perfusion within the GelMa matrix. d) Channels perfused with fluorescein to observe diffusion into the surrounding hydrogel over time. e) Diffusion of fluorescein from a single channel. Scale bar: $1000$ m. f) Quantitative analysis of fluorescein diffusion.

### Bioreactor Setup

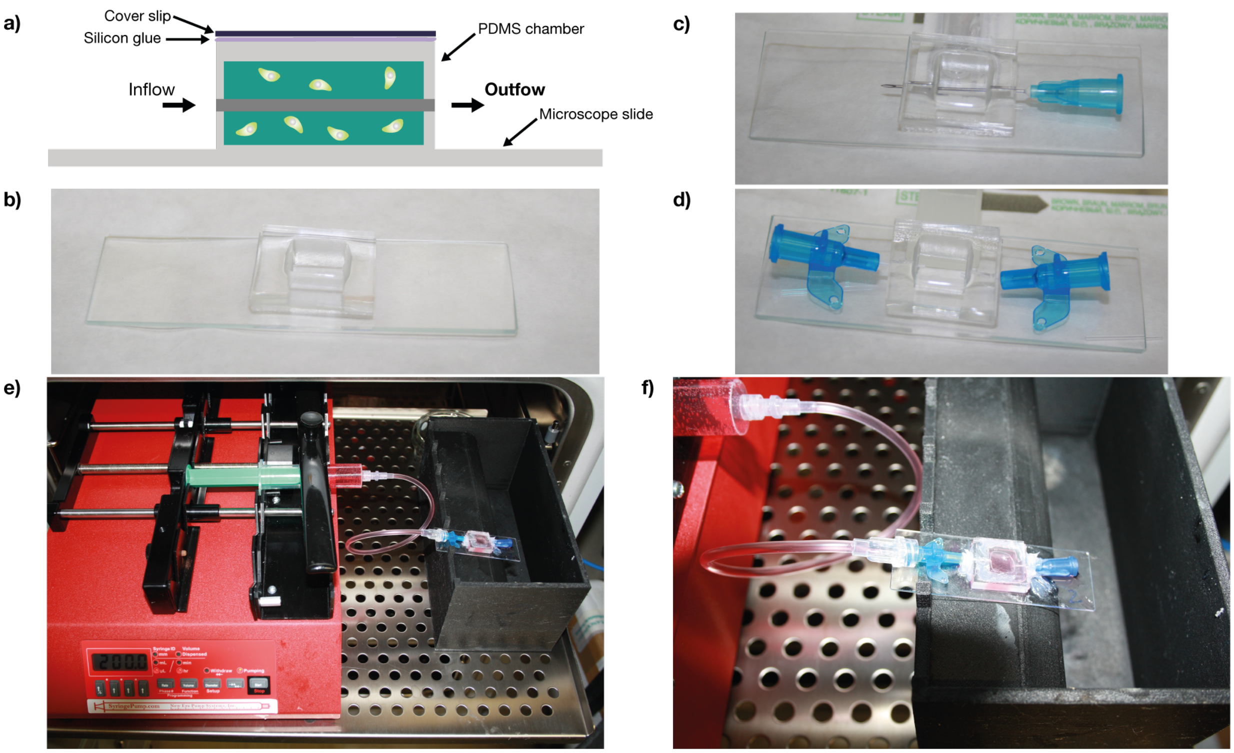


Figure S14: a) Schematic of the bioreactor. b) PDMA chamber bonded to a microscope slide with O2 plasma. c) A needle (ID $0.6\,\mathrm{mm}$) was punched through PDMS for casting gel on top for the creation of a channel. d) In- and outlet connection were achieved with catheters ($22\,\mathrm{G}$). e) Photograph of the whole setup. A syringe pump is used to be able to run 6 bioreactors in parallel. A 3D printed autoclavable PAHT chamber is used as collection tank for material flushed through the bioreactors. f) Zoom in on the bioreator itself.

### Plasmids

#### The full sequence of the plasmid used for transfection studies of mCherry is provided below

cgcgatgtacgggccagatatacgcgttgacattgattattgactagttattaatagtaatcaattacggggtcattagttcatagccca
tatatggagttccgcgttacataacttacggtaaatggcccgcctggctgaccgcccaacgacccccgcccattgacgtcaataatgac
gtatgttcccatagtaacgccaatagggactttccattgacgtcaatgggtggagtatttacggtaaactgcccacttggcagtacatca
agtgtatcatatgccaagtacgccccctattgacgtcaatgacggtaaatggcccgcctggcattatgcccagtacatgaccttatggg
actttcctacttggcagtacatctacgtattagtcatcgctattaccatggtgatgcggttttggcagtacatcaatgggcgtggatagc
ggtttgactcacggggatttccaagtctccaccccattgacgtcaatgggagtttgttttggcaccaaaatcaacgggactttccaaaat
gtcgtaacaactccgccccattgacgcaaatgggcggtaggcgtgtacggtgggaggtctatataagcagagctctctggctaactag
agaacccactgcttactggcttatcTTGACAGCTAGCTCAGTCCTAGGTATAATGCTAGCgaaattaatacg
actcactatagggagacccaagctggctagcgtttaaacttaagcttggtaccgagctcggatccactagtccagtgtggtggaattcg
ccaccatggtgagcaagggcgaggaggataacatggccatcatcaaggagttcatgcgcttcaaggtgcacatggagggctccgtga
acggccacgagttcgagatcgagggcgagggcgagggccgcccctacgagggcacccagaccgccaagctgaaggtgaccaaggg
tggcccccctgcccttcgcctgggacatcctgtcccctcagttcatgtacggctccaaggcctacgtgaagcaccccgccgacatcccg
actacttgaagctgtccttccccgagggcttcaagtgggagcgcgtgatgaacttcgaggacggcggcgtggtgaccgtgacccagg
actcctccctgcaggacggcgagttcatctacaaggtgaagctgcgcggcaccaacttcccctccgacggccccgtaatgcagaagaa
gaccatgggctgggaggcctcctccgagcggatgtaccccgaggacggcgccctgaagggcgagatcaagcagaggctgaagctga
aggacggcggccactacgacgctgaggtcaagaccacctacaaggccaagaagcccgtgcagctgcccggcgcctacaacgtcaac
atcaagttggacatcacctcccacaacgaggactacaccatcgtggaacagtacgaacgcgccgagggccgccactccaccggcggc
atggacgagctgtacaagtgagcggccgctcgagtctagagggcccgggtaaggagggcccgtttaaacccgctgatcagcctcgact
gtgccttctagttgccagccatctgttgtttgcccctcccccgtgccttccttgaccctggaaggtgccactcccactgtcctttcctaata
aaatgaggaaattgcatcgcattgtctgagtaggtgtcattctattctggggggtggggtggggcaggacagcaagggggaggattg
ggaagacaatagcaggcatgctggggatgcggtgggctctatggctcgctttcttgctgtccaatttctattaaaggttcctttgttccct
aagtccaactactaaactggggatgcggccgctcgagtctagagatccggtgtggaaagtccccaggctccccagcaggcagaagta
tgcaaagcatgcatctcaattagtcagcaaccaaagctctagagatccggtgtggaaagtccccaggctccccagcaggcagaagtat

gcaaagcatgcatctcaattagtcagcaaccaaagctttaaaccatccggtgtggaaagtccccaggctccccagcaggcagaagtat

gcaaagcatgcatctcaattagtcagcaaccaaagctttaaacccgctgatcagcctcgactacaacaaggcaaggcttgaccgacaa

ttgcatgaagaatctgcttagggttaggcgttttgcgctgctttgtgacattaagcgcggcgggtgtggtggttacgcgcagcgtgacc

gctacacttgccagcgccctagcgcccgctcctttcgctttcttcccttcctttctcgccacgttcgccggctttccccgtcaagctctaa

atcgggggctccctttagggttccgatttagtgctttacggcacctcgaccccaaaaaacttgattagggtgatggttcacgtagtgggc

catcgccctgatagacggtttttcgccctttgacgttggagtccacgttctttaatagtggactcttgttccaaactggaacaacactcaa

ccctatctcggtctattcttttgatttataagggattttgccgatttcggcctattggttaaaaaatgagctgatttaacaaaaatttaacg

cgaattttaacaaaatattaacgcttacaatttaggtggcacttttcggggaaatgtgcgcggaacccctatttgtttatttttctaaatac

attcaaatatgtatccgctcatgagacaataaccctgataaatgcttcaataatattgaaaaaggaagagtatgagtattcaacatttcc

gtgtcgcccttattcccttttttgcggcattttgccttcctgtttttgctcacccagaaacgctggtgaaagtaaaagatgctgaagatca

gttgggtgcacgagtgggttacatcgaactggatctcaacagcggtaagatccttgagattttcgccccgaagaacgttttccaatgat

gagcacttttaaagttctgctatgtggcgcggtattatcccgtattgacgccgggcaagagcaactcggtcgccgcatacactattctca

gaatgacttggttgagtactcaccagtcacagaaaagcatcttacggatggcatgacagtaagagaattatgcagtgctgccataacc

atgagtgataacactgcggccaacttacttctgacaacgatcggaggaccgaaggagctaaccgcttttttgcacaacatgggggatc

atgtaactcgccttgatcgttgggaaccggagctgaatgaagccataccaaacgacgagcgtgacaccacgatgcctgtagcaatggc

aacaacgttgcgcaaactattaactggcgaactacttactctagcttcccggcaacaattaatagactggatggaggcggataaagttg

caggaccacttctgcgctcggcccttccggctggctggtttattgctgataaatctggagccggtgagcgtgggtctcgcggtatcattg

cagcactggggccagatggtaagccctcccgtatcgtagttatctacacgacggggagtcaggcaactatggatgaacgaaatagac

agatcgctgagataggtgcctcactgattaagcattggtaactgtcagaccaagtttactcatatatactttagattgatttaaaacttca

tttttaatttaaaaggatctaggtgaagatcctttttgataatctcatgaccaaaatcccttaacgtgagttttcgttccactgagcgtcag

accccgtagaaaagatcaaaggatcttcttgagatcctttttttctgcgcgtaatctgctgcttgcaaacaaaaaaaccaccgctaccag

cggtggtttgtttgccggatcaagagctaccaactctttttccgaaggtaactggcttcagcagagcgcagataccaaatactgtccttc

tagtgtagccgtagttaggccaccacttcaagaactctgtagcaccgcctacatacctcgctctgctaatcctgttaccagtggctgctg

ccagtggcgataagtcgtgtcttaccgggttggactcaagacgatagttaccggataaggcgcagcggtcgggctgaacggggggtt

cgtgcacacagcccagcttggagcgaacgacctacaccgaactgagatacctacagcgtgagctatgagaaagcgccacgcttcccg

aagggagaaaggcggacaggtatccggtaagcggcagggtcggaacaggagagcgcacgagggagcttccagggggaaacgcct

ggtatctttatagtcctgtcgggtttcgccacctctgacttgagcgtcgatttttgtgatgctcgtcaggggggcggagcctatggaaaa acgccagcaacgcggcctttttacggttcctggccttttgctggccttttgctcactcatt

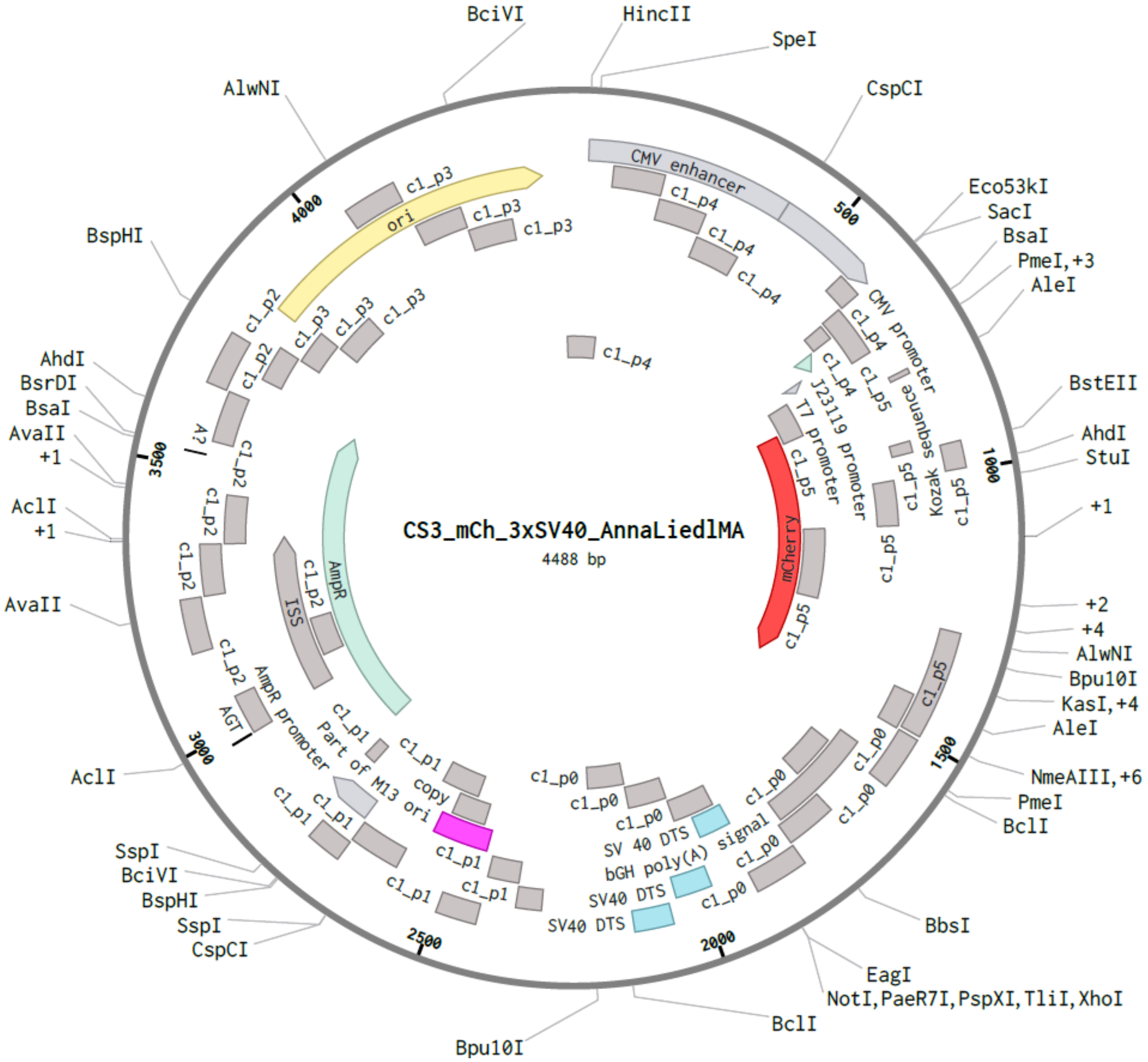


Figure S15: Plasmid used for transfection experiments. The fluorescent protein mCherry is expressed as reporter. The plasmid was a kind gift from the Dietz lab engineered by Anna Liedl, already published in this reference (*1*).

#### The full sequence of the plasmid that was used to create the stable cell line HEK293T that expresses mScarlet-I upon doxycycline induction is provided below

gtggcacttttcggggaaatgtgcgcggaacccctatttgtttatttttctaaatacattcaaatatgtatccgctcatgagacaataacc
ctgataaatgcttcaataatattgaaaaaggaagagtatgagtattcaacatttccgtgtcgcccttattcccttttttgcggcattttgcc
ttcctgtttttgctcacccagaaacgctggtgaaagtaaaagatgctgaagatcagttgggtgcacgagtgggttacatcgaactggat
ctcaacagcggtaagatccttgagagttttcgccccgaagaacgttttccaatgatgagcacttttaaagttctgctatgtggcgcggta
ttatcccgtattgacgccgggcaagagcaactcggtcgccgcatacactattctcagaatgacttggttgagtactcaccagtcacaga
aaagcatcttacggatggcatgacagtaagagaattatgcagtgctgccataaccatgagtgataacactgcggccaacttacttctga
caacgatcggaggaccgaaggagctaaccgcttttttgcacaacatgggggatcatgtaactcgccttgatcgttgggaaccggagct
gaatgaagccataccaaacgacgagcgtgacaccacgatgcctgtagcaatggcaacaacgttgcgcaaactattaactggcgaact
acttactctagcttcccggcaacaattaatagactggatggaggcggataaagttgcaggaccacttctgcgctcggcccttccggctg
gctggtttattgctgataaatctggagccggtgagcgtggatctcgcggtatcattgcagcactggggccagatggtaagccctcccgt
atcgtagttatctacacgacggggagtcaggcaactatggatgaacgaaatagacagatcgctgagataggtgcctcactgattaagc
attggtaactgtcagaccaagtttactcatatatactttagattgatttaaaacttcatttttaatttaaaaggatctaggtgaagatcctt
tttgataatctcatgaccaaaatcccttaacgtgagttttcgttccactgagcgtcagaccccgtagaaaagatcaaaggatcttcttga
gatcctttttttctgcgcgtaatctgctgcttgcaaacaaaaaaaccaccgctaccagcggtggtttgtttgccggatcaagagctacca
actctttttccgaaggtaactggcttcagcagagcgcagataccaaatactgtccttctagtgtagccgtagttaggccaccacttcaag
aactctgtagcaccgcctacatacctcgctctgctaatcctgttaccagtggctgctgccagtggcgataagtcgtgtcttaccgggttg
gactcaagacgatagttaccggataaggcgcagcggtcgggctgaacggggggttcgtgcacacagcccagcttggagcgaacgac
ctacaccgaactgagatacctacagcgtgagctatgagaaagcgccacgcttcccgaagggagaaaggcggacaggtatccggtaa
gcggcagggtcggaacaggagagcgcacgagggagcttccagggggaaacgcctggtatctttatagtcctgtcgggtttcgccacc
tctgacttgagcgtcgatttttgtgatgctcgtcaggggggcggagcctatggaaaaacgccagcaacgcggcctttttacggttcctg
gccttttgctggccttttgctcacatgttctttcctgcgttatcccctgattctgtggataaccgtattaccgcctttgagtgagctgatacc
gctcgccgcagccgaacgaccgagcgcagcgagtcagtgagcgaggaagcggaagagcgcccaatacgcaaaccgcctctccccg
cgcgttggccgattcattaatgcagctggcacgacaggtttcccgactggaaagcgggcagtgagcgcaacgcaattaatgtgagtta

gctcactcattaggcaccccaggctttacactttatgcttccggctcgtatgttgtgtggaattgtgagcggataacaatttcacacagg
aaacagctatgaccatgattacgccaagcgcgtgtatacttaaccctagaaagatagtctgcgtaaaattgacgcatgcattcttgaaa
tattgctctctctttctaaatagcgcgaatccgtcgctgtgcatttaggacatctcagtcgccgcttggagctcccgtgaggcgtgcttgt
caatgcggtaagtgtcactgattttgaactataacgaccgcgtgagtcaaaatgacgcatgattatcttttacgtgacttttaagatttaa
ctcatacgataattatattgttatttcatgttctacttacgtgataacttattatatatatattttcttgttatagatatcgtgactaatatat
aataaaggccggccgccctgcaggtttaaatgatttgccctcccatatgtccttccgagtgagagacacaaaaaattccaacacactat
tgcaatgaaaatacatttcctttattagccagaagtcagatgcaggcccaaggttttgcttttttttttttaagaaaggccaaaaagcaa
aacctgagactttcgcctcaggaaaagaaaaaccttccggcaagaagcatggccaccgaggctccagcgtcgactcacccggggagc
atgtcaaggtcaaaatcgtcaagagcgtcagcaggcagcatatcaaggtcaaagtcgtcaagggcatcggctgggagcatgtctaag
tcaaaatcgtcaagggcgtcggtcggcccgccgctttcgcactttagctgtttctccaggccacatatgattagttccaggccgaaaag
gaaggcaggttcggctccctgccggtcgaacagctcaattgcttgtttcagaagtgggggcatagaatcggtggtaggtgtctctcttt
cctcttttgctacttgatgctcctgttcctccaatacgcagcccagtgtaaagtggcccacggcggacagagcgtacagtgcgttctcca
gggagaagccttgctgacacaggaacgcgagctgattttccagggtttcgtactgtttctctgttgggcgggtgccgagatgcacttta
gccccgtcgcgatgtgagaggagagcacagcggtatgacttggcgttgttccgcagaaagtcttgccatgactcgccttccagggggc
aggagtgggtatgatgcctgtccagcatctcgattggcagggcatcgagcagggcccgcttgttcttcacgtgccagtacagggtagg
ctgctcaactcccagcttttgagcgagtttccttgtcgtcaggccttcgataccgactccattgagtaattccagagcagagtttatgact
ttgctcttgtccagtctagacatggtggcggcccgggcacagctggggagagaggtcggtgattcggtcaacgagggagccgactgc
cgacgtgcgctccggaggcttgcagaatgcggaacaccgcgcgggcaggaacagggcccacactaccgccccacaccccgcctccc
gcaccgccccttcccggccgctgctctcggcacgccctgctgagcagccgctattggccacagcccatcgcggtcggcgcgctgccat
tgctccctggcgctgtccgtctgcgagggtactagtgagacgtgcggcttccgtttgtcacgtccggcacgccgcgaaccgcaaggaa
ccttcccgacttaggggcggagcaggaagcgtcgccgggggggcccacaagggtagcggcgaagatccgggtgacgctgcgaacgg
acgtgaagaatgtgcgagacccagggtcggcgccgctgcgtttcccggaaccacgcccagagcagccgcgtccctgcgcaaacccag
ggctgccttggaaaaggcgcaactccaaccccgtggcggccgcgatcgaattccgtctcacgcgccggattcgacattgattattgact
agttattaatagtaatcaattacggggtcattagttcatagcccatatatggagttccgcgttacataacttacggtaaatggcccgcct
ggctgaccgcccaacgacccccgcccattgacgtcaataatgacgtatgttcccatagtaacgccaatagggactttccattgacgtca
atgggtggagtatttacggtaaactgcccacttggcagtacatcaagtgtatcatatgccaagtacgccccctattgacgtcaatgacg
gtaaatggcccgcctggcattatgcccagtacatgaccttatgggactttcctacttggcagtacatctacgtattagtcatcgctattac

catggtcgaggtgagccccacgttctgcttcactctccccatctcccccccctccccacccccaattttgtatttatttattttttaattattt
tgtgcagcgatgggggcgggggggggggggggggcgcgcgccaggcggggcggggcggggcgaggggcggggcggggcgaggc
ggagaggtgcggcggcagccaatcagagcggcgcgctccgaaagtttccttttatggcgaggcggcggcggcggcggccctataaa
aagcgaagcgcgcggcgggcgggagtcgctgcgcgctgccttcgccccgtgccccgctccgccgccgcctcgcgccgcccgccccgg
ctctgactgaccgcgttactcccacaggtgagcgggcgggacggcccttctcctccgggctgtaattagcgcttggtttaatgacggct
tgtttcttttctgtggctgcgtgaaagccttgaggggctccgggagggccctttgtgcggggggagcggctcggggggtgcgtgcgtg
tgtgtgtgcgtggggagcgccgcgtgcggctccgcgctgcccggcggctgtgagcgctgcgggcgcggcgcggggctttgtgcgctc
cgcagtgtgcgcgaggggagcgcggccgggggcggtgccccgcggtgcggggggggctgcgaggggaacaaaggctgcgtgcgg
ggtgtgtgcgtgggggggtgagcagggggtgtgggcgcgtcggtcgggctgcaaccccccctgcacccccctccccgagttgctgag
cacggcccggcttcgggtgcggggctccgtacggggcgtggcgcggggctcgccgtgccgggcggggggtggcggcaggtggggg
tgccgggcggggcggggccgcctcgggccggggagggctcgggggaggggcgcggcggcccccggagcgccggcggctgtcgag
gcgcggcgagccgcagccattgccttttatggtaatcgtgcgagagggcgcagggacttcctttgtcccaaatctgtgcggagccgaa
atctgggaggcgccgccgcaccccctctagcgggcgcggggcgaagcggtgcggcgccggcaggaaggaaatgggcggggaggg
ccttcgtgcgtcgccgcgccgccgtccccttctccctctccagcctcggggctgtccgcggggggacggctgccttcgggggggacgg
ggcagggcggggttcggcttctggcgtgtgaccggcggctctagagcctctgctaaccatgttcatgccttcttctttttcctacagatc
cttaattaataatacgactcactataggggccgccaccatggtgagcaagggcgaggagctgttcaccggggtggtgcccatcctggt
cgagctggacggcgacgtaaacggccacaagttcagcgtccgcggcgagggcgagggcgatgccaccaacggcaagctgaccctga
agttcatctgcaccaccggcaagctgcccgtgccctggcccaccctcgtgaccaccttaggctacggcgtggcctgcttcgcccgctac
cccgaccacatgaagcagcacgacttcttcaagtccgccatgcccgaaggctacgtccaggagcgcaccatctctttcaaggacgacg
gcacctacaagacccgcgccgaggtgaagttcgagggcgacaccctggtgaaccgcatcgtgctgaagggcatcgacttcaaggag
gacggcaacatcctggggcacaagctggagtacaacttcaacagccacaaggtctatatcacggccgacaagcagaagaacggcatc
aaggctaacttcaagacccgccacaacgttgaggacggcggcgtgcagctcgccgaccactaccagcagaacacccccatcggcgac
ggccccgtgctgctgcccgacaaccactacctgagccatcagtccaaactgagcaaagaccccaacgagaagcgcgatcacatggtc
ctgaaggagagggtgaccgccgccgggattacacatgacatggacgagctctacaaatgaacgcgtcaagcacgcagcaatgcagc
tcaaaacgcttagcctagccacacccccacgggaaacagcagtgattaacctttagcaatatacgaaagtttaactaagctatactaac
cccagggttggtcaatttcgtgccagccacaccgtggatgcgcacccccctctccctcccccccccctaacgttactggccgaagccgc
ttggaataaggccggtgtgcgtttgtctatatgttattttccaccatattgccgtcttttggcaatgtgagggcccggaaacctggccctg

tcttcttgacgagcattcctaggggtctttcccctctcgccaaaggaatgcaaggtctgttgaatgtcgtgaaggaagcagttcctctgg

aagcttcttgaagacaaacaacgtctgtagcgaccctttgcaggcagcggaaccccccacctggcgacaggtgcctctgcggccaaaa

gccacgtgtataagatacacctgcaaaggcggcacaaccccagtgccacgttgtgagttggatagttgtggaaagagtcaaatggctc

tcctcaagcgtattcaacaaggggctgaaggatgcccagaaggtaccccattgtatgggatctgatctggggcctcggtgcacatgctt

tacatgtgtttagtcgaggttaaaaaacgtctaggccccccgaaccacggggacgtggttttcctttgaaaaacacgatgataatatgg

ccaccaccatgaccgagtacaagcctacagtgcggctggctaccagggacgatgtgccaagagctgtgcggacactggccgctgcct

tcgccgattaccctgccacaagacacaccgtggaccccgaccggcacatcgagagagtgaccgagctgcaagaactgtttctgactag

agtgggcctggacatcggcaaagtgtgggtggccgatgatggcgccgctgtggctgtgtggacaacccctgagtctgtggaagcagg

cgctgtgttcgccgagatcggacctagaatggccgagctgagcggctccagactggctgcccagcagcagatggaaggcctgctggc

cccccacagaccaaaagagcctgcctggtttctggctaccgtgggcgtgtcacctgaccaccagggcaagggactgggatctgctgtg

gtgctgcctggggtggaagctgctgaaagggctggcgtgcccgccttcctggaaacaagcgcccccagaaacctgcccttctacgag

agactgggcttcaccgtgaccgccgacgtggaagtgcctgagggccctagaacctggtgcatgaccagaaagcctggcgccggttcc

ggagctacaaacttcagcctgctgaaacaggctggcgacgtggaagagaaccccggtcctgcttcttatccttgtcaccaacacgcca

gcgccttcgatcaagccgctagaagcagaggccacagcaacagaagaacagccctcaggccaagaaggcagcaagaggctacaga

agtgcggctggaacagaagatgcccacactgctgagagtgtatatcgacggccctcatggcatgggcaagaccacaacaacacagct

gctggtggctctgggcagcagagatgacatcgtgtacgtgcccgagcctatgacctactggcaggtcctgggagcctctgagacaatc

gccaacatctacaccacacagcacagactggaccagggcgaaattagcgcaggcgacgctgctgtggtcatgacatctgcccagatc

acaatgggcatgccttacgccgtgacagacgctgtgctggctccacatattggcggcgaggctggatcttctcacgctccaccacctgc

tctgaccctgatcttcgacagacaccccatcttcgccctgctgtgttaccctgccgctcggtatctgatgggcagcatgacacctcaggc

cgtgctggctttcgtggctctgattcctcctacactgcccggcacaaacatcgtgcttggagccctgccagaggacagacacatcgaca

gactggctaagagacagaggcctggcgagagactggatctggctatgctggccgccatcagaagagtgtacggcctgctggctaaca

ccgtgcgctatcttcaaggcggcggttcctggagagaggactggggacagctctctggcacagcagttcctccacaaggcgctgagcc

tcagtctaacgctggacccagacctcacatcggcgacaccctgttcacactgttccgcgcccctgaactgctggcccctaacggcgacc

tgtataacgtgttcgcctgggctctcgacgtgctggcaaaaagactgcggcccatgcatgtgttcatcctggactacgatcagagccca

gccggatgcagagatgccctgctgcaactgacaagcggcatggtgcagacccatgtgacaacccctggcagcatccccaccatctgtg

acctcgccagaaccttcgctagagagatgggcgaagccaactaagtttaaacgctcgctttcttgctgtccaatttctattaaaggttcc

tttgttccctaagtccaactactaaactgggggatattatgaagggccttgagcatctggattctgcctaatataaaacatttattttcatt

gcgacgatcgtcagacatgataagatacattgatgagtttggacaaaccacaactagaatgcagtgaaaaaaatgctttatttgtgaaa
tttgtgatgctattgctttatttgtaaccattataagctgcaataaacaagttaacaacaacaattgcattcattttatgtttcaggttcag
ggggaggtgtgggaggttttttaaagcaagtaaaacctctacaaatgtggtaggcgcgtctacttgtacagctcgtccattccgcctgt
gctgtgtctcccctcgcttctctcgtactgttccaccacggtgtagtcctcgttgtggctggtgatgtccagctttctgtccacgttgtagg
cgccaggcatctgcacaggtttcttggccttgtaggtggtcttgaagtcggccaggtatctgccgccatccttcagtctcagggccatct
tgatgtcgcccttcagcacgccatcttcagggtacagtctctcggtgctggcttcccagcccattgtctttttctgcatcacagggccgtc
tggagggaagtttgtgccccgcagcttcactttgtagatcagggtcccatcttccagagatgtgtcctgtgtcacagtcacggctccgcc
gtcctcgaagttcatcactctctcccacttgaagccctctgggaaagactgcttgtagtagtcggggatgtcagcggggtgcttgatga
aggccctgctgccgtacataaactgtggagacaggatgtcccagctgaaaggcagagggccgcctttggtcactttcagcttggcggt
ctgagttccctcgtaaggtctgccctcgccttcgccttcgatctcgaactcgtggccgttcatgctgccttccatgtgcaccttgaacctc
atgaactctttgatcacggcctcgcctttagaaaccatggtggcggcccctatagtgagtcgtattattaatcgcgtataagacaaaagt
gttgtggaattgctccaggcgatctgacggttcactaaacgagctctgcttttataggcgcccaccgtacacgcctaaagcttatacgtt
ctctatcactgatagggagtaaactggatatacgttctctatcactgatagggagtaaactgtagatacgttctctatcactgataggga
gtaaactggtcatacgttctctatcactgatagggagtaaactccttatacgttctctatcactgatagggagtaaagtctgcatacgttc
tctatcactgatagggagtaaactcttcatacgttctctatcactgatagggagtaaactcgaggaatacttgaagtcgaaagaagaga
aatgttctggcacctgcacttgcactggggacagcctattttgctagtttgttttgtttcgttttgttttgatggagagcgtatgttaatcg
attcacacaaaaaaccaacacactattgcaatgaaaataaatttcctttatttaaatggccggcctaaaagttttgttactttatagaaga
aattttgagtttttgtttttttttaataaataaataaacataaataaattgtttgttgaatttattattagtatgtaagtgtaaatataataaa
acttaatatctattcaaattaataaataaacctcgatatacagaccgataaaacacatgcgtcaattttacgcatgattatctttaacgta
cgtcacaatatgattatctttctagggttaagtatacacgcgctcactggccgtcgttttacaacgtcgtgactgggaaaaccctggcgt
tacccaacttaatcgccttgcagcacatccccctttcgccagctggcgtaatagcgaagaggcccgcaccgatcgcccttcccaacagt
tgcgcagcctgaatggcgaatgggacgcgccctgtagcggcgcattaagcgcggcgggtgtggtggttacgcgcagcgtgaccgcta
cacttgccagcgccctagcgcccgctcctttcgctttcttcccttcctttctcgccacgttcgccggctttccccgtcaagctctaaatcgg
gggctccctttagggttccgatttagtgctttacggcacctcgaccccaaaaaacttgattagggtgatggttcacgtagtgggccatcg
ccctgatagacggtttttcgccctttgacgttggagtccacgttctttaatagtggactcttgttccaaactggaacaacactcaacccta
tctcggtctattcttttgatttataagggattttgccgatttcggcctattggttaaaaaatgagctgatttaacaaaaatttaacgcgaat
tttaacaaaatattaacgcttacaatttag

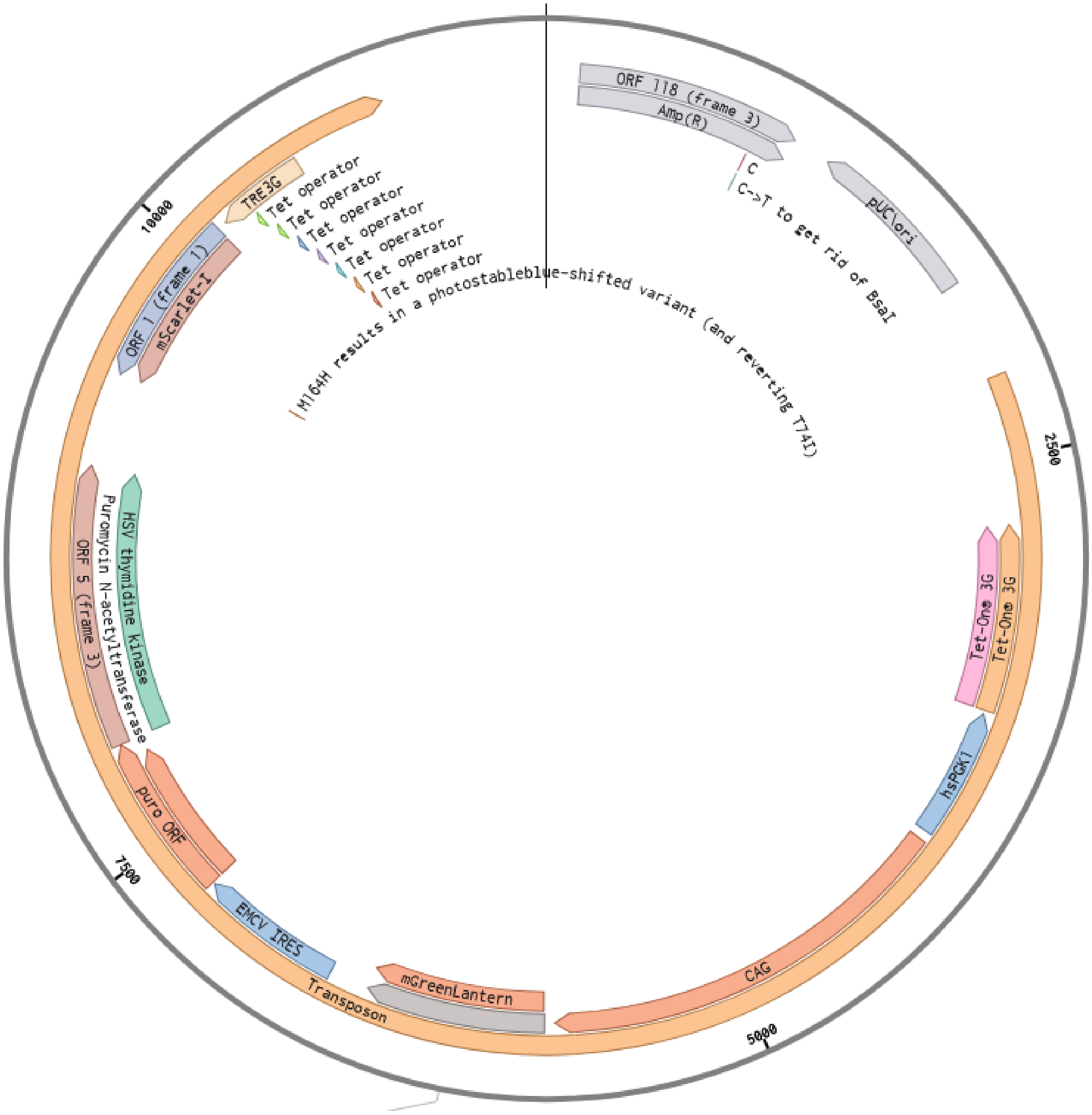


Figure S16: Transposon used for engineering the stable cell line HEK293T which expresses mScarlet-I upon doxycycline addition.

#### The full sequence of the plasmid encoding the transposase is provided below

ggcgcgcctggtacccgttgaattctaccgggtaggggaggcgcttttcccaaggcagtctggagcatgcgctttagcagccccgctg
ggcacttggcgctacacaagtggcctctggcctcgcacacattccacatccaccggtaggcgccaaccggctccgttctttggtggccc
cttcgcgccaccttctactcctcccctagtcaggaagttcccccccgccccgcagctcgcgtcgtgcaggacgtgacaaatggaagtag
cacgtctcactagtctcgtgcagatggacagcaccgctgagcaatggaagcgggtaggcctttggggcagcggccaatagcagctttg
ctccttcgctttctgggctcagaggctgggaaggggtgggtccgggggcgggctcaggggcgggctcaggggcggggcgggcgccc
gaaggtcctccggaggcccggcattctgcacgcttcaaaagcgcacgtctgccgcgctgttctcctcttcctcatctccgggcctttcga
ccaccggtgatccttaattaataatacgactcactataggggccgccaccatgggcagcagcctggacgacgagcacatcctgagcgc
cctgctgcagagcgacgacgagctggtcggcgaggacagcgacagcgagatcagcgaccacgtgagcgaggacgacgtgcagtcc
gacaccgaggaggccttcatcgacgaggtgcacgaggtgcagcctaccagcagcggctccgagatcctggacgagcagaacgtgat
cgagcagcccggcagctccctggccagcaacaggatcctgaccctgccccagaggaccatcaggggcaagaacaagcactgctggt
ccacctccaagagcaccaggcggagcagggtgtccgccctgaacatcgtgagaagccagaggggccccaccaggatgtgcaggaac
atctacgacccccctgctgtgcttcaagctgttcttcaccgacgagatcatcagcgagatcgtgaagtggaccaacgccgagatcagcct
gaagaggcgggagagcatgaccggcgccaccttcagggacaccaacgaggacgagatctacgccttcttcggcatcctggtgatgac
cgccgtgaggaaggacaaccacatgagcaccgacgacctgttcgacagatccctgagcatggtgtacgtgagcgtgatgagcaggg
acagattcgacttcctgatcagatgcctgaggatggacgacaagagcatcaggcccaccctgcgggagaacgacgtgttcacccccgt
gagaaagatctgggacctgttcatccaccagtgcatccagaactacacccctggcgcccacctgaccatcgacgagcagctgctgggc
ttcaggggcaggtgccccttcaggatgtatatccccaacaagcccagcaagtacggcatcaagatcctgatgatgtgcgacagcggca
ccaagtacatgatcaacggcatgccctacctgggcaggggcacccagaccaacggcgtgcccctgggcgagtactacgtgaaggagc
tgtccaagcccgtccacggcagctgcagaaacatcacctgcgacaactggttcaccagcatcccccctggccaagaacctgctgcagga
gccctacaagctgaccatcgtgggcaccgtgagaagcaacaagagagagatccccgaggtcctgaagaacagcaggtccaggcccg
tgggcaccagcatgttctgcttcgacggcccccctgaccctggtgtcctacaagcccaagcccgccaagatggtgtacctgctgtccagc
tgcgacgaggacgccagcatcaacgagagcaccggcaagccccagatggtgatgtactacaaccagaccaagggcggcgtggacac
cctggaccagatgtgcagcgtgatgacctgcagcagaaagaccaacaggtggcccatggccctgctgtacggcatgatcaacatcgc
ctgcatcaacagcttcatcatctacagccacaacgtgagcagcaagggcgagaaggtgcagagccggaaaaagttcatgcggaacct
gtacatgagcctgacctccagcttcatgaggaagaggctggaggcccccaccctgaagagatacctgagggacaacatcagcaacat

cctgcccaacgaggtgcccggcaccagcgacgacagcaccgaggagcccgtgatgaagaagaggacctactgcacctactgtccca
gcaagatcagaagaaaggccaacgccagctgcaagaagtgtaagaaggtcatctgccgggagcacaacatcgacatgtgccagagc
tgtttctgaacgcgtaaatgattgcagatccactagttctagagctcgctgatcagcctcgactgtgccttctagttgccagccatctgtt
gtttgcccctcccccgtgccttccttgaccctggaaggtgccactcccactgtcctttcctaataaaatgaggaaattgcatcgcattgtc
tgagtaggtgtcattctattctggggggtggggtggggcaggacagcaagggggaggattgggaagagaatagcaggcatgctggg
gatgcggtgggctctatggcttctgaggcggaaagaaccagctggggcgccactggccgtcgttttacaacgtcgtgactgggaaaac
cctggcgttacccaacttaatcgccttgcagcacatccccctttcgccagctggcgtaatagcgaagaggcccgcaccgatcgcccttc
ccaacagttgcgcagcctgaatggcgaatgggacgcgccctgtagcggcgcattaagcgcggcgggtgtggtggttacgcgcagcgt
gaccgctacacttgccagcgccctagcgcccgctcctttcgctttcttcccttcctttctcgccacgttcgccggctttccccgtcaagctc
taaatcgggggctccctttagggttccgatttagtgctttacggcacctcgaccccaaaaaacttgattagggtgatggttcacgtagtg
ggccatcgccctgatagacggtttttcgccctttgacgttggagtccacgttctttaatagtggactcttgttccaaactggaacaacact
caaccctatctcggtctattcttttgatttataagggattttgccgatttcggcctattggttaaaaaatgagctgatttaacaaaaattta
acgcgaattttaacaaaatattaacgcttacaatttaggtggcacttttcggggaaatgtgcgcggaacccctatttgtttatttttctaa
atacattcaaatatgtatccgctcatgagacaataaccctgataaatgcttcaataatattgaaaaaggaagagtatgagtattcaacat
ttccgtgtcgcccttattcccttttttgcggcattttgccttcctgtttttgctcacccagaaacgctggtgaaagtaaaagatgctgaaga
tcagttgggtgcacgagtgggttacatcgaactggatctcaacagcggtaagatccttgagagttttcgccccgaagaacgttttccaa
tgatgagcacttttaaagttctgctatgtggcgcggtattatcccgtattgacgccgggcaagagcaactcggtcgccgcatacactatt
ctcagaatgacttggttgagtactcaccagtcacagaaaagcatcttacggatggcatgacagtaagagaattatgcagtgctgccata
accatgagtgataacactgcggccaacttacttctgacaacgatcggaggaccgaaggagctaaccgcttttttgcacaacatggggg
atcatgtaactcgccttgatcgttgggaaccggagctgaatgaagccataccaaacgacgagcgtgacaccacgatgcctgtagcaat
ggcaacaacgttgcgcaaactattaactggcgaactacttactctagcttcccggcaacaattaatagactggatggaggcggataaa
gttgcaggaccacttctgcgctcggcccttccggctggctggtttattgctgataaatctggagccggtgagcgtgggtctcgcggtatc
attgcagcactggggccagatggtaagccctcccgtatcgtagttatctacacgacggggagtcaggcaactatggatgaacgaaata
gacagatcgctgagataggtgcctcactgattaagcattggtaactgtcagaccaagtttactcatatatactttagattgatttaaaac
ttcatttttaatttaaaaggatctaggtgaagatcctttttgataatctcatgaccaaaatcccttaacgtgagttttcgttccactgagcg
tcagaccccgtagaaaagatcaaaggatcttcttgagatcctttttttctgcgcgtaatctgctgcttgcaaacaaaaaaaccaccgcta
ccagcggtggtttgtttgccggatcaagagctaccaactctttttccgaaggtaactggcttcagcagagcgcagataccaaatactgt

ccttctagtgtagccgtagttaggccaccacttcaagaactctgtagcaccgcctacatacctcgctctgctaatcctgttaccagtggct gctgccagtggcgataagtcgtgtcttaccgggttggactcaagacgatagttaccggataaggcgcagcggtcgggctgaacgggg ggttcgtgcacacagcccagcttggagcgaacgacctacaccgaactgagatacctacagcgtgagctatgagaaagcgccacgctt cccgaagggagaaaggcggacaggtatccggtaagcggcagggtcggaacaggagagcgcacgagggagcttccagggggaaac gcctggtatctttatagtcctgtcgggtttcgccacctctgacttgagcgtcgatttttgtgatgctcgtcaggggggcggagcctatgg aaaaacgccagcaacgcggcctttttacggttcctggccttttgctggccttttgctcacatgttctttcctgcgttatcccctgattctgt ggataaccgtattaccgcctttgagtgagctgataccgctcgccgcagccgaacgaccgagcgcagcgagtcagtgagcgaggaagc ggaagagcgcccaatacgcaaaccgcctctccccgcgcgttggccgattcattaatgcagctggcacgacaggtttcccgactggaaa gcgggcagtgagcgcaacgcaattaatgtgagttagctcactcattaggcaccccaggctttacactttatgcttccggctcgtatgttg tgtggaattgtgagcggataacaatttcacacaggaaacagctatgaccatga

#### The full sequence of the plasmid encoding the pegRNA is provided below

ataaaacctgcaggcatgcaagcgatcgcggggccgccccccttcaccgagggcctatttcccatgattccttcatatttgcatatacgat acaaggctgttagagagataattggaattaatttgactgtaaacacaaagatattagtacaaaatacgtgacgtagaaagtaataattt cttgggtagtttgcagttttaaaattatgttttaaaatggactatcatatgcttaccgtaacttgaaagtatttcgatttcttggctttatat atcttgtggaaaggacgaaacaccgggcgaagcaggccacgccggtttcagagccaccagaagatatggcttcggtggcaagttgaa ataaggctagtccgttatcaacttgaaaaagtggcaccgagtcggtgctgtgccctggcccaccttggtcactacactcggatatggcg tggcctgcttccgcggttctatctagttacgcgttaaaccaactagaatttttttactagttctagagcggccccaattcgccctatagtga gtcgtattacgcgcgctcactggccgtcgttttacaacgtcgtgactgggaaaaccctggcgttacccaacttaatcgccttgcagcac atccccctttcgccagctggcgtaatagcgaagaggcccgcaccgatcgcccttcccaacagttgcgcagcctgaatggcgaatggga cgcgccctgtagcggcgcattaagcgcggcgggtgtggtggttacgcgcagcgtgaccgctacacttgccagcgccctagcgcccgct cctttcgctttcttcccttcctttctcgccacgttcgccggctttccccgtcaagctctaaatcgggggctccctttagggttccgatttagt gctttacggcacctcgaccccaaaaaacttgattagggtgatggttcacgtagtgggccatcgccctgatagacggtttttcgccctttg acgttggagtccacgttctttaatagtggactcttgttccaaactggaacaacactcaaccctatctcggtctattcttttgatttataagg gattttgccgatttcggcctattggttaaaaaatgagctgatttaacaaaaatttaacgcgaattttaacaaaatattaacgcttacaatt taggtggcacttttcggggaaatgtgcgcggaacccctatttgtttatttttctaaatacattcaaatatgtatccgctcatgagacaata accctgataaatgcttcaataatattgaaaaaggaagagtatgagtattcaacatttccgtgtcgcccttattcccttttttgcggcatttt

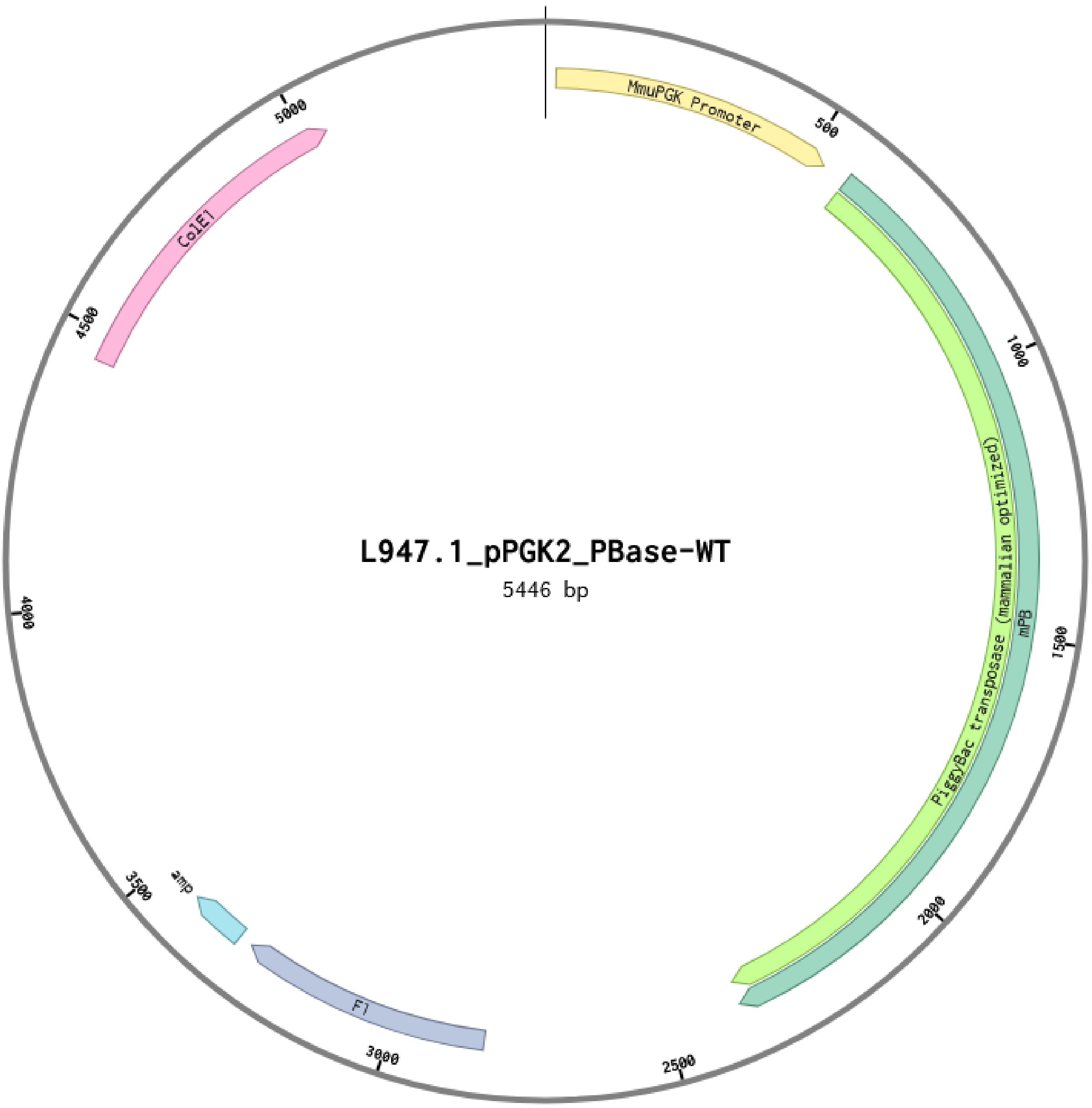


Figure S17: Transposase used for engineering the stable cell line HEK293T which expresses mScarlet-I upon doxycycline addition.

gccttcctgtttttgctcacccagaaacgctggtgaaagtaaaagatgctgaagatcagttgggtgcacgagtgggttacatcgaactg
gatctcaacagcggtaagatccttgagagttttcgccccgaagaacgttttccaatgatgagcacttttaaagttctgctatgtggcgcg
gtattatcccgtattgacgccgggcaagagcaactcggtcgccgcatacactattctcagaatgacttggttgagtactcaccagtcac
agaaaagcatcttacggatggcatgacagtaagagaattatgcagtgctgccataaccatgagtgataacactgcggccaacttactt
ctgacaacgatcggaggaccgaaggagctaaccgcttttttgcacaacatgggggatcatgtaactcgccttgatcgttgggaaccgg
agctgaatgaagccataccaaacgacgagcgtgacaccacgatgcctgtagcaatggcaacaacgttgcgcaaactattaactggcg
aactacttactctagcttcccggcaacaattaatagactggatggaggcggataaagttgcaggaccacttctgcgctcggcccttccg
gctggctggtttattgctgataaatctggagccggtgagcgtgggtctcgcggtatcattgcagcactggggccagatggtaagccctc
ccgtatcgtagttatctacacgacggggagtcaggcaactatggatgaacgaaatagacagatcgctgagataggtgcctcactgatt
aagcattggtaactgtcagaccaagtttactcatatatactttagattgatttaaaacttcatttttaatttaaaaggatctaggtgaagat
cctttttgataatctcatgaccaaaatcccttaacgtgagttttcgttccactgagcgtcagaccccgtagaaaagatcaaaggatcttct
tgagatcctttttttctgcgcgtaatctgctgcttgcaaacaaaaaaaccaccgctaccagcggtggtttgtttgccggatcaagagcta
ccaactctttttccgaaggtaactggcttcagcagagcgcagataccaaatactgtccttctagtgtagccgtagttaggccaccacttc
aagaactctgtagcaccgcctacatacctcgctctgctaatcctgttaccagtggctgctgccagtggcgataagtcgtgtcttaccggg
ttggactcaagacgatagttaccggataaggcgcagcggtcgggctgaacggggggttcgtgcacacagcccagcttggagcgaacg
acctacaccgaactgagatacctacagcgtgagctatgagaaagcgccacgcttcccgaagggagaaaggcggacaggtatccggt
aagcggcagggtcggaacaggagagcgcacgagggagcttccagggggaaacgcctggtatctttatagtcctgtcgggtttcgcca
cctctgacttgagcgtcgatttttgtgatgctcgtcaggggggcggagcctatggaaaaacgccagcaacgcggcctttttacggttcc
tggccttttgctggccttttgctcacatgttctttcctgcgttatcccctgattctgtggataaccgtattaccgcctttgagtgagctgata
ccgctcgccgcagccgaacgaccgagcgcagcgagtcagtgagcgaggaagcggaagagcgcccaatacgcaaaccgcctctcccc
gcgcgttggccgattcattaatgcagctggcacgacaggtttcccgactggaaagcgggcagtgagcgcaacgcaattaatgtgagtt
agctcactcattaggcaccccaggctttacactttatgcttccggctcgtatgttgtgtggaattgtgagcggataacaatttcacacag
gaaacagctatgaccatgattacgccaagcgcgcaattaaccctcactaaagggaacaaaagctggaaacatgcatgaagttcctatt
ccgaagttcctattctctagaaagtataggaacttc

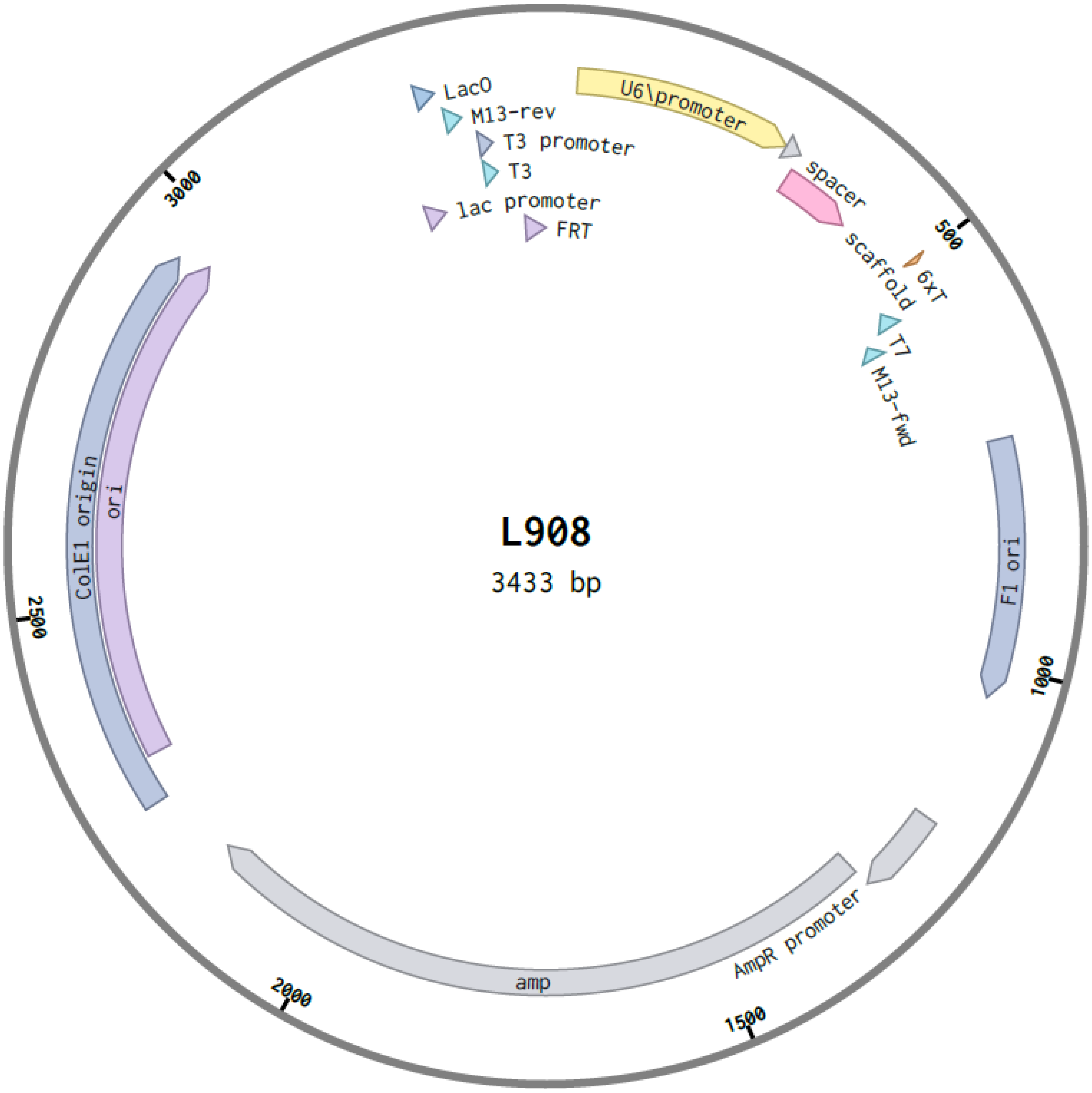


Figure S18: Plasmid used for prime editing experiments. pegRNA is expressed which leads the prime editor.

#### The full sequence of the plasmid encoding the pegRNA for RNA production is provided below

caattaatgtgagttagctcactcattaggcaccccaggctttacactttatgcttccggctcgtatgttgtgtggaattgtgagcggata
acaatttcacacaggaaacagctatgaccatgattacgccaagcgcgctaatacgactcactatagggcgaagcaggccacgccggtt
tcagagccaccagaagatatggcttcggtggcaagttgaaataaggctagtccgttatcaacttgaaaaagtggcaccgagtcggtgc
tgtgaccaccttaggctacggcgtggcctgcttccgcggttctatctagttacgcgttaaaccaactagaatttttggagaccgcgcgc
tcactggccgtcgttttacaacgtcgtgactgggaaaaccctggcgttacccaacttaatcgccttgcagcacatccccctttcgccagc
tggcgtaatagcgaagaggcccgcaccgatcgcccttcccaacagttgcgcagcctgaatggcgaatgggacgcgccctgtagcggc
gcattaagcgcggcgggtgtggtggttacgcgcagcgtgaccgctacacttgccagcgccctagcgcccgctcctttcgctttcttccct
tcctttctcgccacgttcgccggctttccccgtcaagctctaaatcgggggctccctttagggttccgatttagtgctttacggcacctcg
accccaaaaaacttgattagggtgatggttcacgtagtgggccatcgccctgatagacggtttttcgccctttgacgttggagtccacgt
tctttaatagtggactcttgttccaaactggaacaacactcaaccctatctcggtctattcttttgatttataagggattttgccgatttcg
gcctattggttaaaaaatgagctgatttaacaaaaatttaacgcgaattttaacaaaatattaacgcttacaatttaggtggcacttttcg
gggaaatgtgcgcggaacccctatttgtttatttttctaaatacattcaaatatgtatccgctcatgagacaataaccctgataaatgctt
caataatattgaaaaaggaagagtatgagtattcaacatttccgtgtcgcccttattcccttttttgcggcattttgccttcctgtttttgct
cacccagaaacgctggtgaaagtaaaagatgctgaagatcagttgggtgcacgagtgggttacatcgaactggatctcaacagcggt
aagatccttgagagttttcgccccgaagaacgttttccaatgatgagcacttttaaagttctgctatgtggcgcggtattatcccgtattg
acgccgggcaagagcaactcggtcgccgcatacactattctcagaatgacttggttgagtactcaccagtcacagaaaagcatcttac
ggatggcatgacagtaagagaattatgcagtgctgccataaccatgagtgataacactgcggccaacttacttctgacaacgatcgga
ggaccgaaggagctaaccgcttttttgcacaacatgggggatcatgtaactcgccttgatcgttgggaaccggagctgaatgaagcca
taccaaacgacgagcgtgacaccacgatgcctgtagcaatggcaacaacgttgcgcaaactattaactggcgaactacttactctagc
ttcccggcaacaattaatagactggatggaggcggataaagttgcaggaccacttctgcgctcggcccttccggctggctggtttattg
ctgataaatctggagccggtgagcgtgggtctcgcggtatcattgcagcactggggccagatggtaagccctcccgtatcgtagttatc
tacacgacggggagtcaggcaactatggatgaacgaaatagacagatcgctgagataggtgcctcactgattaagcattggtaactgt
cagaccaagtttactcatatatactttagattgatttaaaacttcatttttaatttaaaaggatctaggtgaagatcctttttgataatctca
tgaccaaaatcccttaacgtgagttttcgttccactgagcgtcagaccccgtagaaaagatcaaaggatcttcttgagatcctttttttct
gcgcgtaatctgctgcttgcaaacaaaaaaaccaccgctaccagcggtggtttgtttgccggatcaagagctaccaactctttttccga

aggtaactggcttcagcagagcgcagataccaaatactgtccttctagtgtagccgtagttaggccaccacttcaagaactctgtagca
ccgcctacatacctcgctctgctaatcctgttaccagtggctgctgccagtggcgataagtcgtgtcttaccgggttggactcaagacga
tagttaccggataaggcgcagcggtcgggctgaacggggggttcgtgcacacagcccagcttggagcgaacgacctacaccgaactg
agatacctacagcgtgagctatgagaaagcgccacgcttcccgaagggagaaaggcggacaggtatccggtaagcggcagggtcgg
aacaggagagcgcacgagggagcttccagggggaaacgcctggtatctttatagtcctgtcgggtttcgccacctctgacttgagcgt
cgatttttgtgatgctcgtcaggggggcggagcctatggaaaaacgccagcaacgcggcctttttacggttcctggccttttgctggcc
ttttgctcacatgttctttcctgcgttatcccctgattctgtggataaccgtattaccgcctttgagtgagctgataccgctcgccgcagcc
gaacgaccgagcgcagcgagtcagtgagcgaggaagcggaagagcgcccaatacgcaaaccgcctctccccgcgcgttggccgatt
cattaatgcagctggcacgacaggtttcccgactggaaagcgggcagtgagcgcaacg

#### The full sequence of the plasmid encoding the prime editor is provided below

gacattgattattgactagttattaatagtaatcaattacggggtcattagttcatagcccatatatggagttccgcgttacataacttacggtaaatggcccgcc

#### The full sequence of the plasmid encoding the prime editor for mRNA production is provided below

gacattgattattgactagttattaatagtaatcaattacggggtcattagttcatagcccatatatggagttccgcgttacataacttac
ggtaaatggcccgcctggctgaccgcccaacgacccccgcccattgacgtcaataatgacgtatgttcccatagtaacgccaataggg
actttccattgacgtcaatgggtggagtatttacggtaaactgcccacttggcagtacatcaagtgtatcatatgccaagtacgcccccct
attgacgtcaatgacggtaaatggcccgcctggcattatgcccagtacatgaccttatgggactttcctacttggcagtacatctacgta
ttagtcatcgctattaccatggtcgaggtgagccccacgttctgcttcactctccccatctcccccccctccccacccccaattttgtattt
atttattttttaattattttgtgcagcgatgggggcgggggggggggggggcgcgcgccaggcggggcggggcggggcgaggggc
ggggcggggcgaggcggagaggtgcggcggcagccaatcagagcggcgcgctccgaaagtttccttttatggcgaggcggcggcg
gcggcggccctataaaaagcgaagcgcgcggcgggcgggagtcgctgcgtcgcgccttcgccccgtgccccgctccgccgccgcctc
gcgccgcccgccccggctctgactgaccgcgttactcccacaggtgagcgggcgggacggcccttctcctccgggctgtaattagcgc
ttggtttaatgacggctcgtttcttttctgtggctgcgtgaaagccttaaagggctccgggagggccctttgtgcggggggggagcggct

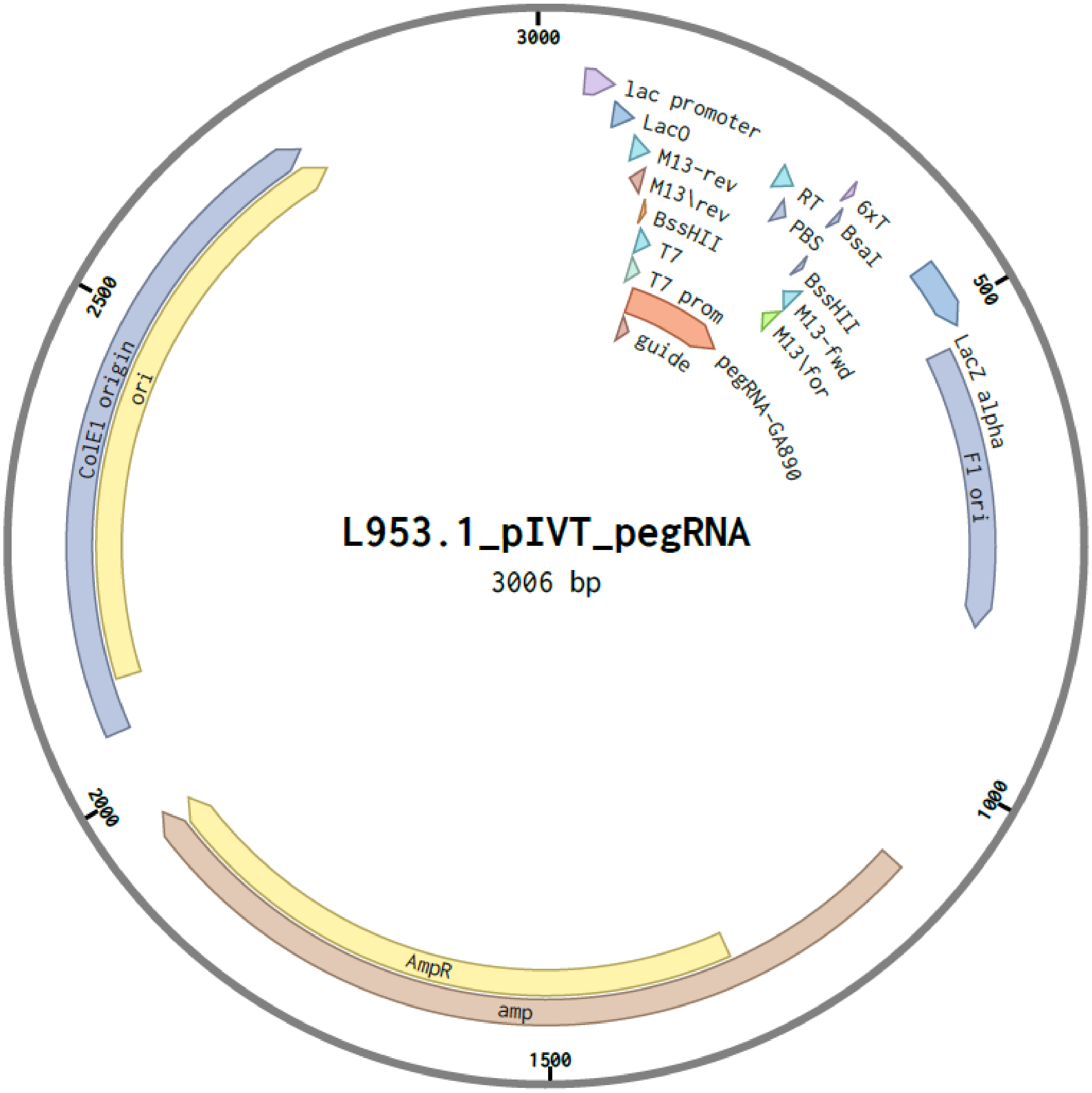


Figure S19: Plasmid used for prime editing experiments. Plasmids used to produce pegRNA via IVT.

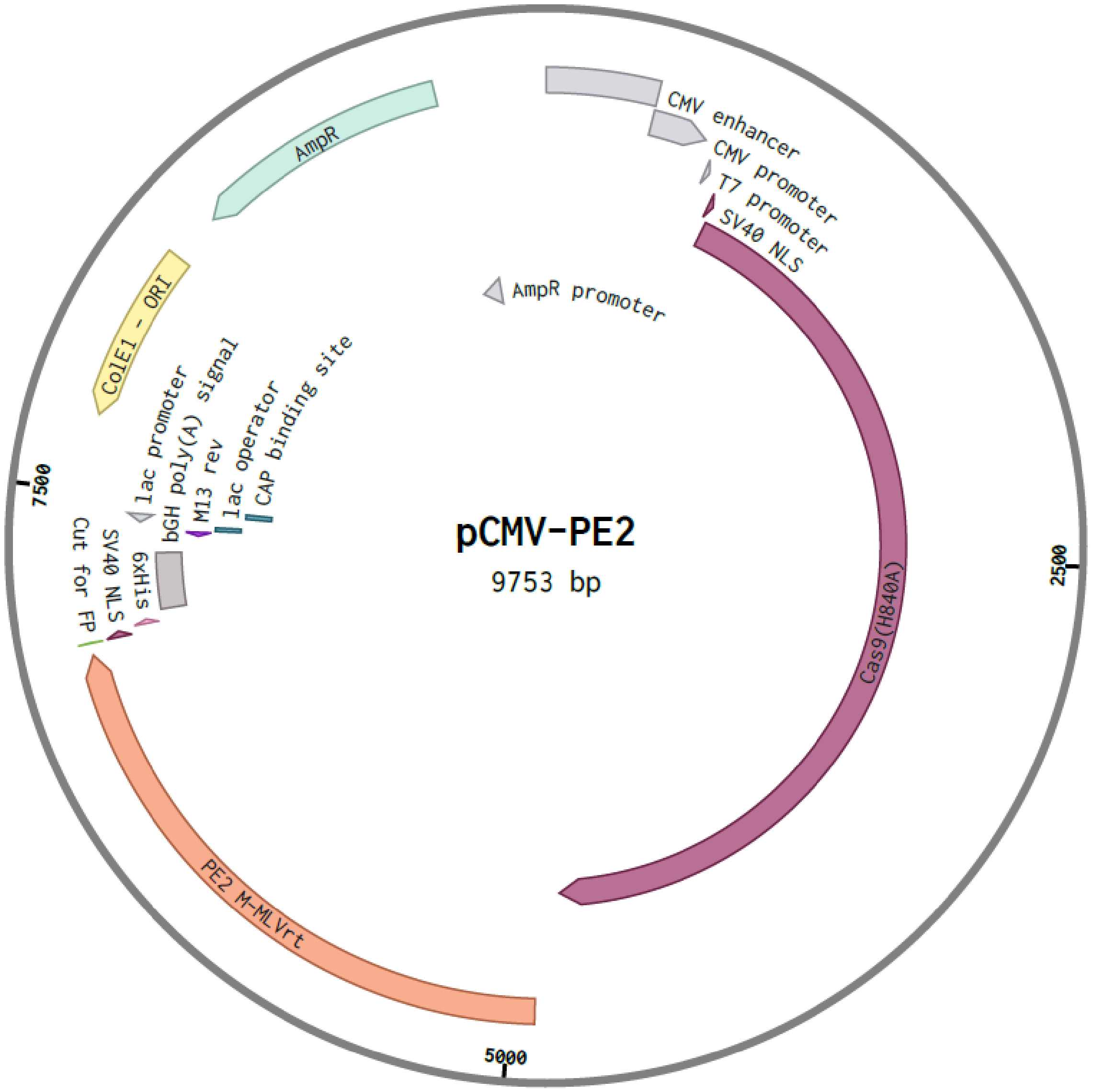


Figure S20: Plasmid used for prime editing experiments. The prime editor is expressed.

cggggggtgcgtgcgtgtgtgtgtgcgtggggagcgccgcgtgcggcccgcgctgcccggcggctgtgagcgctgcgggcgcggcg
cggggctttgtgcgctccgcgtgtgcgcgaggggagcgcggccgggggcggtgccccgcggtgcggggggggctgcgaggggaaca
aaggctgcgtgcggggtgtgtgcgtgggggggtgagcagggggtgtgggcgcggcggtcgggctgtaacccccccctgcacccccc
tccccgagttgctgagcacggcccggcttcgggtgcggggctccgtgcggggcgtggcgcggggctcgccgtgccgggcggggggt
ggcggcaggtgggggtgccgggcggggcggggccgcctcgggccggggagggctcgggggaggggcgcggcggccccggagcg
ccggcggctgtcgaggcgcggcgagccgcagccattgccttttatggtaatcgtgcgagagggcgcagggacttcctttgtcccaaat
ctggcggagccgaaatctgggaggcgccgccgcaccccctctagcgggcgcgggcgaagcggtgcggcgccggcaggaaggaaat
gggcggggagggccttcgtgcgtcgccgcgccgccgtccccttctccatctccagcctcggggctgccgcagggggacggctgccttc
gggggggacggggcagggcggggttcggcttctggcgtgtgaccggcggctctagagcctctgctaaccatgttcatgccttcttctt
ttcctacagatccttaattaataatacgactcactataaggaatacaagctacttgttctttttgcattgtacaactcactatttgttttcgc
gcccagttgcaaaaagtgtcgccaccatgaccctgaacatcgaggacgagtaccggctgcacgagacaagcaaagaacccgatgtgt
ccctgggcagcacctggcttagtgatttccctcaggcctgggccgagacaggcggaatgggacttgctgttagacaggcccctctgat
catccctctgaaggccacaagcacccctgtgtccatcaagcagtaccccatgagccaagaggcccggctgggaatcaagccccacatt
cagagactgctggaccagggcatcctggtgccttgtcagagcccttggaatacccctctgctgcccgtgaagaagcccggcaccaacg
attacagacccgtgcaggacctgcgggaagtgaacaagagagtggaagatattcaccccaccgtgccgaatccttacaacctgctgtc
tggcctgcctcctagccaccagtggtacacagtgctggacctgaaggacgccttcttctgtctgcggctgcaccctacaagccagcctc
tgtttgcctttgagtggcgggaccctgagatgggcattagcggacagctgacctggaccagactgccccagggcttcaagaacagccc
cacactgttcaacgaggccctgcatagggacctcgccgacttcagaatccagcatcctgacctgatcctgctccagtacgtggacgatc
tgctgctggccgctacaagcgagctggattgtcagcagggaacaagagccctgctgcaaaccctgggcaacctgggctatagagcct
ctgccaagaaggcccagatttgccagaaacaagtgaagtatctgggctacctgctgaaagagggccagcgttggctgaccgaggcca
gaaaagaaaccgtgatgggccagcctacacctaagacacccagacagctgagagagttcctgggcaaagccggattctgtcggctgt
tcatccctggctttgccgagatggctgcccctctgtacccactgacaaagcccggaactctgttcaactggggcccagatcagcagaag
gcctaccaagagatcaagcaggctctgctgacagcccctgctctgggactgcctgatctgaccaagcctttcgagctgttcgtggacga
gaagcagggctatgccaaaggcgtgctgacacagaagctcggcccttggagaaggcctgtggcctacctgagcaagaaactggacc
ctgtggctgccggatggcctccttgtctgagaatggtggccgccattgccgtgctgaccaaggatgccggaaagctgacaatgggaca
gcctctggtcattctggcccctcatgccgtggaagccctcgtgaaacagcctcctgatcggtggctgagcaacgccaggatgacacac
tatcaggcactgctgctcgacaccgacagagtgcagtttggacctgtggtggccctgaatcctgccacacttctgcctctgcctgagga

aggcctccagcacaattgcctggacatcctggccgaggctcacggcacaagacccgatctgacagatcagccactgcctgacgccga
ccacacctggtatacagatggcagctctctgctgcaagaaggacagagaaaagccggggctgccgtgaccaccgagacagaagtga
tttgggccaaagctctgcccgctggcacatctgctcagagagccgaactgatcgccctgacacaggccctgaaaatggccgagggca
agaagctgaacgtctacaccgactccagatacgccttcgccaccgctcacatccacggcgaaatctatcggcggagaggatggctgac
cagcgagggcaaagagattaagaacaaggacgagattctcgccctgctcaaggccctgttcctgcctaagcggctgagcatcatccac
tgtccaggccaccagaagggccactctgctgaagctagaggcaacagaatggccgaccaggctgccagaaaggccgccatcacaga
gacacccgataccagcacactgctgatcgagaacagcagccctggatccacgccgcccaagaagaagagaaaggttgaggacggcg
agggcaccggtggacctggaagcgagcctgctacatctggctctgagacacctggcacctccgagtctgctacacctgaatctggacc
tggcggatccggagacaagaagtacagcatcggcctggacatcggcaccaactctgtgggctgggccgtgatcaccgacgagtacaa
ggtgcccagcaagaaattcaaggtgctgggcaacaccgaccggcacagcatcaagaagaacctgatcggagccctgctgttcgacag
cggcgaaacagccgaggccacccggctgaagagaaccgccagaagaagatacaccagacggaagaaccggatctgctatctgcaa
gagatcttcagcaacgagatggccaaggtggacgacagcttcttccacagactggaagagtccttcctggtggaagaggataagaag
cacgagcggcaccccatcttcggcaacatcgtggacgaggtggcctaccacgagaagtaccccaccatctaccacctgagaaagaaa
ctggtggacagcaccgacaaggccgacctgcggctgatctatctggccctggcccacatgatcaagttccggggccacttcctgatcg
agggcgacctgaaccccgacaacagcgacgtggacaagctgttcatccagctggtgcagacctacaaccagctgttcgaggaaaacc
ccatcaacgccagcggcgtggacgccaaggccatcctgtctgccagactgagcaagagcagacggctggaaaatctgatcgcccagc
tgcccggcgagaagaagaatggcctgttcggcaacctgattgccctgagcctgggcctgacccccaacttcaagagcaacttcgacct
ggccgaggatgccaaactgcagctgagcaaggacacctacgacgacgacctggacaacctgctggcccagatcggcgaccagtacg
ccgacctgtttctggccgccaagaacctgtccgacgccatcctgctgagcgacatcctgagagtgaacaccgagatcaccaaggcccc
cctgagcgcctctatgatcaagagatacgacgagcaccaccaggacctgaccctgctgaaagctctcgtgcggcagcagctgcctga
gaagtacaaagagattttcttcgaccagagcaagaacggctacgccggctacattgacggcggagccagccaggaagagttctacaa
gttcatcaagcccatcctggaaaagatggacggcaccgaggaactgctcgtgaagctgaacagagaggacctgctgcggaagcagc
ggaccttcgacaacggcagcatcccccaccagatccacctgggagagctgcacgccattctgcggcggcaggaagatttttacccatt
cctgaaggacaaccgggaaaagatcgagaagatcctgaccttccgcatcccctactacgtgggccctctggccaggggaaacagcag
attcgcctggatgaccagaaagagcgaggaaaccatcaccccctggaacttcgaggaagtggtggacaagggcgcttccgcccaga
gcttcatcgagcggatgaccaacttcgataagaacctgcccaacgagaaggtgctgcccaagcacagcctgctgtacgagtacttcac
cgtgtataacgagctgaccaaagtgaaatacgtgaccgagggaatgagaaagcccgccttcctgagcggcgagcagaaaaaggcca

tcgtggacctgctgttcaagaccaaccggaaagtgaccgtgaagcagctgaaagaggactacttcaagaaaatcgagtgcttcgactc
cgtggaaatctccggcgtggaagatcggttcaacgcctccctgggcacataccacgatctgctgaaaattatcaaggacaaggacttc
ctggacaatgaggaaaacgaggacattctggaagatatcgtgctgaccctgacactgtttgaggacagagagatgatcgaggaacgg
ctgaaaacctatgcccacctgttcgacgacaaagtgatgaagcagctgaagcggcggagatacaccggctggggcaggctgagccg
gaagctgatcaacggcatccgggacaagcagtccggcaagacaatcctggatttcctgaagtccgacggcttcgccaacagaaacttc
atgcagctgatccacgacgacagcctgacctttaaagaggacatccagaaagcccaggtgtccggccagggcgatagcctgcacgag
cacattgccaatctggccggcagccccgccattaagaagggcatcctgcagacagtgaaggtggtggacgagctcgtgaaagtgatg
ggccggcacaagcccgagaacatcgtgatcgaaatggccagagagaaccagaccacccagaagggacagaagaacagccgcgaga
gaatgaagcggatcgaagagggcatcaaagagctgggcagccagatcctgaaagaacaccccgtggaaaacacccagctgcagaa
cgagaagctgtacctgtactacctgcagaatgggcgggatatgtacgtggaccaggaactggacatcaaccggctgtccgactacga
tgtggacgccatcgtgcctcagagctttctgaaggacgactccatcgacaacaaggtgctgaccagaagcgacaagaaccggggcaa
gagcgacaacgtgccctccgaagaggtcgtgaagaagatgaagaactactggcggcagctgctgaacgccaagctgattacccaga
gaaagttcgacaatctgaccaaggccgagagaggcggcctgagcgaactggataaggccggcttcatcaagagacagctggtggaa
acccggcagatcacaaagcacgtggcacagatcctggactcccggatgaacactaagtacgacgagaatgacaagctgatccggga
agtgaaagtgatcaccctgaagtccaagctggtgtccgatttccggaaggatttccagttttacaaagtgcgcgagatcaacaactacc
accacgcccacgacgcctacctgaacgccgtcgtgggaaccgccctgatcaaaaagtaccctaagctggaaagcgagttcgtgtacg
gcgactacaaggtgtacgacgtgcggaagatgatcgccaagagcgagcaggaaatcggcaaggctaccgccaagtacttcttctaca
gcaacatcatgaactttttcaagaccgagattaccctggccaacggcgagatccggaagcggcctctgatcgagacaaacggcgaaa
ccggggagatcgtgtgggataagggccgggattttgccaccgtgcggaaagtgctgagcatgccccaagtgaatatcgtgaaaaaga
ccgaggtgcagacaggcggcttcagcaaagagtctatcctgcccaagaggaacagcgataagctgatcgccagaaagaaggactgg
gaccctaagaagtacggcggcttcgacagccccaccgtggcctattctgtgctggtggtggccaaagtggaaaagggcaagtccaag
aaactgaagagtgtgaaagagctgctggggatcaccatcatggaaagaagcagcttcgagaagaatcccatcgactttctggaagcc
aagggctacaaagaagtgaaaaaggacctgatcatcaagctgcctaagtactccctgttcgagctggaaaacggccggaagagaatg
ctggcctctgccggcgaactgcagaagggaaacgaactggccctgccctccaaatatgtgaacttcctgtacctggccagccactatg
agaagctgaagggctcccccgaggataatgagcagaaacagctgtttgtggaacagcacaagcactacctggacgagatcatcgagc
agatcagcgagttctccaagagagtgatcctggccgacgctaatctggacaaagtgctgtccgcctacaacaagcaccgggataagc
ccatcagagagcaggccgagaatatcatccacctgtttaccctgaccaatctgggagcccctgccgccttcaagtactttgacaccacc

atcgaccggaagaggtacaccagcaccaaagaggtgctggacgccaccctgatccaccagagcatcaccggcctgtacgagacacg
gatcgacctgtctcagctgggaggcgacggatccacaccacctaagaagaaacggaaggtcgaggacggcgagggccctgctgcta
agagagtgaaactggactccggagctgctccagccgccaagaagaagaagctcgactacaaggacgacgacgataagtgaacgcgt
aaatgattgcagatccactagttctagagccaagcacgcagcaatgcagctcaaaacgcttagcctagccacacccccacgggaaac
agcagtgattaacctttagcaataaacgaaagtttaactaagctatactaaccccagggttggtcaatttcgtgccagccacaccctgg
tactgcatgcacgcaatgctagctgcccctttcccgtcctgggtaccccgagtctcccccgacctcgggtcccaggtatgctcccacctc
cacctgccccactcaccacctctgctagttccagacacctccatcgatggcgcgtcttaaataaaaaaaaaaaaaaaaaaaaaaaa
aaaaaaaaaaaaaaaaaaaaaaaaaaaaaaaaaaaaaaagcgatcgcgggcggcctctagaggcgcgccgatatcggccgcca
ctggccgtcgttttacaacgtcgtgactgggaaaaccctggcgttacccaacttaatcgccttgcagcacatccccctttcgccagctgg
cgtaatagcgaagaggcccgcaccgatcgcccttcccaacagttgcgcagcctgaatggcgaatgggacgcgccctgtagcggcgca
ttaagcgcggcgggtgtggtggttacgcgcagcgtgaccgctacacttgccagcgccctagcgcccgctcctttcgctttcttcccttcc
tttctcgccacgttcgccggctttccccgtcaagctctaaatcgggggctccctttagggttccgatttagtgctttacggcacctcgacc
ccaaaaaacttgattagggtgatggttcacgtagtgggccatcgccctgatagacggtttttcgccctttgacgttggagtccacgttct
ttaatagtggactcttgttccaaactggaacaacactcaaccctatctcggtctattcttttgatttataagggattttgccgatttcggcc
tattggttaaaaaatgagctgatttaacaaaaatttaacgcgaattttaacaaaatattaacgcttacaatttaggtggcacttttcggg
gaaatgtgcgcggaacccctatttgtttatttttctaaatacattcaaatatgtatccgctcatgagacaataaccctgataaatgcttca
ataatattgaaaaaggaagagtatgagtattcaacatttccgtgtcgcccttattccctttttttgcggcattttgccttcctgtttttgctca
cccagaaacgctggtgaaagtaaaagatgctgaagatcagttgggtgcacgagtgggttacatcgaactggatctcaacagcggtaa
gatccttgagagttttcgccccgaagaacgttttccaatgatgagcacttttaaagttctgctatgtggcgcggtattatcccgtattgac
gccgggcaagagcaactcggtcgccgcatacactattctcagaatgacttggttgagtactcaccagtcacagaaaagcatcttacgg
atggcatgacagtaagagaattatgcagtgctgccataaccatgagtgataacactgcggccaacttacttctgacaacgatcggagg
accgaaggagctaaccgcttttttgcacaacatgggggatcatgtaactcgccttgatcgttgggaaccggagctgaatgaagccata
ccaaacgacgagcgtgacaccacgatgcctgtagcaatggcaacaacgttgcgcaaactattaactggcgaactacttactctagctt
cccggcaacaattaatagactggatggaggcggataaagttgcaggaccacttctgcgctcggcccttccggctggctggtttattgct
gataaatctggagccggtgagcgtgggtctcgcggtatcattgcagcactggggccagatggtaagccctcccgtatcgtagttatcta
cacgacggggagtcaggcaactatggatgaacgaaatagacagatcgctgagataggtgcctcactgattaagcattggtaactgtca
gaccaagtttactcatatatactttagattgatttaaaacttcatttttaatttaaaaggatctaggtgaagatcctttttgataatctcatg

accaaaatcccttaacgtgagttttcgttccactgagcgtcagaccccgtagaaaagatcaaaggatcttcttgagatcctttttttctgc
gcgtaatctgctgcttgcaaacaaaaaaaccaccgctaccagcggtggtttgtttgccggatcaagagctaccaactctttttccgaa
ggtaactggcttcagcagagcgcagataccaaatactgttcttctagtgtagccgtagttaggccaccacttcaagaactctgtagcac
cgcctacatacctcgctctgctaatcctgttaccagtggctgctgccagtggcgataagtcgtgtcttaccgggttggactcaagacgat
agttaccggataaggcgcagcggtcgggctgaacggggggttcgtgcacacagcccagcttggagcgaacgacctacaccgaactg
agatacctacagcgtgagctatgagaaagcgccacgcttcccgaagggagaaaggcggacaggtatccggtaagcggcagggtcgg
aacaggagagcgcacgagggagcttccagggggaaacgcctggtatctttatagtcctgtcgggtttcgccacctctgacttgagcgt
cgatttttgtgatgctcgtcaggggggcggagcctatggaaaaacgccagcaacgcggcctttttacggttcctggccttttgctggcc
ttttgctcacatgttctttcctgcgttatcccctgattctgtggataaccgtattaccgcctttgagtgagctgataccgctcgccgcagcc
gaacgaccgagcgcagcgagtcagtgagcgaggaagcggaagagcgcccaatacgcaaaccgcctctccccgcgcgttggccgatt
cattaatgcagctggcacgacaggtttcccgactggaaagcgggcagtgagcgcaacgcaattaatgtgagttagctcactcattagg
caccccaggctttacactttatgcttccggctcgtatgttgtgtggaattgtgagcggataacaatttcacacaggaaacagctatgacc
atgaggcgcgccggattc

#### The full sequence of the plasmid encoding the prime editor and an additional mScarlet-I for observation of prime editor expression is provided below

acggcgagatccggaagcggcctctgatcgagacaaacggcgaaaccggggagatcgtgtgggataagggccgggattttgccacc
gtgcggaaagtgctgagcatgccccaagtgaatatcgtgaaaaagaccgaggtgcagacaggcggcttcagcaaagagtctatcctg
cccaagaggaacagcgataagctgatcgccagaaagaaggactgggaccctaagaagtacggcggcttcgacagccccaccgtggc
ctattctgtgctggtggtggccaaagtggaaaagggcaagtccaagaaactgaagagtgtgaaagagctgctggggatcaccatcat
ggaaagaagcagcttcgagaagaatcccatcgactttctggaagccaagggctacaaagaagtgaaaaaggacctgatcatcaagct
gcctaagtactccctgttcgagctggaaaacggccggaagagaatgctggcctctgccggcgaactgcagaagggaaacgaactggc
cctgccctccaaatatgtgaacttcctgtacctggccagccactatgagaagctgaagggctcccccgaggataatgagcagaaacag
ctgtttgtggaacagcacaagcactacctggacgagatcatcgagcagatcagcgagttctccaagagagtgatcctggccgacgcta
atctggacaaagtgctgtccgcctacaacaagcaccgggataagcccatcagagagcaggccgagaatatcatccacctgtttaccct

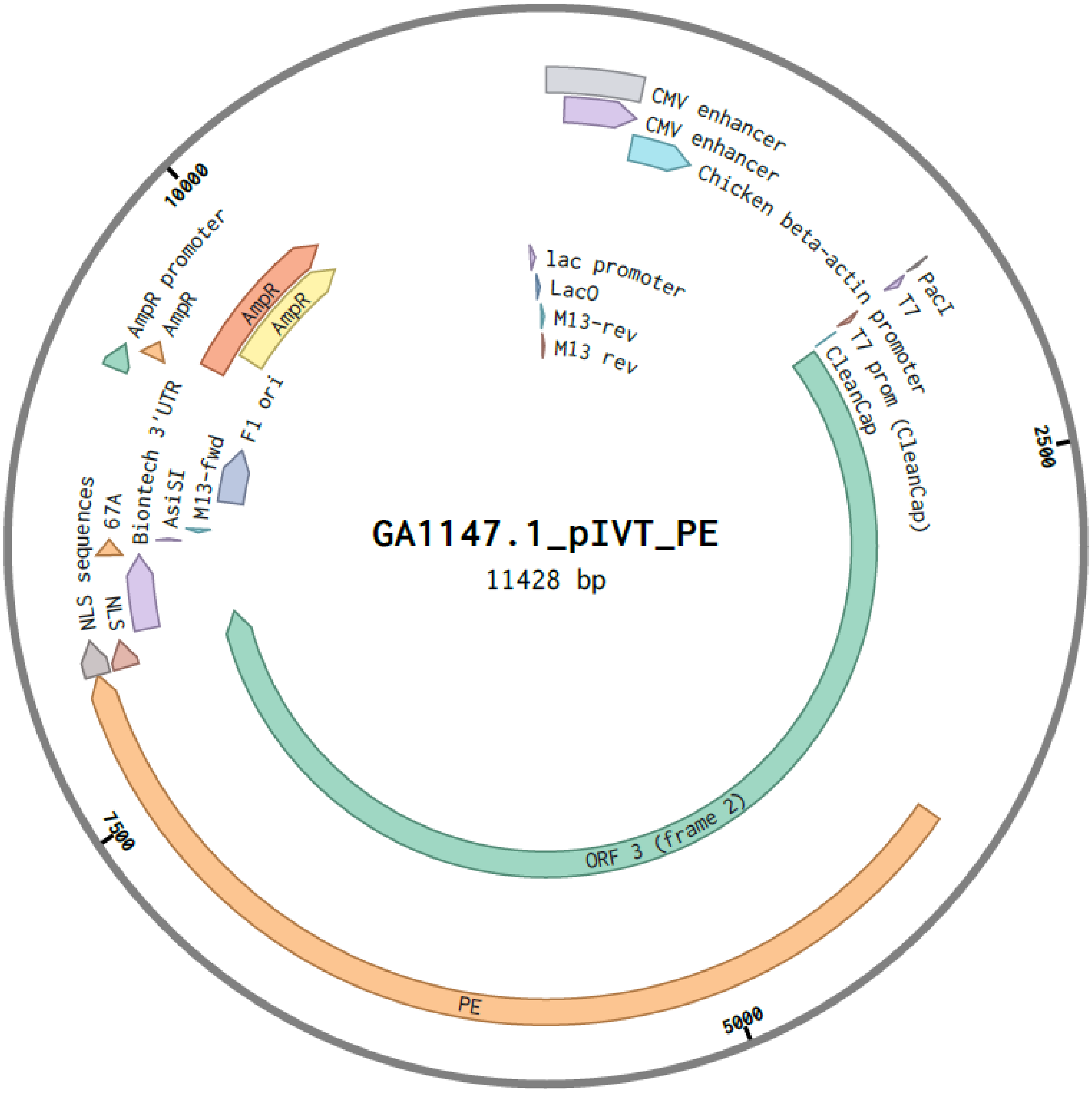


Figure S21: Plasmid used for prime editing experiments. The prime editor and mScarlet-I are expressed to visualize expression of the prime editor.

gaccaatctgggagcccctgccgccttcaagtactttgacaccaccatcgaccggaagaggtacaccagcaccaaagaggtgctgga
cgccaccctgatccaccagagcatcaccggcctgtacgagacacggatcgacctgtctcagctgggaggtgactctggaggatctagc
ggaggatcctctggcagcgagacaccaggaacaagcgagtcagcaacaccagagagcagtggcggcagcagcggcggcagcagca
ccctaaatatagaagatgagtatcggctacatgagacctcaaaagagccagatgtttctctagggtccacatggctgtctgattttcctc
aggcctgggcggaaaccgggggcatgggactggcagttcgccaagctcctctgatcatacctctgaaagcaacctctacccccgtgtc
cataaaacaataccccatgtcacaagaagccagactggggatcaagccccacatacagagactgttggaccagggaatactggtacc
ctgccagtcccccctggaacacgcccctgctacccgttaagaaaccagggactaatgattataggcctgtccaggatctgagagaagtc
aacaagcgggtggaagacatccaccccaccgtgcccaacccttacaacctcttgagcgggctcccaccgtcccaccagtggtacactg
tgcttgatttaaaggatgcctttttctgcctgagactccaccccaccagtcagcctctcttcgcctttgagtggagagatccagagatgg
gaatctcaggacaattgacctggaccagactcccacagggtttcaaaaacagtcccaccctgtttaatgaggcactgcacagagacct
agcagacttccggatccagcacccagacttgatcctgctacagtacgtggatgacttactgctggccgccacttctgagctagactgcc
aacaaggtactcgggccctgttacaaaccctagggaacctcgggtatcgggcctcggccaagaaagcccaaatttgccagaaacagg
tcaagtatctggggtatcttctaaaagagggtcagagatggctgactgaggccagaaaagagactgtgatggggcagcctactccga
agacccctcgacaactaagggagttcctagggaaggcaggcttctgtcgcctcttcatccctgggtttgcagaaatggcagccccccctg
taccctctcaccaaaccggggactctgtttaattggggcccagaccaacaaaaggcctatcaagaaatcaagcaagctcttctaactgc
cccagccctggggttgccagatttgactaagccctttgaactctttgtcgacgagaagcagggctacgccaaaggtgtcctaacgcaa
aaactgggaccttggcgtcggccggtggcctacctgtccaaaaagctagacccagtagcagctgggtggcccccttgcctacggatgg
tagcagccattgccgtactgacaaaggatgcaggcaagctaaccatgggacagccactagtcattctggcccccccatgcagtagaggc
actagtcaaacaacccccccgaccgctggctttccaacgcccggatgactcactatcaggccttgcttttggacacggaccgggtccagt
tcggaccggtggtagccctgaacccggctacgctgctcccactgcctgaggaagggctgcaacacaactgccttgatatcctggccga
agcccacggaacccgacccgacctaacggaccagccgctcccagacgccgaccacacctggtacacggatggaagcagtctcttaca
agagggacagcgtaaggcgggagctgcggtgaccaccgagaccgaggtaatctgggctaaagccctgccagccgggacatccgctc
agcgggctgaactgatagcactcacccaggccctaaagatggcagaaggtaagaagctaaatgtttatactgatagccgttatgctttt
gctactgcccatatccatggagaaatatacagaaggcgtgggtggctcacatcagaaggcaaagagatcaaaaataaagacgagatc
ttggccctactaaaagccctctttctgcccaaaagacttagcataatccattgtccaggacatcaaaagggacacagcgccgaggcta
gaggcaaccggatggctgaccaagcggcccgaaaggcagccatcacagagactccagacacctctaccctcctcatagaaaattcat
caccctctggcggctcaaaaagaaccgccgacggcagcgaattcgagcccaagaagaagaggaaagtcggaagcggagctactaac

ttcagcctgctgaagcaggctggagacgtggaggagaaccctggacctatggatagcaccgaggcagtgatcaaggagttcatgcgg
ttcaaggtgcacatggagggctccatgaacggccacgagttcgagatcgagggcgagggcgagggccgcccctacgagggcaccca
gaccgccaagctgagggtgaccaagggtggcccctgcccttctcctgggacatcctgtcccctcagttcatgtacggctccagggcct
tcacgaagcaccccgccgacatccccgactactggaagcagtccttccccgagggcttcaagtgggagcgcgtgatgaacttcgagg
acggcggcgccgtgtccgtggcccaggacacctccctggaggacggcaccctgatctacaaggtgaagctccgcggcaccaacttcc
ctcctgacggccccgtaatgcagaagaagacaatgggctgggaagcatccaccgagcggttgtaccccgaggacgtcgtgctgaagg
gcgacattaagatggccctgcgcctgaaggacggcggccgctacctggcggacttcaagaccacctacagggccaagaagcccgtgc
agatgcccggcgccttcaacatcgaccgcaagttggacatcacatcccacaacgaggactacaccgtggtggaacagtacgaacgct
ccgtggcccgccactccaccggcggctccggtggctccttgtacaagtctggtggttctcccaagaagaagaggaaagtctaaccggt
catcatcaccatcaccattgagtttaaacccgctgatcagcctcgactgtgccttctagttgccagccatctgttgtttgcccctcccccg
tgccttccttgaccctggaaggtgccactcccactgtcctttcctaataaaatgagaaattgcatcgcattgtctgagtaggtgtcattc
tattctggggggtggggtggggcaggacagcaagggggaggattgggaagacaatagcaggcatgctggggatgcggtgggctcta
tggcttctgaggcggaaagaaccagctggggctcgataccgtcgacctctagctagagcttggcgtaatcatggtcatagctgtttcct
gtgtgaaattgttatccgctcacaattccacacaacatacgagccggaagcataaagtgtaaagcctagggtgcctaatgagtgagct
aactcacattaattgcgttgcgctcactgcccgctttccagtcgggaaacctgtcgtgccagctgcattaatgaatcggccaacgcgcg
gggagaggcggtttgcgtattgggcgctcttccgcttcctcgctcactgactcgctgcgctcggtcgttcggctgcggcgagcggtatc
agctcactcaaaggcggtaatacggttatccacagaatcaggggataacgcaggaaagaacatgtgagcaaaaggccagcaaaagg
ccaggaaccgtaaaaaggccgcgttgctggcgtttttccataggctccgcccccctgacgagcatcacaaaaatcgacgctcaagtca
gaggtggcgaaacccgacaggactataaagataccaggcgtttccccctggaagctccctcgtgcgctctcctgttccgaccctgccg
cttaccggatacctgtccgcctttctcccttcgggaagcgtggcgctttctcatagctcacgctgtaggtatctcagttcggtgtaggtcg
ttcgctccaagctgggctgtgtgcacgaaccccccgttcagcccgaccgctgcgccttatccggtaactatcgtcttgagtccaacccg
gtaagacacgacttatcgccactggcagcagccactggtaacaggattagcagagcgaggtatgtaggcggtgctacagagttcttga
agtggtggcctaactacggctacactagaagaacagtatttggtatctgcgctctgctgaagccagttaccttcggaaaaagagttggt
agctcttgatccggcaaacaaaccaccgctggtagcggtggtttttttgtttgcaagcagcagattacgcgcagaaaaaaaggatctca
agaagatcctttgatcttttctacggggtctgacgctcagtggaacgaaaactcacgttaagggattttggtcatgagattatcaaaaag
gatcttcacctagatccttttaaattaaaaatgaagttttaaatcaatctaaagtatatatgagtaaacttggtctgacagttaccaatgc
ttaatcagtgaggcacctatctcagcgatctgtctatttcgttcatccatagttgcctgactccccgtcgtgtagataactacgatacggg

agggcttaccatctggccccagtgctgcaatgataccgcgagacccacgctcaccggctccagatttatcagcaataaaccagccagc
cggaagggccgagcgcagaagtggtcctgcaactttatccgcctccatccagtctattaattgttgccgggaagctagagtaagtagtt
cgccagttaatagtttgcgcaacgttgttgccattgctacaggcatcgtggtgtcacgctcgtcgtttggtatggcttcattcagctccgg
ttcccaacgatcaaggcgagttacatgatcccccatgttgtgcaaaaaagcggttagctccttcggtcctccgatcgttgtcagaagta
agttggccgcagtgttatcactcatggttatggcagcactgcataattctcttactgtcatgccatccgtaagatgcttttctgtgactgg
tgagtactcaaccaagtcattctgagaatagtgtatgcggcgaccgagttgctcttgcccggcgtcaatacgggataataccgcgccac
atagcagaactttaaaagtgctcatcattggaaaacgttcttcggggcgaaaactctcaaggatcttaccgctgttgagatccagttcg
atgtaacccactcgtgcacccaactgatcttcagcatcttttactttcaccagcgtttctgggtgagcaaaaacaggaaggcaaaatgc
cgcaaaaaagggaataagggcgacacggaaatgttgaatactcatactcttcctttttcaatattattgaagcatttatcagggttattg
tctcatgagcggatacatatttgaatgtatttagaaaaataaacaaataggggttccgcgcacatttccccgaaaagtgccacctgacg
tcgacggatcgggagatcgatctcccgatcccctagggtctactctcagtacaatctgctctgatgccgcatagttaagccagtatctgc
tccctgcttgtgtgttggaggtcgctgagtagtgcgcgagcaaaatttaagctacaacaaggcaaggcttgaccgacaattgcatgaa
gaatctgcttagggttaggcgttttgcgctgcttcgcgatgtacgggccagatatacgcgttgacattgattattgactagttattaatag
taatcaattacggggtcattagttcatagcccatatatggagttccgcgttacataacttacggtaaatggcccgcctggctgaccgccc
aacgacccccgcccattgacgtcaataatgacgtatgttcccatagtaacgccaatagggactttccattgacgtcaatgggtggagta
tttacggtaaactgcccacttggcagtacatcaagtgtatcatatgccaagtacgccccctattgacgtcaatgacggtaaatggcccg
cctggcattatgcccagtacatgaccttatgggactttcctacttggcagtacatctacgtattagtcatcgctattaccatggtgatgcg
gttttggcagtacatcaatgggcgtggatagcggtttgactcacggggatttccaagtctccaccccattgacgtcaatgggagtttgtt
ttggcaccaaaatcaacgggactttccaaaatgtcgtaacaactccgccccattgacgcaaatgggcggtaggcgtgtacggtgggag
gtctatataagcagagctggtttagtgaaccgtcagatccgctagagatccgcggccgctaatacgactcactatagggagagccgcc
accatgaaacggacagccgacggaagcgagttcgagtcaccaaagaagaagcggaaagtcgacaagaagtacagcatcggcctgg
acatcggcaccaactctgtgggctgggccgtgatcaccgacgagtacaaggtgcccagcaagaaattcaaggtgctgggcaacaccg
accggcacagcatcaagaagaacctgatcggagccctgctgttcgacagcggcgaaacagccgaggccacccggctgaagagaacc
gccagaagaagatacaccagacggaagaaccggatctgctatctgcaagagatcttcagcaacgagatggccaaggtggacgacag
cttcttccacagactggaagagtccttcctggtggaagaggataagaagcacgagcggcaccccatcttcggcaacatcgtggacgag
gtggcctaccacgagaagtaccccaccatctaccacctgagaaagaaactggtggacagcaccgacaaggccgacctgcggctgatc
tatctggccctggcccacatgatcaagttccggggccacttcctgatcgagggcgacctgaaccccgacaacagcgacgtggacaagc

tgttcatccagctggtgcagacctacaaccagctgttcgaggaaaaccccatcaacgccagcggcgtggacgccaaggccatcctgtc
tgccagactgagcaagagcagacggctggaaaatctgatcgcccagctgcccggcgagaagaagaatggcctgttcggaaacctgat
tgccctgagcctgggcctgacccccaacttcaagagcaacttcgacctggccgaggatgccaaactgcagctgagcaaggacaccta
cgacgacgacctggacaacctgctggcccagatcggcgaccagtacgccgacctgtttctggccgccaagaacctgtccgacgccatc
ctgctgagcgacatcctgagagtgaacaccgagatcaccaaggcccccctgagcgcctctatgatcaagagatacgacgagcaccac
caggacctgaccctgctgaaagctctcgtgcggcagcagctgcctgagaagtacaaagagattttcttcgaccagagcaagaacggct
acgccggctacattgacggcggagccagccaggaagagttctacaagttcatcaagcccatcctggaaaagatggacggcaccgagg
aactgctcgtgaagctgaacagagaggacctgctgcggaagcagcggaccttcgacaacggcagcatcccccaccagatccacctgg
gagagctgcacgccattctgcggcggcaggaagatttttacccattcctgaaggacaaccgggaaaagatcgagaagatcctgacctt
ccgcatcccctactacgtgggccctctggccaggggaaacagcagattcgcctggatgaccagaaagagcgaggaaaccatcacccc
ctggaacttcgaggaagtggtggacaagggcgcttccgcccagagcttcatcgagcggatgaccaacttcgataagaacctgcccaa
cgagaaggtgctgcccaagcacagcctgctgtacgagtacttcaccgtgtataacgagctgaccaaagtgaaatacgtgaccgaggg
aatgagaaagcccgccttcctgagcggcgagcagaaaaaggccatcgtggacctgctgttcaagaccaaccggaaagtgaccgtga
agcagctgaaagaggactacttcaagaaaatcgagtgcttcgactccgtggaaatctccggcgtggaagatcggttcaacgcctccct
gggcacataccacgatctgctgaaaattatcaaggacaaggacttcctggacaatgaggaaaacgaggacattctggaagatatcgt
gctgaccctgacactgtttgaggacagagagatgatcgaggaacggctgaaaacctatgcccacctgttcgacgacaaagtgatgaa
gcagctgaagcggcggagatacaccggctggggcaggctgagccggaagctgatcaacggcatccgggacaagcagtccggcaag
acaatcctggatttcctgaagtccgacggcttcgccaacagaaacttcatgcagctgatccacgacgacagcctgacctttaaagagg
acatccagaaagcccaggtgtccggccagggcgatagcctgcacgagcacattgccaatctggccggcagccccgccattaagaagg
gcatcctgcagacagtgaaggtggtggacgagctcgtgaaagtgatgggccggcacaagcccgagaacatcgtgatcgaaatggcc
agagagaaccagaccacccagaagggacagaagaacagccgcgagagaatgaagcggatcgaagagggcatcaaagagctgggc
agccagatcctgaaagaacaccccgtggaaaacacccagctgcagaacgagaagctgtacctgtactacctgcagaatgggcgggat
atgtacgtggaccaggaactggacatcaaccggctgtccgactacgatgtggacgctatcgtgcctcagagctttctgaaggacgact
ccatcgacaacaaggtgctgaccagaagcgacaagaaccggggcaagagcgacaacgtgccctccgaagaggtcgtgaagaagat
gaagaactactggcggcagctgctgaacgccaagctgattacccagagaaagttcgacaatctgaccaaggccgagagaggcggcc
tgagcgaactggataaggccggcttcatcaagagacagctggtggaaacccggcagatcacaaagcacgtggcacagatcctggact
cccggatgaacactaagtacgacgagaatgacaagctgatccgggaagtgaaagtgatcaccctgaagtccaagctggtgtccgattt

ccggaaggatttccagttttacaaagtgcgcgagatcaacaactaccaccacgcccacgacgcctacctgaacgccgtcgtgggaacc
gccctgatcaaaaagtaccctaagctggaaagcgagttcgtgtacggcgactacaaggtgtacgacgtgcggaagatgatcgccaag
agcgagcaggaaatcggcaaggctaccgccaagtacttcttctacagcaacatcatgaactttttcaagaccgagattaccctggcca

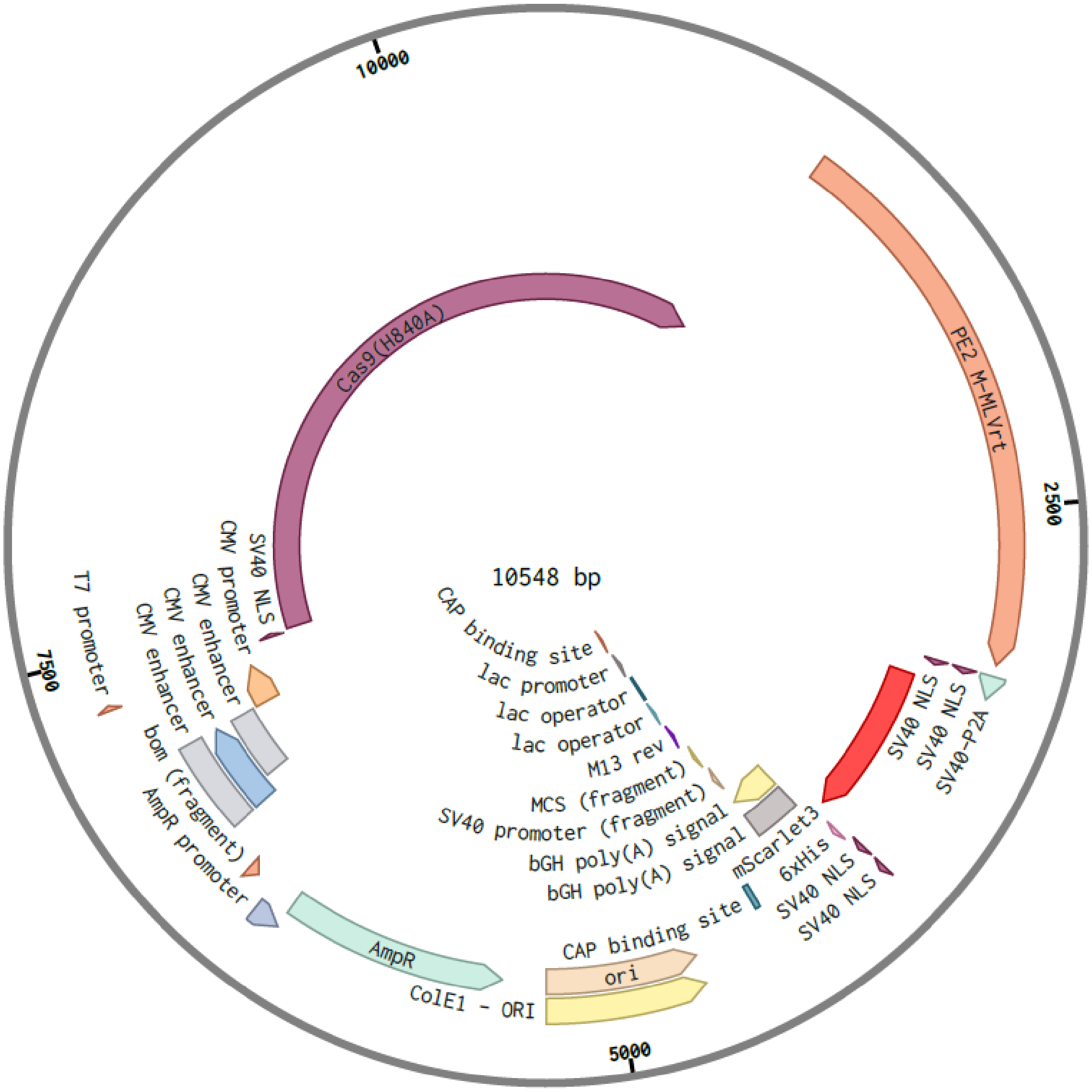


Figure S22: Plasmid used for prime editing experiments. Plasmid used to produce RNA for the prime editor via IVT.

#### The full sequence of the plasmid encoding the prime editor and an additional mScarlet-I for observation of prime editor expression for mRNA production is provided below

gacattgattattgactagttattaatagtaatcaattacggggtcattagttcatagcccatatatggagttccgcgttacataacttac
ggtaaatggcccgcctggctgaccgcccaacgacccccgcccattgacgtcaataatgacgtatgttcccatagtaacgccaataggg
actttccattgacgtcaatgggtggagtatttacggtaaactgcccacttggcagtacatcaagtgtatcatatgccaagtacgcccccct
attgacgtcaatgacggtaaatggcccgcctggcattatgcccagtacatgaccttatgggactttcctacttggcagtacatctacgta
ttagtcatcgctattaccaggtcgaggtgagccccacgttctgcttcactctccccatctcccccccctccccacccccaattttgtattta
tttatttttttaattattttgtgcagcgatgggggcggggggggggggggggcgcgcgccaggcggggcggggcggggcgaggggcg
gggcggggcgaggcggagaggtgcggcggcagccaatcagagcggcgcgctccgaaagtttccttttatggcgaggcggcggcgg
cggcggccctataaaaagcgaagcgcgcggcgggcgggagtcgctgcgcgctgccttcgccccgtgccccgctccgccgccgcctcg
cgccgcccgccccggctctgactgaccgcgttactcccacaggtgagcgggcgggacggcccttctcctccgggctgtaattagcgct
tggtttaatgacggcttgtttcttttctgtggctgcgtgaaagccttgaggggctccgggagggccctttgtgcggggggagcggctcg
gggggtgcgtgcgtgtgtgtgtgcgtggggagcgccgcgtgcggctccgcgctgcccggcggctgtgagcgctgcgggcgcggcgc
ggggctttgtgcgctccgcagtgtgcgcgaggggagcgcggccgggggcggtgccccgcggtgcggggggggctgcgaggggaac
aaaggctgcgtgcggggtgtgtgcgtgggggggtgagcagggggtgtgggcgcgtcggtcgggctgcaaccccccctgcacccccc
tccccgagttgctgagcacggcccggcttcgggtgcggggctccgtacggggcgtggcgcggggctcgccgtgccgggcggggggt
ggcggcaggtgggggtgccgggcggggcggggccgcctcgggccggggagggctcgggggaggggcgcggcggcccccggagc
gccggcggctgtcgaggcgcggcgagccgcagccattgccttttatggtaatcgtgcgagagggcgcagggacttcctttgtcccaaa
tctgtgcggagccgaaatctgggaggcgccgccgcaccccctctagcgggcgcggggcgaagcggtgcggcgccggcaggaagga
aatgggcggggagggccttcgtgcgtcgccgcgccgccgtccccttctccctctccagcctcggggctgtccgcggggggacggctgc
cttcgggggggacggggcagggcggggttcggcttctggcgtgtgaccggcggctctagagcctctgctaaccatgttcatgccttct
tctttttcctacagatccttaattaataatacgactcactataggaatacaagctacttgttctttttgcagccgccaccatgaaacgga
cagccgacggaagcgagttcgagtcaccaaagaagaagcggaaagtcgacaagaagtacagcatcggcctggacatcggcaccaa
ctctgtgggctgggccgtgatcaccgacgagtacaaggtgcccagcaagaaattcaaggtgctgggcaacaccgaccggcacagcat
caagaagaacctgatcggagccctgctgttcgacagcggcgaaacagccgaggccacccggctgaagagaaccgccagaagaaga

tacaccagacggaagaaccggatctgctatctgcaagagatcttcagcaacgagatggccaaggtggacgacagcttcttccacaga

ctggaagagtccttcctggtggaagaggataagaagcacgagcggcaccccatcttcggcaacatcgtggacgaggtggcctaccac

gagaagtaccccaccatctaccacctgagaaagaaactggtggacagcaccgacaaggccgacctgcggctgatctatctggccctg

gcccacatgatcaagttccggggccacttcctgatcgagggcgacctgaaccccgacaacagcgacgtggacaagctgttcatccag

ctggtgcagacctacaaccagctgttcgaggaaaaccccatcaacgccagcggcgtggacgccaaggccatcctgtctgccagactg

agcaagagcagacggctggaaaatctgatcgcccagctgcccggcgagaagaagaatggcctgttcggaaacctgattgccctgagc

ctgggcctgacccccaacttcaagagcaacttcgacctggccgaggatgccaaactgcagctgagcaaggacacctacgacgacgac

ctggacaacctgctggcccagatcggcgaccagtacgccgacctgtttctggccgccaagaacctgtccgacgccatcctgctgagcg

acatcctgagagtgaacaccgagatcaccaaggcccccctgagcgcctctatgatcaagagatacgacgagcaccaccaggacctga

ccctgctgaaagctctcgtgcggcagcagctgcctgagaagtacaaagagattttcttcgaccagagcaagaacggctacgccggcta

cattgacggcggagccagccaggaagagttctacaagttcatcaagcccatcctggaaaagatggacggcaccgaggaactgctcgt

gaagctgaacagagaggacctgctgcggaagcagcggaccttcgacaacggcagcatccccccaccagatccacctgggagagctgc

acgccattctgcggcggcaggaagatttttacccattcctgaaggacaaccgggaaaagatcgagaagatcctgaccttccgcatccc

ctactacgtgggccctctggccaggggaaacagcagattcgcctggatgaccagaaagagcgaggaaaccatcaccccctggaactt

cgaggaagtggtggacaagggcgcttccgcccagagcttcatcgagcggatgaccaacttcgataagaacctgcccaacgagaaggt

gctgcccaagcacagcctgctgtacgagtacttcaccgtgtataacgagctgaccaaagtgaaatacgtgaccgagggaatgagaaa

gcccgccttcctgagcggcgagcagaaaaaggccatcgtggacctgctgttcaagaccaaccggaaagtgaccgtgaagcagctgaa

agaggactacttcaagaaaatcgagtgcttcgactccgtggaaatctccggcgtggaagatcggttcaacgcctccctgggcacatac

cacgatctgctgaaaattatcaaggacaaggacttcctggacaatgaggaaaacgaggacattctggaagatatcgtgctgaccctga

cactgtttgaggacagagagatgatcgaggaacggctgaaaacctatgcccacctgttcgacgacaaagtgatgaagcagctgaagc

ggcggagatacaccggctggggcaggctgagccggaagctgatcaacggcatccgggacaagcagtccggcaagacaatcctggat

ttcctgaagtccgacggcttcgccaacagaaacttcatgcagctgatccacgacgacagcctgacctttaaagaggacatccagaaag

cccaggtgtccggccagggcgatagcctgcacgagcacattgccaatctggccggcagccccgccattaagaagggcatcctgcaga

cagtgaaggtggtggacgagctcgtgaaagtgatgggccggcacaagcccgagaacatcgtgatcgaaatggccagagagaaccag

accacccagaagggacagaagaacagccgcgagagaatgaagcggatcgaagagggcatcaaagagctgggcagccagatcctga

aagaacaccccgtggaaaacacccagctgcagaacgagaagctgtacctgtactacctgcagaatgggcgggatatgtacgtggacc

aggaactggacatcaaccggctgtccgactacgatgtggacgctatcgtgcctcagagctttctgaaggacgactccatcgacaacaa

ggtgctgaccagaagcgacaagaaccggggcaagagcgacaacgtgccctccgaagaggtcgtgaagaagatgaagaactactgg
cggcagctgctgaacgccaagctgattacccagagaaagttcgacaatctgaccaaggccgagagaggcggcctgagcgaactgga
taaggccggcttcatcaagagacagctggtggaaacccggcagatcacaaagcacgtggcacagatcctggactcccggatgaacac
taagtacgacgagaatgacaagctgatccgggaagtgaaagtgatcaccctgaagtccaagctggtgtccgatttccggaaggatttc
cagttttacaaagtgcgcgagatcaacaactaccaccacgcccacgacgcctacctgaacgccgtcgtgggaaccgccctgatcaaaa
agtaccctaagctggaaagcgagttcgtgtacggcgactacaaggtgtacgacgtgcggaagatgatcgccaagagcgagcaggaa
atcggcaaggctaccgccaagtacttcttctacagcaacatcatgaactttttcaagaccgagattaccctggccaacggcgagatccg
gaagcggcctctgatcgagacaaacggcgaaaccggggagatcgtgtgggataagggccgggattttgccaccgtgcggaaagtgc
tgagcatgccccaagtgaatatcgtgaaaaagaccgaggtgcagacaggcggcttcagcaaagagtctatcctgcccaagaggaaca
gcgataagctgatcgccagaaagaaggactgggaccctaagaagtacggcggcttcgacagccccaccgtggcctattctgtgctgg
tggtggccaaagtggaaaagggcaagtccaagaaactgaagagtgtgaaagagctgctggggatcaccatcatggaaagaagcagc
ttcgagaagaatcccatcgactttctggaagccaagggctacaaagaagtgaaaaaggacctgatcatcaagctgcctaagtactccc
tgttcgagctggaaaacggccggaagagaatgctggcctctgccggcgaactgcagaagggaaacgaactggccctgccctccaaa
tatgtgaacttcctgtacctggccagccactatgagaagctgaagggctcccccgaggataatgagcagaaacagctgtttgtggaac
agcacaagcactacctggacgagatcatcgagcagatcagcgagttctccaagagagtgatcctggccgacgctaatctggacaaag
tgctgtccgcctacaacaagcaccgggataagcccatcagagagcaggccgagaatatcatccacctgtttaccctgaccaatctggg
agcccctgccgccttcaagtactttgacaccaccatcgaccggaagaggtacaccagcaccaaagaggtgctggacgccaccctgat
ccaccagagcatcaccggcctgtacgagacacggatcgacctgtctcagctgggaggtgactctggaggatctagcggaggatcctc
tggcagcgagacaccaggaacaagcgagtcagcaacaccagagagcagtggcggcagcagcggcggcagcagcaccctaaatata
gaagatgagtatcggctacatgagacctcaaaagagccagatgtttctctagggtccacatggctgtctgattttcctcaggcctgggc
ggaaaccgggggcatgggactggcagttcgccaagctcctctgatcatacctctgaaagcaacctctacccccgtgtccataaaacaa
taccccatgtcacaagaagccagactggggatcaagccccacatacagagactgttggaccagggaatactggtaccctgccagtccc
cctggaacacgcccctgctacccgttaagaaaccagggactaatgattataggcctgtccaggatctgagagaagtcaacaagcggg
tggaagacatccaccccaccgtgcccaacccttacaacctcttgagcgggctcccaccgtcccaccagtggtacactgtgcttgattta
aaggatgcctttttctgcctgagactccaccccaccagtcagcctctcttcgcctttgagtggagagatccagagatgggaatctcagg
acaattgacctggaccagactcccacagggtttcaaaaacagtcccaccctgtttaatgaggcactgcacagagacctagcagacttc
cggatccagcacccagacttgatcctgctacagtacgtggatgacttactgctggccgccacttctgagctagactgccaacaaggtac

tcgggccctgttacaaaccctagggaacctcgggtatcgggcctcggccaagaaagcccaaatttgccagaaacaggtcaagtatctg
gggtatcttctaaaagagggtcagagatggctgactgaggccagaaaagagactgtgatggggcagcctactccgaagacccctcga
caactaagggagttcctagggaaggcaggcttctgtcgcctcttcatccctgggtttgcagaaatggcagcccccctgtaccctctcac
caaaccggggactctgtttaattggggcccagaccaacaaaaggcctatcaagaaatcaagcaagctcttctaactgccccagccctg
gggttgccagatttgactaagccctttgaactctttgtcgacgagaagcagggctacgccaaaggtgtcctaacgcaaaaactgggac
cttggcgtcggccggtggcctacctgtccaaaaagctagacccagtagcagctgggtggccccttgcctacggatggtagcagccat
tgccgtactgacaaaggatgcaggcaagctaaccatgggacagccactagtcattctggcccccatgcagtagaggcactagtcaaa
caaccccccgaccgctggctttccaacgcccggatgactcactatcaggccttgcttttggacacggaccgggtccagttcggaccggt
ggtagccctgaacccggctacgctgctcccactgcctgaggaagggctgcaacacaactgccttgatatcctggccgaagcccacgga
acccgacccgacctaacggaccagccgctcccagacgccgaccacacctggtacacggatggaagcagtctcttacaagagggacag
cgtaaggcgggagctgcggtgaccaccgagaccgaggtaatctgggctaaagccctgccagccgggacatccgctcagcgggctga
actgatagcactcacccaggccctaaagatggcagaaggtaagaagctaaatgtttatactgatagccgttatgcttttgctactgccc
atatccatggagaaatatacagaaggcgtgggtggctcacatcagaaggcaaagagatcaaaaataaagacgagatcttggccctac
taaaagccctctttctgcccaaaagacttagcataatccattgtccaggacatcaaaagggacacagcgccgaggctagaggcaaccg
gatggctgaccaagcggcccgaaaggcagccatcacagagactccagacacctctaccctcctcatagaaaattcatcaccctctggc
ggctcaaaaagaaccgccgacggcagcgaattcgagcccaagaagaagaggaaagtcggaagcggagctactaacttcagcctgct
gaagcaggctggagacgtggaggagaaccctggacctatggatagcaccgaggcagtgatcaaggagttcatgcggttcaaggtgc
acatggagggctccatgaacggccacgagttcgagatcgagggcgagggcgagggccgcccctacgagggcacccagaccgccaa
gctgagggtgaccaagggtggcccctgcccttctcctgggacatcctgtcccctcagttcatgtacggctccagggccttcacgaagc
accccgccgacatccccgactactggaagcagtccttccccgagggcttcaagtgggagcgcgtgatgaacttcgaggacggcggcg
ccgtgtccgtggcccaggacacctccctggaggacggcaccctgatctacaaggtgaagctccgcggcaccaacttccctcctgacgg
ccccgtaatgcagaagaagacaatgggctgggaagcatccaccgagcggttgtaccccgaggacgtcgtgctgaagggcgacattaa
gatggccctgcgcctgaaggacggcggccgctacctggcggacttcaagaccacctacagggccaagaagcccgtgcagatgcccg
gcgccttcaacatcgaccgcaagttggacatcacatcccacaacgaggactacaccgtggtggaacagtacgaacgctccgtggccc
gccactccaccggcggctccggtggctccttgtacaagtctggtcctgctgctaagagagtgaaactggactgaacgcgtaaatgatt
gcagatccactagttctagagccaagcacgcagcaatgcagctcaaaacgcttagcctagccacacccccacgggaaacagcagtga
ttaacctttagcaataaacgaaagtttaactaagctatactaaccccagggttggtcaatttcgtgccagccacaccctggtactgcatg

cacgcaatgctagctgccccttttcccgtcctgggtaccccgagtctcccccgacctcgggtcccaggtatgctcccacctccacctgccc
cactcaccacctctgctagttccagacacctccatcgatggcgcgtcttaaataaaaaaaaaaaaaaaaaaaaaaaaaaaaaaaaaaaa
aaaaaaaaaaaaaaaaaaaaaaaaaaaaaaaaagcgatcgcgggcggcctctagaggcgcgccgatatcggccgccactggccgt
cgttttacaacgtcgtgactgggaaaaccctggcgttacccaacttaatcgccttgcagcacatccccctttcgccagctggcgtaatag
cgaagaggcccgcaccgatcgcccttcccaacagttgcgcagcctgaatggcgaatgggacgcgccctgtagcggcgcattaagcgc
ggcgggtgtggtggttacgcgcagcgtgaccgctacacttgccagcgccctagcgcccgctcctttcgctttcttcccttcctttctcgcc
acgttcgccggctttccccgtcaagctctaaatcgggggctccctttagggttccgatttagtgctttacggcacctcgaccccaaaaaa
cttgattagggtgatggttcacgtagtgggccatcgccctgatagacggtttttcgccctttgacgttggagtccacgttctttaatagtg
gactcttgttccaaactggaacaacactcaaccctatctcggtctattcttttgatttataagggattttgccgatttcggcctattggtta
aaaaatgagctgatttaacaaaaatttaacgcgaattttaacaaaatattaacgcttacaatttaggtggcacttttcggggaaatgtgc
gcggaacccctatttgtttatttttctaaatacattcaaatatgtatccgctcatgagacaataaccctgataaatgcttcaataatattga
aaaaggaagagtatgagtattcaacatttccgtgtcgcccttattcccttttttgcggcattttgccttcctgtttttgctcacccagaaac
gctggtgaaagtaaaagatgctgaagatcagttgggtgcacgagtgggttacatcgaactggatctcaacagcggtaagatccttgag
agttttcgccccgaagaacgttttccaatgatgagcacttttaaagttctgctatgtggcgcggtattatcccgtattgacgccgggcaa
gagcaactcggtcgccgcatacactattctcagaatgacttggttgagtactcaccagtcacagaaaagcatcttacggatggcatgac
agtaagagaattatgcagtgctgccataaccatgagtgataacactgcggccaacttacttctgacaacgatcggaggaccgaaggag
ctaaccgcttttttgcacaacatgggggatcatgtaactcgccttgatcgttgggaaccggagctgaatgaagccataccaaacgacga
gcgtgacaccacgatgcctgtagcaatggcaacaacgttgcgcaaactattaactggcgaactacttactctagcttcccggcaacaat
taatagactggatggaggcggataaagttgcaggaccacttctgcgctcggcccttccggctggctggtttattgctgataaatctgga
gccggtgagcgtgggtctcgcggtatcattgcagcactggggccagatggtaagccctcccgtatcgtagttatctacacgacgggga
gtcaggcaactatggatgaacgaaatagacagatcgctgagataggtgcctcactgattaagcattggtaactgtcagaccaagttta
ctcatatatactttagattgatttaaaacttcatttttaatttaaaaggatctaggtgaagatcctttttgataatctcatgaccaaaatcc
cttaacgtgagttttcgttccactgagcgtcagaccccgtagaaaagatcaaaggatcttcttgagatcctttttttctgcgcgtaatctg
ctgcttgcaaacaaaaaaaccaccgctaccagcggtggtttgtttgccggatcaagagctaccaactctttttccgaaggtaactggc
ttcagcagagcgcagataccaaatactgttcttctagtgtagccgtagttaggccaccacttcaagaactctgtagcaccgcctacatac
ctcgctctgctaatcctgttaccagtggctgctgccagtggcgataagtcgtgtcttaccgggttggactcaagacgatagttaccggat
aaggcgcagcggtcgggctgaacggggggttcgtgcacacagcccagcttggagcgaacgacctacaccgaactgagatacctaca

gcgtgagctatgagaaagcgccacgcttcccgaagggagaaaggcggacaggtatccggtaagcggcagggtcggaacaggagag
cgcacgagggagcttccagggggaaacgcctggtatctttatagtcctgtcgggtttcgccacctctgacttgagcgtcgatttttgtga
tgctcgtcaggggggcggagcctatggaaaaacgccagcaacgcggcctttttacggttcctggccttttgctggccttttgctcacat
gttctttcctgcgttatcccctgattctgtggataaccgtattaccgcctttgagtgagctgataccgctcgccgcagccgaacgaccga
gcgcagcgagtcagtgagcgaggaagcggaagagcgcccaatacgcaaaccgcctctccccgcgcgttggccgattcattaatgcag
ctggcacgacaggtttcccgactggaaagcgggcagtgagcgcaacgcaattaatgtgagttagctcactcattaggcaccccaggct
ttacactttatgcttccggctcgtatgttgtgtggaattgtgagcggataacaatttcacacaggaaacagctatgaccatgaggcgcg
ccggattc

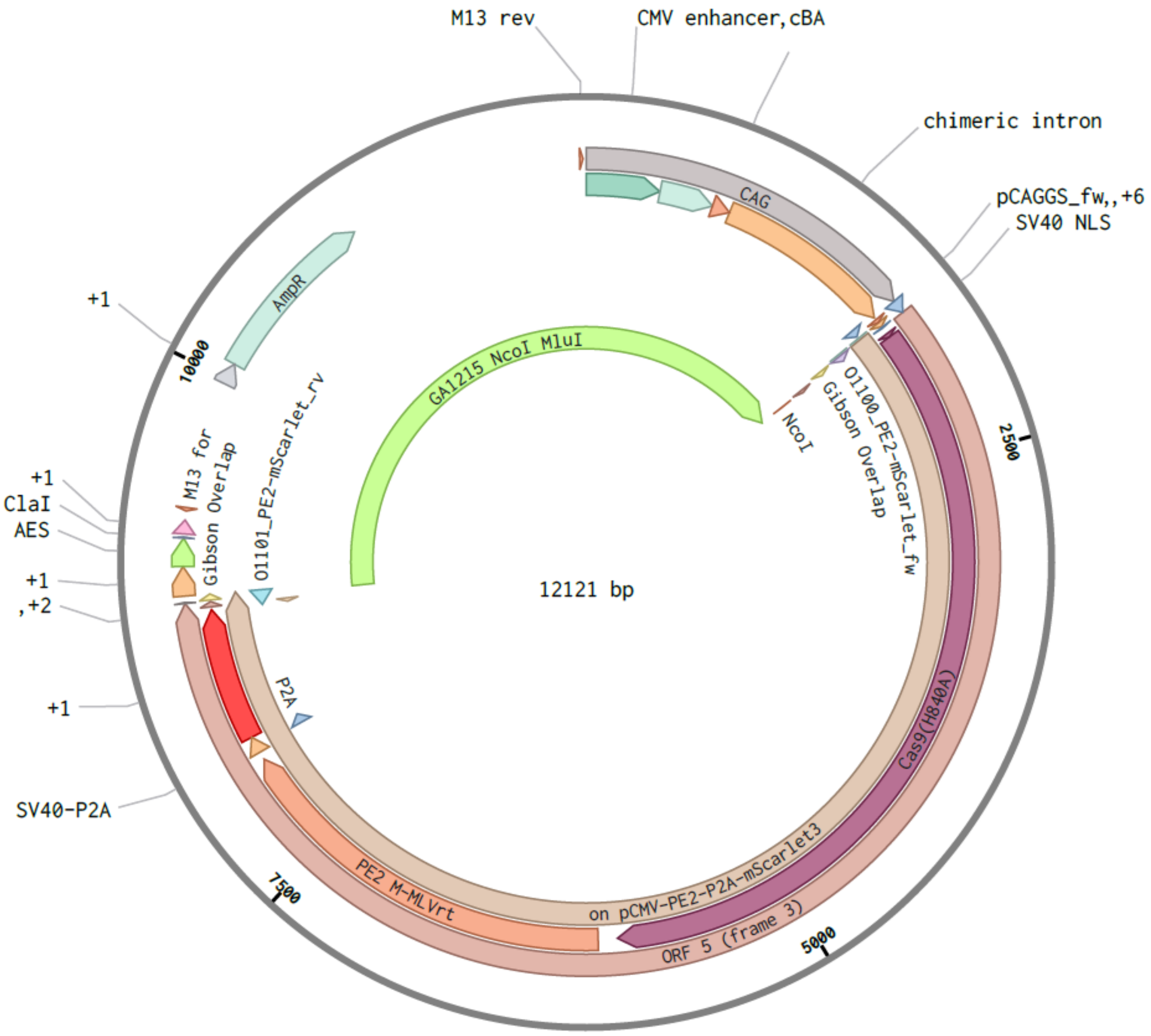


Figure S23: Plasmid used for prime editing experiments. Plasmid used to produce RNA for the prime editor and mScarlet-I for visualization via IVT.
